# Metagenomic screening for lipolytic genes reveals an ecology-clustered distribution pattern

**DOI:** 10.1101/2021.02.01.429111

**Authors:** Mingji Lu, Dominik Schneider, Rolf Daniel

## Abstract

Lipolytic enzymes are one of the most important enzyme types for application in various industrial processes. Despite the continuously increasing demand, only a small portion of the so far encountered lipolytic enzymes exhibit adequate stability and activities for biotechnological applications. To explore novel and/or extremophilic lipolytic enzymes, microbial consortia in two composts at thermophilic stage were analyzed using function-driven and sequence-based metagenomic approaches. Analysis of community composition by amplicon-based 16S rRNA genes and transcripts, and direct metagenome sequencing revealed that the communities of the compost samples were dominated by members of the phyla *Actinobacteria, Proteobacteria, Firmicutes, Bacteroidetes* and *Chloroflexi.* Function-driven screening of the metagenomic libraries constructed from the two samples yielded 115 unique genes encoding lipolytic enzymes. The family assignment of these enzymes was conducted by analyzing the phylogenetic relationship and generation of a protein sequence similarity network according to an integral classification system. The sequence-based screening was performed by using a newly developed database, containing a set of profile Hidden Markov models, highly sensitive and specific for detection of lipolytic enzymes. By comparing the lipolytic enzymes identified through both approaches, we demonstrated that the activity-directed complements sequence-based detection, and vice versa. The sequence-based comparative analysis of lipolytic genes regarding diversity, function and taxonomic origin derived from 175 metagenomes indicated significant differences between habitats. Analysis of the prevalent and distinct microbial groups providing the lipolytic genes revealed characteristic patterns and groups driven by ecological factors. The here presented data suggests that the diversity and distribution of lipolytic genes in metagenomes of various habitats are largely constrained by ecological factors.

## Introduction

Lipolytic enzymes (LEs) acting on carboxyl ester bonds in lipids, include esterases (EC 3.1.1.3, carboxylesterases) and true lipases (EC 3.1.1.1, triacylglycerol acyl hydrolases). Due to the catalytic versatility, LEs have remarkable applications in various processes relevant to food, paper, medical, detergent, and pharmaceutical industries (Hita et al. 2009; Romdhane et al. 2010; Ferrer et al. 2015; Sarmah et al. 2018). Nowadays, LEs are considered to be one of the most important biocatalysts for biotechnological applications.

In principle, LEs can be classified on the basis of the substrate preference (Sarmah et al. 2018) and sequence similarity (Chen et al. 2016). The latter provides an easy-to-perform way for classification and indication of the similarity and evolutionary relationship between LEs. Arpigny and Jaeger (1999) have elaborated the most widely accepted classification of bacterial lipases into eight families (I to VIII). The classification system was based on conserved sequence motifs and biological properties of 53 LEs. A recent update of this system resulted in addition of 11 families (IX to XIX) (Kovacic et al. 2019). Besides the nineteen families, there are claims of novel families, such as Est22 (Li et al. 2017), Est9X(Jeon et al. 2009), LipSM54 (Li et al. 2016) and EstDZ2 (Zarafeta et al. 2016). To avoid an artificial inflation of the number of families, these novel families were mostly excluded during classification of newly identified lipolytic enzymes in previous studies (Hitch and Clavel 2019). To the best of our knowledge, we aimed to incorporate all the ‘so-called’ novel families during classification to avoid false ‘novelty’.

LEs are ubiquitous among all aspects of life, with most of them originating from microorganisms (Kovacic et al. 2019). Environmental microbes, including the so far uncultured species, encode a largely untapped reservoir of novel LEs. Metagenomic function-driven and sequence-based approaches provided access to the genetic resources from so far uncultured and uncharacterized microorganisms (Simon and Daniel 2009; Simon and Daniel 2011). LEs are among the most frequent targets in function-based screens of metagenomic libraries derived from diverse habitats, such as compost (Kang et al. 2011; Lu et al. 2019), landfill leachate (Rashamuse et al. 2009), marine sediment (Peng et al. 2011; Zhang et al. 2017), activated sludges (Liaw et al. 2010) and hot springs (López-López et al. 2015).

In contrast to the easy access to metagenome-derived sequencing data, most published metagenomic screenings for LEs were enzyme activity-driven and not sequence-based (Ferrer et al. 2015; Berini et al. 2017). Only a few studies explored LEs by sequence-driven approaches, including analysis based on regular expression patterns (Masuch et al. 2015), ancestral sequence reconstruction (Verma et al. 2019) and conserved motifs (Zhang et al. 2009; Barriuso and Jesús Martínez 2015; Zarafeta et al. 2016). For various reasons, only a very limited number of LEs were identified by these strategies. Sequence-based approaches primarily rely on the reference database to infer functions of newly-discovered genes and the corresponding enzymes (Hugenholtz and Tyson 2008; Quince et al. 2017; Berini et al. 2017; Ngara and Zhang 2018). With protein-of-interest-specific databases, biomolecules such as antibiotic resistance genes (Gibson et al. 2015; Willmann et al. 2015; Pehrsson et al. 2016) and CAZymes (Wang et al. 2016; Montella et al. 2017; Wang et al. 2019) were successfully profiled across habitats. Recently, with the rapid accumulation of genomic/metagenomic data in public repositories (Keegan et al. 2016; Chen et al. 2017; Eric Sayers et al. 2019), it is likely that current knowledge regarding LEs only reflects the tip of the iceberg and that the full diversity of these enzymes is far from being completely described.

In order to quantitatively analyze LEs distributed in environmental samples, we developed a LE-specific profile Hidden Markov Model (HMM) database. Profile HMMs have been widely adopted for detection of remote homologs (Gibson et al. 2015; Walsh et al. 2017; Berglund et al. 2017) and annotation of general functions in microbial genomes and metagenomes (Skewes-Cox et al. 2014; Reyes et al. 2017; Bzhalava et al. 2018). However, they have not yet been specifically applied to LEs. Once developed and validated, the database was applied to profile the lipolytic genes in metagenomes from various habitats. Profiling the distribution of LEs among various habitats provides researchers a straightforward approach for their downstream analysis. In this study, two composts were sampled and LEs identified through function-based and sequence-based approaches were compared. The distribution of lipolytic genes in 175 metagenomes was also investigated by sequence-based screening.

## Material and Methods

### Sample collection

Compost samples were collected as described previously (Lu et al. 2019). Briefly, two compost piles fermenting mainly wood chips (Pile_1) or kitchen waste (Pile_2) were sampled. Temperatures at the sampling spots were measured, and the two samples were designated as compst55 (55 °C for Pile_1) and compst76 (76°C for Pile_2). Approximately 50 g compost per sample was collected in sterile plastic tubes and stored at −20 or −80 °C until further use.

### Isolation of nucleic acids

Metagenomic DNA of the compost sample was isolated by using the phenol-chloroform method (Zhou et al. 1996) and MoBio Power Soil DNA extraction kit as recommended by the manufacturer (MO BIO Laboratories, Hilden, Germany). DNA obtained from these two methods was pooled per sample and stored at −20 °C until use.

RNA was extracted by employing the MoBio PowerSoil RNA isolation kit as recommended by the manufacturer (MO BIO Laboratories). Residual DNA was removed by treatment with 2 U Turbo DNase (Applied Biosystems, Darmstadt, Germany) at 37 °C for 1 h and recovered by using RNeasy MinElute Cleanup kit as recommended by the manufacturer (Qiagen, Hilden, Germany). RNA yields were estimated by employing a Qubit® Fluorometer as recommended by the manufacturer (Thermo Fisher Scientific, Schwerte, Germany). A PCR reaction targeting the 16S rRNA gene was performed to verify the complete removal of DNA as described by Schneider et al. (2015). Subsequently, the DNA-free RNA was converted to cDNA using the SuperScript™ III reverse transcriptase (Thermo Fisher Scientific). Briefly, a mixture (14 μl) containing 100 ng of DNA-free RNA in DEPC-treated water, 2 μM of reverse primer (5’ - CCGTCAATTCMTTTGAGT-’) and 10 mM dNTP mix was incubated at 65 °C for 5 min and chilled on ice for at least 1 min. Then, 10 μl of cDNA synthesis mix including reaction buffer, 5 mM MgCl2, 0.01 M DTT, 1 μl 40U RiboLock™ RNase inhibitor (Thermo Fisher Scientific) and 200U SuperScript™ III reverse transcriptase (Thermo Fisher Scientific) was added to each RNA/primer mixture in the previous step, and incubated at 55 °C for 90 min. The reaction was terminated at 70 °C for 15 min.

### Sequencing of 16S rRNA genes and transcripts

The PCR amplification of the V3-V5 regions of bacterial 16S rRNA genes and transcripts were performed with the following set of primers comprising the Roche 454 pyrosequencing adaptors (underlined), a key (TCAG), a unique 10-bp multiplex identifier (MID), and template-specific sequence per sample: the forward primer V3for_B (5 ‘ -<u>CGTATCGCCTCCCTCGCGCCA</u>TCAG-MID-TACGGRAGGCAGCAG-3 ‘), (Liu et al. 2007), and reverse primer V5rev_B 5’-<u>CTATGCGCCTTGCCAGCCCGC</u>TCAG-MID-CCGTCAATTCMTTTGAGT-3’ (Wang and Qian 2009). The PCR reaction mixture (50 μl) contained 10 μl of fivefold reaction buffer, 200 μM of each of the four deoxynucleoside triphosphates, 0.2 μM of each primer, 5% DMSO, 1 U of Phusion hot start high-fidelity DNA Polymerase (Finnzymes, Vantaa, Finland) and 50 ng template (DNA or cDNA). The thermal cycling scheme comprised initial denaturation at 98 °C for 5 min, 25 cycles of denaturation at 98 °C for 45 s, annealing for 45 s at 60 °C, and extension at 72 °C for 30 s, followed by a final extension period at 72 °C for 5 min. All amplicon PCR reactions were performed in triplicate and pooled in equimolar amounts for sequencing. The Göttingen Genomics Laboratory determined the sequences of the partial 16S rRNA gene and transcript amplicons by using a 454 GS-FLX sequencer and titanium chemistry as recommended by the manufacturer (Roche, Mannheim, Germany).

Quality-filtering and denoising of the recovered 16S rRNA pyrotag reads were performed with the QIIME (1.9.1) software package (Bolyen et al. 2019) by employing the scheme outlined by Schneider et al. (2015). Forward and reverse primer sequences were removed with the *split_libraries.py* script. Pyrosequencing noise and chimeric sequences were removed with UCHIME (Edgar et al. 2011). Operational taxonomic unit (OTU) determination was performed by employing the *pick_open_reference_otus.py* script at genetic divergence level of 3 %. Taxonomic classification of OTUs was performed by *parallel_assign_taxonomy_blast.py* script against the Silva SSU database release 128 (Quast et al. 2013). The *filter_otu_table.py* script was used to remove singletons, chloroplast sequences, extrinsic domain OTUs, and unclassified OTUs. Rarefaction curves was calculated with QIIME software by using *alpha-rarefaction.py.*

### Metagenomic sequencing and data processing

The sequencing libraries were constructed and indexed with Nextera DNA Sample Preparation kit and Index kit as recommended by the manufacturer (Illumina, San Diego, CA, USA). Paired-end sequencing was performed using a HiSeq 4000 instrument (2 x 150 bp) as recommended by the manufacturer (Illumina). Raw reads were trimmed with Trimmomatic version 0.36 (Bolger et al. 2014) and verified with FastQC version 0.11.5 (Andrew, 2010). Then, reads were submitted to MG-RAST metagenomics analysis server and processed by the default quality control pipeline (Keegan et al. 2016). Microbial composition analysis was performed using MG-RAST best hit classification tool against the databases of M5RNA (Non-redundant multisource ribosomal RNA annotation) and M5NR (M5 non-redundant protein) with default settings. Functional classification was performed based on clusters of orthologous groups (COGs) and Subsystem categories with default settings. Since we mainly focused on the bacterial community, the baseline for all fractions reported referred to the reads assigned to the bacterial domain.

### Construction of metagenomic plasmid libraries and function-based screening for lipolytic activity

Lipolytic genes were screened by constructing small-insert plasmid libraries as described by Lu et al. (2019). Briefly, DNA was sheared by sonication for 3 s at 30% amplitude and cycle 0.5 (UP200S Sonicator, Stuttgart, Germany), and size-separated using a 0.8% low-melting point agarose gel. DNA fragments from 6 to 12 kb were recovered by gel extraction using the peqGold Gel Extraction kit as recommended by the manufacturer (Peqlab Biotechnologie GmbH, Erlangen, Germany). The metagenomic small-insert library was constructed using the vectors pFLD or pCR-XL-TOPO (Thermo Fisher Scientific).

Vector pFLD was digested with *PmlI* at 37°C for 2 h and dephosphorylated with 5 U Antarctic phosphatase at 37 °C for 30 min as recommended by the manufacturer (NEB, Ipswich, MA). Subsequently, the ends of DNA fragments were blunt-ended and phosphorylated by employing the Fast DNA End Repair kit (Thermo Fisher Scientific). SureClean was applied to purify DNA or vector between steps as described by the manufacturer (Bioline GmbH, Luckenwalde, Germany). Finally, metagenomic fragments and pFLD vector were ligated using T4 DNA ligase (Thermo Fisher Scientific) at 16 °C, overnight. Metagenomic DNA fragments were cloned into vector pCR-XL-TOPO following the protocol of the manufacturer recommended in the TOPO-XL-PCR cloning kit (Thermo Fisher Scientific).

To screen for lipolytic activity, *Escherichia coli* TOP10 was used as the host (Dukunde et al. 2017). Library-bearing cells were plated onto LB agar plates (15 g/L) containing 1% (v/v) emulsified tributyrin (Sigma) as the indicator substrate and the appropriate antibiotic (pFLD, 100 μg/ml Ampicillin; pCR-XL-TOPO, 50 μg/ml Kanamycin). The quality of the libraries was controlled by checking the average insert sizes and the percentage of insert-bearing *E. coli* clones (Table 1). Cells were incubated on indicator agar at 37 °C for 24 h and subsequently for 1 to 7 d at 30 °C. Lipolytic-positive *E. coli* clones were identified by the formation of clear zones (halos) around individual colonies.

**Table 1.** Summary of metagenomic libraries used for lipolytic activity screening in this and other studies

| Environmental sample<sup>b</sup> | Vector type<br>(average insert<br>size in kb) | No. of library-containing<br>clones (confirmed<br>positive hits, No. of hits<br>per million of clones) | Probability<br>(No. of hits per Gb<br>of DNA screened) | Reference |
| --- | --- | --- | --- | --- |
| Compost | Plasmid (5.3) | 675,200 (156, 213) | 43.6 | compost55 (this study) |
| Compost | Plasmid (5.6) | 234,912 (43, 183) | 32.7 | compost55 (this study) |
| Compost | Plasmid (6) | 281,281 (37, 132) | 21.9 | compost76 (this study) |
| Compost | Plasmid (6.2) | 140,747 (14, 100) | 16.1 | compost76 (this study) |
| Compost | Plasmid (3.2) | 21,000 (14, 670) | 208 | Lämmle et al. 2007 |
| Compost | Fosmid (35) | 23,400 (19, 810) | 23.2 | Kim et al. 2010 |
| Compost | Fosmid (37.5) | 1,920 (2, 1040) | 27.8 | Leis et al. 2015 |
| Compost | Fosmid (-<sup>a</sup>) | 13,000 (10, 770) | -<sup>a</sup> | Kang et al. 2011 |
| Compost | plasmid (-<sup>a</sup>) | 66,000 (6, 0.90) | -<sup>a</sup> | Popovic et al. 2017 |
| Grassland soil | Plasmid (5.7) | 510,808 (2, 0.4) | 0.714 | Nacke et al. 2011 |
| Grassland soil | Fosmid (27.8) | 50,952 (2, 40) | 1.41 | Nacke et al. 2011 |
| Forest soil | Fosmid (35) | 33,700 (8, 240) | 6.78 | Lee et al. 2004 |
| Forest soil | Plasmid (3.1) | 70,000 (3, 42) | 13.8 | Berlemont et al. 2013 |
| River surface water | BAC (50) | 8,000 (1, 120) | 2.5 | Wu and Sun 2009 |
| Hot spring biofilm | BAC (50) | 68,352 (10, 150) | 2.93 | Yan et al. 2017 |
| Surface sea water | BAC (70) | 20,000 (4, 200) | 2.86 | Chu et al. 2008 |
| Marine sediment | plasmid (4.5) | 29,000 (6, 200) | 46.0 | Ranjan et al. 2018 |
| Marine sediment | Fosmid (36) | 40,000 (19, 480) | 13.2 | Hu et al. 2010 |
| Marine mud | fosmid (40) | 40,000 (5, 120) | 3.12 | Gao et al. 2016 |
| Deep-sea hydrothermal vent | fosmid (35) | 18,000 (7, 390) | 11.1 | Fu et al. 2015 |
| Paper mill sludge | plasmid (5.1) | 15,000 (13,870) | 170 | Jia et al. 2019 |
| Activated sludge | plasmid (5.1) | 3,818 (12, 3140) | 616 | Liaw et al. 2010 |
| Activated sludge | plasmid (2.5) | 40,000 (1, 24) | 10.0 | Shao et al. 2013 |
| Solar saltern | fosmid (35) | 51,00 (1, 200) | 5.60 | Jayanath et al. 2018 |
| Oil field soil | plasmid (3.9) | 83,000 (1, 12) | 3.09 | Fan et al. 2011 |

<sup>a</sup> This information is not specified in the reference
<sup>b</sup> Except compost metagenomic libraries, only those included the full library information were listed.

The recombinant plasmid DNA derived from positive clones was isolated by using the QIAGEN plasmid mini kit (QIAGEN) and digested by *PmlI* (vector PFLD) or *EcoRI* (vector pCR-XL-TOPO) at 37 °C for 2 h. The digestion pattern was analyzed, and phenotype of positive clones was confirmed by transformation of the selected plasmids (from previous step) into the host and rescreening on indicator agar plates. In addition, lipolytic activity towards different triacylglycerides was measured qualitatively by incubating the confirmed lipolytic positive clones on agar plates emulsified with tributyrin (C4), tricaproin (C6), tricaprylin (C8), tricaprin (C10), trilaurin (C12), trimyristin (C14), or tripalmitin (C16). Formation of clearing zones (halos) on agar plates indicated lipolytic activity.

### Analysis of lipolytic genes from function-based screenings

The plasmids recovered from the confirmed positive clones were pooled in equal amounts (50 ng of each clone) for compost55 and compost76. Then, the two plasmid DNA mixtures were sequenced using an Illumina MiSeq instrument with reagent kit version 3 (2x 300 cycles) as recommended by the manufacturer (Illumina)). To remove the vector sequences, raw reads were initially mapped against vector sequences (pFLD or pCR-XL-TOPO) using Bowtie 2 (Langmead and Salzberg 2012). The unmapped reads were quality-filtered by Trimmomatic v0.30 (Bolger et al. 2014) and assembled into contigs by Metavelvet v1.2.01 (Namiki et al. 2012) and MIRA 4 (Chevreux et al. 1999). In addition, both ends of the inserts of each plasmid were sequenced using Sanger technology and the following primers: pFLD504_F (5’-GCCTTACCTGATCGCAATCAGGATTTC-3’) and pFLD706_R (5’-CGAGGAGAGGGTTAGGGATAGGCTTAC-3’) for vector pFLD, and M13_Forward (5’-GTAAAACGACGGCCAG-3’) and M13_Reverse (5’-CAGGAAACAGCTATGAC-3’) for vector pCR-XL-TOPO. The raw Sanger reads were processed with the Staden package (Staden et al. 2003). Finally, the full insert sequence for each plasmid was reconstructed by mapping the processed Sanger reads on the contigs assembled from the Illumina reads. Open reading frames (ORFs) were predicted by MetaGeneMark (Zhu et al. 2010) using default parameters. Lipolytic genes were annotated by searches against NCBI Non-redundant sequence database (http://www.ncbi.nlm.nih.gov/gorf/gorf.html).

### Family classification of lipolytic enzymes revealed from function-based screening

LEs were clustered according to the classification standard defined by Arpigny and Jaeger (1999). In order to classify LEs identified from function-based screening, we have integrated all the so far reported lipolytic families, including families I to XIX, and potential novel families reported in recent studies (Supplementary Table S1). A neighbor-joining tree was constructed with LEs identified from this study and reference proteins (Supplementary Table S2) using MEGA version 7 (Tamura et al. 2013). The robustness of the tree was tested by bootstrap analysis using 500 replications. The phylogenetic tree was depicted by GraPhlAn (Asnicar et al. 2015). To confirm the classification and group proteins in clusters, a protein sequence similarity network was generated. In a protein sequence similarity network, members in a potential isofunctional group consist of nodes (symbol) that share a sequence similarity larger than a selected value and are connected by edges (line). As similarity increases, edges decrease and finally proteins can be separated into defined clusters (Gerlt et al. 2015). In this study, a protein sequence similarity network was generated by submitting the same sequence dataset used in the phylogenetic analysis to the Enzyme Function Initiative-Enzyme Similarity Tool web server (EFI-EST; http://efi.igb.illinois.edu/efi-est/index.php) (Atkinson et al. 2009) with an E-value cutoff of ≤1e^-10^ and alignment score ≥ 16. The resulting network was visualized in Cytoscape 3.2.1 using the organic layout (Shannon et al. 2003). In addition, multiple-sequence alignments were conducted to explore the presence of catalytic residues, and conservative and distinct motifs in each lipolytic family by employing ClustalW (Larkin et al. 2007).

### Building profile Hidden Markov Model (HMM) database for sequence-based screening

A search method based on profile HMMs was developed to identify and annotate putative lipolytic genes in metagenomes (Supplementary Figure 1). In order to target homologous sequences, profile HMMs were built from multiple sequence alignments, which requires relatedness in the input protein sequences. Thus, consistent to the classification of functional-derived LEs, we generally followed the clustering system of Arpigny and Jaeger (1999).

With the exception of LEs belonging to families II and VIII, and patatin-like-proteins, LEs in the other families generally share a conserved α/β-hydrolase fold and a canonical G-x-S-x-G pentapetide around the catalytic serine (Kovacic et al. 2019). ESTHER is a database dedicated to proteins with α/β-hydrolase-fold and their classifications (Lenfant et al. 2013), containing approximately 60,000 α/β hydrolases grouped in 214 clusters so far (as of November 2019). In ESTHER, families I-XIX were integrated into an own classification with corresponding entries (http://bioweb.supagro.inra.fr/ESTHER/Arpigny_Jaeger.table). We thereby designated lipolytic families that were classified and named according to ESTHER database as ELFs (abbreviation of ESTHER Lipolytic Families). For lipolytic families that were not incorporated into the 19 families (I-XIX), their corresponding ELFs were determined by searching LEs against ESTHER database. Generally, a LE was assigned to an ELF if its BLASTp top hit (with lowest e-value) had ≥60 % amino acid identity and ≥80% query coverage. Protein sequences in all of the determined ELFs were downloaded from ESTHER database for profile HMM construction.

Firstly, multiple sequence alignments were performed with protein sequences in each ELF, using the following three algorithms and default settings: ClustalW (Thompson et al. 1994), Clustal Omega (Sievers et al. 2011) and Muscle (Edgar 2004). Subsequently, the three alignment sets were run through *hmmbuild* in HMMER3 (Eddy 2018) to create three sets of profile HMMs. Moreover, profile HMMs supplied in the ESTHER database were downloaded. Finally, four profile HMM databases were constructed by concatenating and compressing the respective set of profile HMMs using *hmmpress.* Thereafter, we designated the four profile HMM databases with respect to the corresponding alignment algorithm (clustalw-pHMMs, omega-pHMMs and muscle-pHMMs) or source (ESTHER-pHMMs).

For families II, VIII and patatin-like-proteins, profile HMMs were retrieved directly from Pfam database (Finn et al. 2014) using the searching keywords of “GDSL”, “beta-lactamase” and “patatin”, respectively. The profile HMM database was constructed as described above and designated as pfam-pHMMs, specifying for LEs in families II and VIII, and patatin-like-protein.

### Validating profile HMM database

The prediction sensitivity and specificity of the profile HMM databases were evaluated using four datasets. Dataset 1, LEs recruited in the UniProtKB database (as of November 2019) using as search strategy the EC numbers 3.1.1.1 or 3.1.1.3, and protein length between 200 to 800 amino acids. Only the prokaryotic LEs were selected for analysis (Supplementary Table S3a). Dataset 2, LEs reported in literature. Most of these enzymes were obtained through metagenomic approaches and biochemically characterized, and with a confirmed lipolytic family assignment by constructing a multiple sequence alignment and/or phylogenetic tree (Supplementary Table S3b). Dataset 3, protein sequences predicted by MetaGeneMark (Zhu et al. 2010) from identified inserts harboring functional lipolytic genes (Supplementary Table S3c). Dataset 4, randomly selected protein sequences (not recruited from ESTHER database) that were annotated in Uniprot or NCBI database as non-lipolytic proteins but with sequence homology to LEs (Supplementary Table S3d). Proteins in the four datasets were screened against the profile HMM databases successively with *hmmscan* using an E-value cutoff of ≤ 1e^-10^. The sensitivity and specificity of each database were evaluated by the recalls and false positive returns. In addition, we compared our homology-based method (profile HMMs) with the similarity-based pairwise sequence alignment method (BLAST; Altschul et al. 1990). The database for BLAST-based searching was built with the same dataset used for profile HMM construction. BLASTp was performed at an E-value cutoff of ≤ 1e^-10^.

In order to improve the accuracy for assigning proteins to lipolytic families and distinguishing “true” LEs from the non-lipolytic proteins, protein sequences were annotated by two methods and combined for final assignment. Briefly, putative lipolytic proteins (PLPs) identified by screening against the selected profile HMM database (one from clustalw-pHMMs, omega-pHMMs, muscle-pHMMs and ESTHER-pHMMs) were further searched against the ESTHER database (all entries in the database were included; as of November 2019) by BLASTp using an Evalue cutoff of ≤ 1e^-10^ (Supplementary Figure 1). A PLP was assigned to a lipolytic family only if it was annotated into the same ELF by *hmmscan* and BLASTp. Otherwise, according to the BLAST results, the remaining PLPs were either annotated as “unassigned” PLPs or non-lipolytic proteins (Supplementary Figure 1). In principle, PLPs with the best Blast hits were affiliated to the miscellaneous ESTHER families (functions were not determined, including *5_AlphaBeta_hydrolase, 6_AlphaBeta_hydrolase, Abhydrolase_7* and *AlphaBeta_hydrolase),* or other ESTHER families (with <60 % identity or <70 % query coverage) were classified as unassigned PLPs. The remaining PLPs with the best Blast hits showing ≥60 % amino acid identity and ≥70 % query coverage to the non-lipolytic ESTHER families were classified as non-lipolytic proteins.

Family annotation of PLPs obtained by screening against pfam-phmms were confirmed by a further scan against the CATH HMMs database (Knudsen and Wiuf 2010) using the Github repository *cath-tools-genomescan* (https://github.com/UCLOrengoGroup/cath-tools-genomescan). PLPs were assigned to lipolytic families VIII and II, or patatin-like-proteins only if the PLP was assigned to the specific Funfams (functional families) dedicated to lipolytic-related activities, which were inferred from the functionally characterized LEs and gene ontology (GO) annotations (Supplementary Table S4). Additionally, based on our literature search, the LEs in family VIII were generally restricted to PLPs with sequence length between 350 to 450 amino acids. In other cases, the PLP was grouped into non-lipolytic proteins.

For the unassigned PLPs, these sequences show low similarity to any ESTHER family with known function or CATH Funfams, and hence, could contain novel lipolytic or non-lipolytic proteins. Non-lipolytic proteins were excluded from the downstream analysis.

### Sequence-based screening for putative lipolytic genes

Sequence-based screening for putative lipolytic genes in the two compost metagenomes were performed as described above. Briefly, the processed metagenomic short reads were assembled into contigs with SPADES version 3.10 (Bankevich et al. 2012). Then, protein sequences were deduced from PROKKA v1.14.5 annotation (Seemann 2014). In order to obtain full-length lipolytic genes, only proteins with amino acid sequence length between 200 and 800 amino acids were retained. Subsequently, the resulting protein sequences were screened against the selected profile HMM databases using *hmmscan* (Eddy 2011) with an E-value cutoff of ≤1e^-10^. Identified PLPs were further assigned into different lipolytic families as described above (Supplementary Figure 1). Moreover, the lipolytic family classification of assigned PLPs was confirmed by constructing the protein sequence similarity network (Atkinson et al. 2009). The phylogenetic origins of PLP-encoding genes and their corresponding contigs were determined using Kaiju web server (http://kaiju.binf.ku.dk/server; Menzel et al. 2016). Phylogenetic distributions of assigned PLPs in each lipolytic family were visualized via Circos software (Krzywinski et al. 2009).

### Comparative analysis of metagenomic datasets

A total of 175 assembled metagenomes from 15 different habitats were retrieved from the Integrated Microbial Genomes and Microbiomes database (IMG/M). These included metagenomes from anaerobic digestor active sludges (ADAS, n=9), agriculture soils (AS, n=10), composts (COM, n=18), grassland soils (GS, n=11), human gut systems (HG, n=16), hypersaline mats (HM, n=7), hydrocarbon resource environments (HRE, n=6), hot springs (HS, n=14), landfill leachates (LL, n=10), marine sediments (MS, n=12), marine waters (MW, n=10), oil reservoirs (OR, n=13), river waters (RW, n=11), tropical forest soils (TFS, n=14) and wastewater bioreactors (WB, n=13) (Supplementary Table S5). Data processing including open reading frame prediction in assembled contigs and taxonomic assignment of the corresponding deduced protein sequences were conducted by the IMG/M built-in pipelines (Chen et al. 2017). The protein sequences were downloaded from IMG/M database and used in the sequence-based screening as described above (Supplementary Figure 1).

For comparative analysis, the abundance of PLP-encoding genes in each metagenome were normalized according to the method described by Kaminski et al. (2015). The normalized count is in units of LPGM (Lipolytic hits Per Gigabase per Million mapped genes). Unless otherwise stated, LPGM values were used for all calculations. Heatmap was built in R v3.5.2 (R Core Team, 2016) with the function *heatmap.2* using the “Heatplus” package (Ploner et al., 2020). The heatmap hierarchical clustering was performed with “vegan” package (vegdist = “bray”, data.dist = “ward.D”). Non-metric multidimensional scaling (NMDS) was also performed with the “vegan” package (Oksanen et al. 2018). The analysis of similarities (ANOSIM) was performed with 9,999 permutations using PAST 4 (Hammer et al. 2001). The phylogenetic annotation of PLPs was retrieved from IMG/M. Association networks between habitats and phylogenetic distribution of PLPs at genus level were generated by mapping significant point biserial correlation values with the “indicspecies” package in R (Cáceres 2013). Only genera with significant correlation coefficients *(P* = 0.05) were included. The resulting bipartite networks were visualized with Cytoscape v3.5 by using the *edge-weighted spring embedded layout* algorithm, whereby the habitats were source nodes, genera target nodes and edges (lines connecting nodes) weighted positive associations between genera and specific habitat or habitats combinations.

In addition, due to the ambiguity of unassigned PLPs, all analyses were performed successively using two datasets: (1) only assigned PLPs, as the consideration of excluding the potential non-lipolytic ones, (2) assigned and unassigned PLPs combined (total PLPs), in order to include all the possible lipolytic ones. This paper mainly focuses on the assigned PLPs for the sake of accuracy, but the comparative analysis of total PLPs was also performed.

## Data availability

The short reads and insert sequences were submitted to NCBI databases. Metagenomic short reads are available at in the NCBI sequence read archive (SRA) under accession numbers SRR13115019 (compost55) and SRR13115018 (compost76) and 16S rRNA pyrotag reads under SAMN06859928 (compost55 genes), SAMN06859946 (compost55 transcripts), SAMN06859935 (compost76 genes) and SAMN06859953 (compost76 transcripts). The insert sequences of the plasmids are available in GenBank under accession numbers MW408002--MW408112 (Supplementary Table S9).

## Results and Discussion

### Phylogenetic and functional profile of microbes in the compost metagenomes

During the heating-up process of composting, the succession of microorganisms plays a key role in degrading organic matter (Dougherty et al. 2012). In this study, the bacterial community compositions in two compost samples with different pile core temperatures of 55 (compost55) and 76 °C (compost76) were revealed by amplicon-based sequencing of 16S rRNA genes (DNA-based, total community) and transcripts (RNA-based, active community) (Supplementary Figures S2 and S3). To extend the taxonomic analysis, the environmental DNA from both metagenomes were also directly sequenced (Supplementary Table S6). Generally, the bacterial community determined by direct sequencing were consistent with that derived from 16S rRNA gene-based analysis. The bacterial phyla *Actinobacteria, Proteobacteria, Firmicutes, Bacteroidetes* and *Chloroflexi* were predominant (relative abundance >5 %) in compost55 and compost76 (Supplementary Figures S3 and S4). This is in agreement with previous studies of bacterial communities in thermophilic composts (Ryckeboer et al., 2003; Antunes et al., 2016; Zhen et al., 2018; Zhou et al., 2018). Differences were detected, which were derived mainly from the different feedstock composition (wood chips vs. kitchen waste) and composting conditions (core temperature 55 vs. 76 °C). *Actinobacteria* was the most abundant phylum (> 25 %) in compost55 (Supplementary Figures S3 and S4), which is accordance with the bacterial communities in composts using mainly plant material as feedstock (Yu et al. 2007; Zhang et al. 2014). In compost76, members of the *Firmicutes* were most abundant (> 55 %), which was also reported for composts harboring high-nitrogen feedstock, such as animal manure and kitchen waste (Niu et al. 2013; Antunes et al. 2016; Ma et al. 2018; Zhou et al. 2019). The 16S rRNA gene and transcript analysis (Supplementary Table S7) revealed genera that were present (> 1%) in compost55 such as *Brockia, Rhodothermus, Thermobispora, Longispora, Geobacillus, Filomicrobium* and *Thermomonospora,* and in compost76 such as *Symbiobacterium, Calditerricola* and *Thermaerobacter* were among the typical bacterial taxa previously identified in composting processes (Ryckeboer et al. 2003; Antunes et al. 2016; Yu et al. 2018; Zhou et al. 2018).

Additionally, the metagenomic data were searched against the COG and subsystem databases to assess the functions prominent in compost microbes (Supplementary Figure S5). In principle, compost55 and compost76 share similar metabolic patterns (Supplementary Figure S5). Particularly, the broad diversity and abundance of gene functions in carbohydrate metabolism and transport (COG) and carbohydrates (subsystems) indicated that composts were potential candidates for exploring biocatalysts (Hu et al. 2010a; Leis et al. 2015; Wang et al. 2016; Egelkamp et al. 2019). Notably, the COG category of lipid transport and metabolism as well as the subsystems category of fatty acids, lipids, and isoprenoids were more abundant in the compost55 community than in the compost76 community, suggesting a higher possibility to identify lipolytic genes in the compost55 metagenome.

### Function-based screening of LEs in compost metagenomes

In this study, four metagenomic libraries were prepared to probe the diversity of LEs from compost microbes by the function-driven approach using tributyrin-containing indicator agar (Table 1). Overall, approximately 4.89 and 2.56 Gb of cloned compost DNA were screened, yielding 199 and 51 positive clones for compost55 and compost76, respectively. Previous studies have used various vectors such as BACs, fosmids and plasmids for function-based screening of LEs from different bioresources (Lee et al. 2004; Lämmle et al. 2007; Kim et al. 2010; Nacke 2011; Berlemont et al. 2013; Shao et al. 2013; Leis et al. 2015; Jia et al. 2019). The hit rate to recover a lipolytic-positive clone ranged from 0.714 to 208 per Gb of cloned DNA (Table 1). Among the compost metagenomic libraries, the targeting probability towards a LE in our study ranged from 16.1 to 43.6 per Gb and is generally consistent with the values from other studies (Lämmle et al. 2007; Kim et al. 2010; Leis et al. 2015). Also notably, the targeting probabilities in metagenomic libraries from compost and sludge are generally higher than those from other environments, such as grassland, forest soil and river water (Wu and Sun 2009; Nacke 2011; Berlemont et al. 2013). According to Liaw et al. (2010), the targeting probability and/or hit rate for discovering a lipolytic clone is largely attributed to the sample source.

Other studies further suggested that samples subjected to specific enrichment processes, such as composting and waste treatment procedures, usually resulted in a high hit rate (Mayumi et al. 2008; Kang et al. 2011; Popovic et al. 2017).

The insert sizes of the recovered plasmids (250 in total) with a confirmed phenotype ranged from 1,038 to 12,587 bp. In all inserts, at least one putative gene showing similarities to known genes encoding lipolytic enzymes was detected. In total, 210 and 60 lipolytic genes were identified from compost55 and compost76 libraries, respectively. To identify unique and full-length LEs, the amino acid sequences deduced from the corresponding lipolytic genes were clustered at 100 % identity. This resulted in 115 (92 for compost55, 23 for compost76, with 7 shared by both samples) unique and full-length LEs (Supplementary Table S8). The length of the unique LEs ranged from 223 to 707 amino acids, with calculated molecular masses from 23.9 to 72.3 kDa (Supplementary Table S9). Forty of these showed the highest similarity to esterases/lipases from uncultured bacteria, and one (EstC55-13) to an enzyme from an uncultured archaeon. Among them, seven LEs showed the highest identity (53 to 65 %) to lipolytic enzymes obtained during function-based screening of metagenomes derived from marine sediment (Hu et al. 2010b), forest topsoil (Lee et al. 2004), mountain soil (Ko et al. 2012), activated sludge (Liaw et al. 2010), wheat field (Stroobants et al. 2015) and compost (Okano et al. 2015). In the remaining 34 cases, the matching esterases/lipases were mainly detected by sequence-based metagenomic surveys of composts (15 LEs), soil (7 LEs), marine sediment (6 LEs) and marine water (3 LEs).

### Functionally derived LEs are affiliated with various LE families

The LEs identified through function-based screening were grouped into families based on the classification system reported by Arpigny and Jaeger (1999). With the increasing amount of reports on LEs, claims of new families have been reported (Arpigny and Jaeger 1999; Jeon et al. 2011; Wang et al. 2013; Esteban-Torres et al. 2014; Fang et al. 2014; Rahman et al. 2016; Castilla et al. 2017). In this study, we integrated 29 so-called “novel” families into the classification system for phylogenetic analysis. As shown in the phylogenetic tree (Figure 1), LEs were assigned to 12 families, including families I, II, III, IV, V, VII, VIII, XVII, EM3L4 (Lee et al. 2011), FLS18 (Hu et al. 2010b), EstGS (Nacke et al. 2011), LipT (Chow et al. 2012), patatin-like-proteins and tannases (Supplementary Table S8). The majority of the LEs were affiliated to families V (25 LEs), VIII (21 LEs), IV (15 LEs), I (8 LEs) and patatin-like-proteins (9 LEs). Noteworthy, 7 LEs could not be classified into any known lipolytic family, indicating new branches of LEs. In agreement with previous studies (Arpigny and Jaeger 1999; Glogauer et al. 2011; Akmoussi-Toumi et al. 2018), the “true lipases”, which can hydrolyze long-chain substrates (≥ C10) were all affiliated to family I (Figure 1). The remaining LEs exhibiting a preference for short-chain substrates (<C10) were esterases.

**Figure 1.**
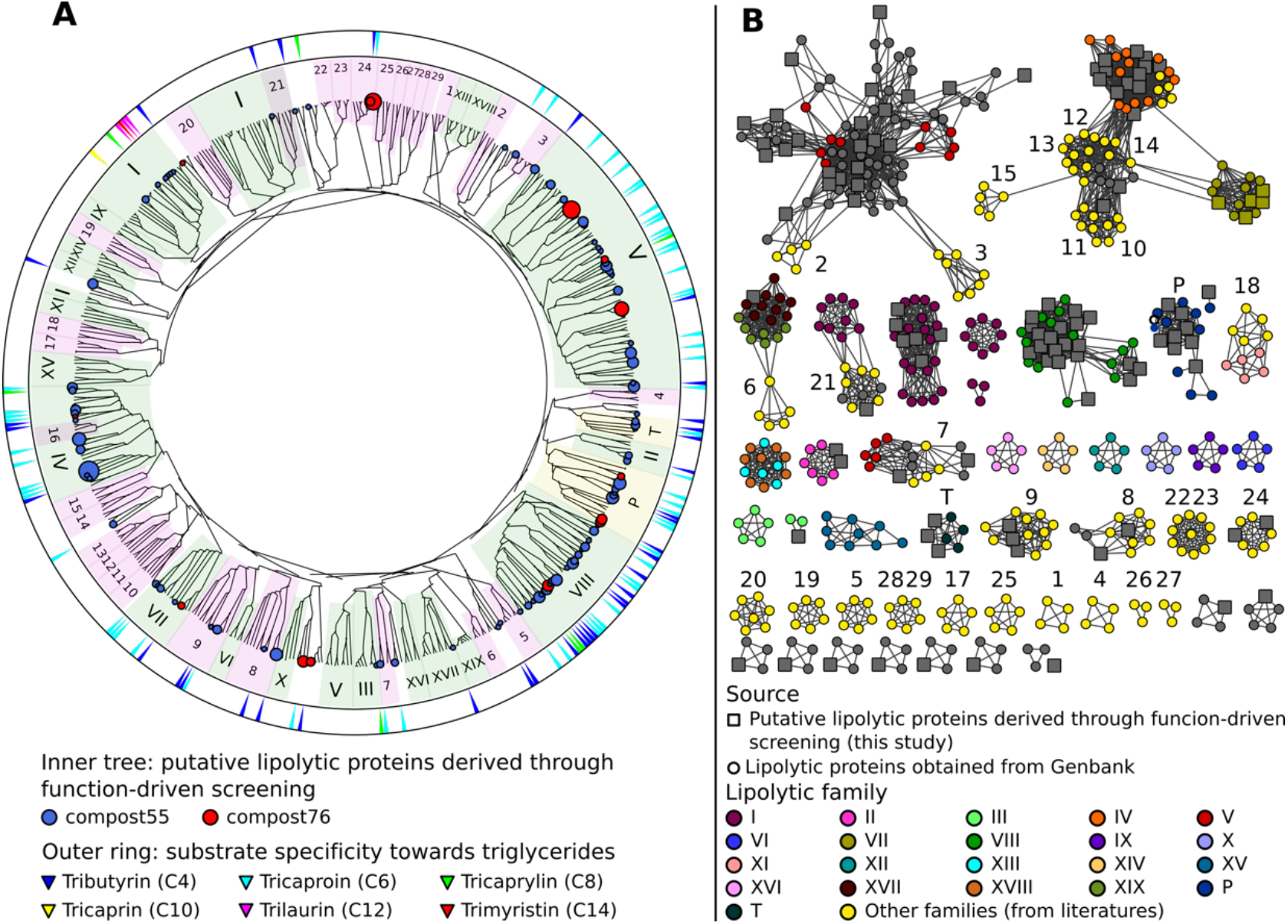
Classification of LEs identified through the function-driven approach. **A,** Unrooted phylogenetic tree was constructed using FA-identified LEs in this study obtained and references retrieved from GenBank (Supplementary Table S2). Phylogenetic tree was constructed using MEGA 7 with neighbor-joining method. The robustness of the tree was tested by bootstrap analysis with 500 replications. Inner tree: the circles represent LEs detected in compost55 (blue) and compost76 (red), sized by abundance (counts of replicates). LEs assigned to families of I-XIX were shaded in green background. Patatin-like-proteins and tannases (designated as P and T, respectively) were shaded in yellow. Other recent reported lipolytic families were shaded in magenta: 1, Est22 (Li et al. 2017); 2, EstL28 (Seo et al. 2014); 3, Rv0045c (Guo et al. 2010); 4, EstGX1 (Jiménez et al. 2012); 5, EstLiu (Rahman et al. 2016); 6, EstY (Wu and Sun 2009); 7; EstGS (Nacke et al. 2011); 8, EM3L4 (Lee et al. 2011); 9, FLS18 (Hu et al. 2010b); 10, Est903 (Jia et al. 2019); 11, EstJ (Choi et al. 2013); 12, PE10 (Jiang et al. 2012); 13, Est12 (Wu et al. 2013); 14, EstDZ2 (Zarafeta et al. 2016); 15, Est9x (Jeon et al. 2009); 16, Lip10 (Guo et al. 2016); 17, EstGH (Nacke et al. 2011); 18, EML1 (Jeon et al. 2009); 19, FnL (Yu et al. 2010); 20, EstP2K (Ouyang et al. 2013); 21, LipA (Couto et al. 2010); 22, LipSM54 (Li et al. 2016); 23, MtEst45 (Lee 2016); 24, LipT (Chow et al. 2012); 25, EstSt7 (Wei et al. 2013); 26, Rlip1 (Liu et al. 2009); 27, EstA (Chu et al. 2008); 28, FLS12 (Hu et al. 2010b); 29, lp_3505 (Esteban-Torres et al. 2014). Outer ring: substrate specificity of corresponding clones towards different carbon chain length (C4 – C14) of triglycerides. **B,** Protein sequence similarity network of LEs belonging to different families. Networks were generated from all-by-all BLAST comparisons of amino acid sequences from the same dataset used for the construction of the phylogenetic tree. Each node represents a sequence. *Larger square nodes* represent LEs derived from function-based screening performed in this study. *Small circle nodes* represent LEs retrieved from GenBank. Nodes were arranged using the *yFiles* organic layout provided in Cytoscape version 3.4.0. Each edge in the network represents a BLAST connection with an E-value cutoff of ≤1e^-16^. At this cut-off, sequences have a mean percent identity and alignment length of 36.3% and 273 amino acids, respectively.

To verify the classification result, a protein sequence similarity network was built (Figure 1). The network visualizes relationships among evolutionarily related proteins and is usually considered as an approach complementary to the phylogenetic analysis (Atkinson et al. 2009; Gerlt et al. 2015). At a threshold of 1×10^-16^, the network produced clusters that almost matched all the lipolytic families, with the same classification results as obtained by phylogenetic analysis (Figure 1).

Multiple sequence alignments revealed the catalytic residues and conserved motifs in each family (Supplementary Figure S6). For LEs that harbor the canonical α/β-hydrolase fold, the catalytic triad is consistently composed of a nucleophilic serine, an aspartic acid/glutamic acid and a histidine residue (Nardini & Dijkstra, 1999). Most of these LEs contain the conserved motif Gly-x-Ser-x-Gly in which the catalytic serine is embedded (Supplementary Figure S6). Alternatively, three LEs in family I show variations of this conserved motif. The variations were Ala-x-Ser-x-Gly, Thr-x-Ser-x-Gly (Diamond et al. 2019) and Ser-x-Ser-x-Gly (Dalcin Martins et al. 2018) (Supplementary Figure S6).

Family II LEs share a canonical α/β/α-hydrolase fold, which is characterized by a conserved hydrophobic core consisting of five β-strands and at least four α-helices (Akoh et al. 2004). As shown in Supplementary Figure 6a, there are four homology blocks and one conserved residue in each block (serine, glycine, asparagine, and histidine, respectively), which is essential for catalysis (Akoh et al. 2004; Hong et al. 2008). The structures of family VIII enzymes show remarkable sequence similarities to β-lactamases and penicillin-binding proteins (Bornscheuer 2002). Site-directed mutagenesis demonstrated that the catalytic triad is composed of serine and lysine located in a Ser-X-X-Lys motif, and a tyrosine (Supplementary Figure 6) (Biver and Vandenbol 2013; Kovacic et al. 2019). The patatin-like-proteins display an α/β/α-hydrolase fold, in which a central six-stranded beta-sheet is sandwiched between alpha-helices front and back (Banerji and Flieger 2004). Unlike the catalytic triad of Ser-Asp/Glu-His for most lipolytic proteins, the catalytic Ser-Asp dyad is responsible for the catalytic activity of patatin-like-proteins. In addition, they also contained the Gly-x-Ser-x-Gly motif with the catalytic serine embedded (Supplementary Figure 6).

### Development of a LE profile HMM database for sequence-based screening

Profile HMMs are statistical models that convert patterns, motifs and other properties from a multiple sequence alignment into a set of position-specific hidden states, i.e. frequencies, insertions, and deletions (Reyes et al. 2017). Profile HMMs are sensitive in detecting remote homologs. Thus, they have been utilized to detect, e.g. viral protein sequences (Skewes-Cox et al. 2014; Bzhalava et al. 2018), antibiotic resistance genes (Gibson et al. 2015), GDSL esterase/lipase family genes (Li et al. 2019) in metagenomes.

In this study, a total of 32 ELFs were determined for profile HMM database construction (Supplementary Table S10). Subsequently, four profile HMM databases (Omega-phmms, Muscle-phmms, Clustalw-phmms, ESTHER-phmms) specific for LEs affiliated to α/ ß hydrolase superfamily were constructed. Each database consists of 32 profile HMMs (Supplementary Table S11). The prediction sensitivity and specificity of the four databases were evaluated using four datasets (Table 2). All of the four databases obtained high recalls for the datasets 1, 2 and 3 (Table 2), with the highest ones for omega-pHMMs (4,446 in total), followed by muscle-pHMMs (4,444), clustalw-pHMMs (4,425) and ESTHER-pHMMs (4,425). Noteworthy, omega-pHMMs did not identify any false positive LEs for dataset 3. Thus, omega-pHMMs was chosen for downstream screening. In addition, we compared omega-pHMMs with the pairwise sequence alignment method (BLASTp) for their ability to predict LEs. The omega-pHMM database exhibited improved sensitivity for datasets 1, 2, and 3. In total, 135 more LEs were identified using omega-pHMMs than BLASTp (Table 2).

**Table 2.** Comparison of profile HMM databases based on different alignment tools to detect LEs

| Datasets<sup>a</sup> | Nr. of LEs<br>($\alpha/\beta$<br>hydrolase) | Nr. of LEs<br>(non- $\alpha/\beta$<br>hydrolase) | Recall of LEs ($\alpha/\beta$ hydrolase) | | | | | Recall of LEs<br>(non-$\alpha/\beta$<br>hydrolase)<br>pfam-phmms |
| --- | --- | --- | --- | --- | --- | --- | --- | --- |
|  |  |  | Omega-phmms | Muscle-phmms | Clustalw-phmms | ESTHER-phmms | BLASTp |  |
| Dataset 1 | 4382 | 554 | 4243 | 4244 | 4228 | 4225 | 4122 | 554 |
| Dataset 2 | 130 | 32 | 125 | 125 | 121 | 124 | 117 | 32 |
| Dataset 3 | 80 | 36 | 78 | 75 | 76 | 76 | 70 | 36 |
| Dataset 4 | 68 | 0 | 56 | 55 | 53 | 53 | 51 | 0 |

<sup>a</sup> Dataset 1, LEs from Uniprot database; Dataset 2, recently reported LEs; Dataset 3, MetaGeneMark-predicted proteins from inserts conferring lipolytic activity; Dataset 4, potential non-lipolytic proteins with homology to LEs.

The accuracy of omega-pHMMs for lipolytic family assignment was also assessed. For datasets 2 and 3, we achieved high precision of annotating LEs to the known lipolytic families, with the exception of LEs from novel families (Supplementary Table S12). Dataset 4 included non-lipolytic proteins, such as epoxide hydrolases, dehalogenases and haloperoxidases and exhibited significant homology with LEs in subfamilies V.1 and V.2 (Arpigny and Jaeger 1999). Our “homology-based” method only differentiated part of these non-lipolytic homologies from “true” LEs (Table 2).

To improve the annotation accuracy, putative lipolytic proteins (PLPs) were further searched against the entire ESTHER database by BLASTp. By combining the annotations from both methods (Supplementary Figure 1), these “novel” LEs in datasets 2 and 3 were correctly identified as “unassigned”, in terms of not assigned to any known ELF (Supplementary Table S12). Moreover, almost all of the non-lipolytic proteins (> 92 %) in dataset 4 were distinguished from LEs (Supplementary Table S12).

To identify LEs affiliated to families VIII and II, and patatin-like proteins, enzymes were successively screened against pfam-pHMMs and CATH HMMs database. For the first three datasets, all the LEs in the three families were correctly identified by screening against pfam-pHMMs (Table 2, Supplementary Table S13).

As demonstrated in other sequence-based metagenomic approaches (Liu et al. 2015b; Maimanakos et al. 2016; Azziz et al. 2019), our screening strategy is also vastly dependent on the completeness and accuracy of the reference databases (ESTHER and CATH database in this study). Hence, PLPs exhibiting closest similarity to members affiliated to the miscellaneous ESTHER families or no ESTHER/CATH hits returned, were classified into the “unassigned” group in this study (Supplementary Table S12). This might have resulted in an underestimation of assigned lipolytic proteins (Supplementary Table S13).

### Sequence-based screening confirmed compost metagenomes as reservoir for putative lipolytic genes

Initial screening of the assembled metagenomes of compost55 and compost76 resulted in the identification of 4,157 and 2,234 PLPs, respectively. Among them, 1,234 and 759 were further assigned into 28 and 26 families, respectively. The assigned PLPs belonged mainly to family VIII, hormone-sensitive lipase-like proteins, patatin-like proteins, II, A85-Feruloyl-Esterase, Carb_B_Bacteria and homoserine transacetylase (Supplementary Figure S7). The family assignment was also verified by constructing a protein sequence similarity network (supplementary Figure S8). The large number of unassigned PLPs (2,460 for compost55 and 1,208 for compost76) indicated the presence of candidates for novel lipolytic families.

The assigned PLPs were generally of bacterial origin (>95 %), and mainly affiliated to the phyla (> 5 %) *Actinobacteria*, *Proteobacteria*, *Firmicutes* and *Bacteroidetes* (Figure 2). The corresponding contigs were also taxonomically assigned and exhibited a similar phylogeny as seen for the embedded PLP-encoding gene sequences (Figure 2).

**Figure 2.**
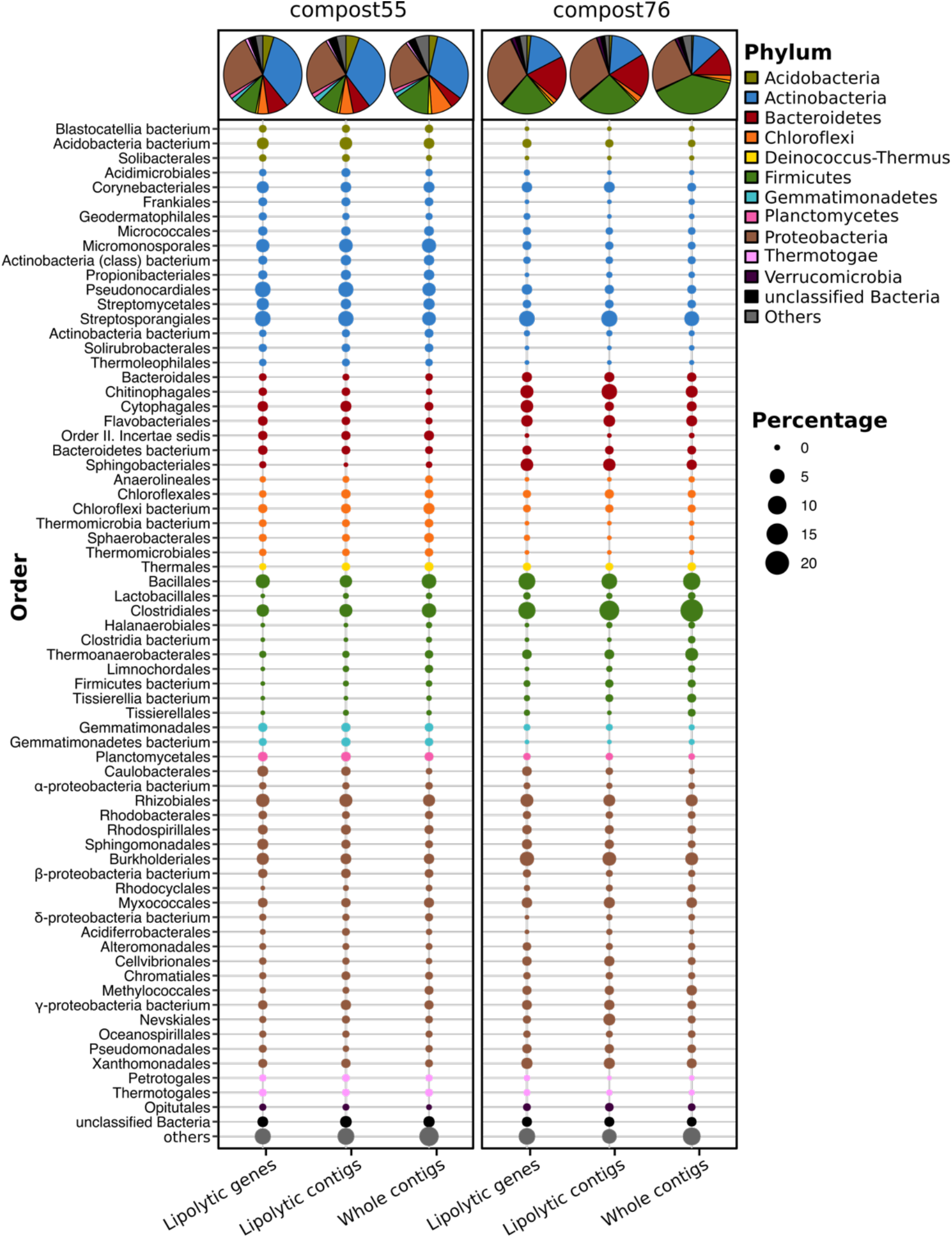
Phylogenetic distribution of assigned PLP-encoding genes identified in compost55 and compost76 metagenomes. The phylogenetic origin of PLP-encoding genes, the contigs harboring these genes, and the whole assembled contigs were annotated by Kaiju (Menzel et al. 2016), and expressed as the proportion of the respective total counts in each sample. The pie charts represent the taxonomic composition at phylum level. Taxa with an abundance of less than 1% were grouped into “others”.

Members of the *Actinobacteria* have been reported as important biomass degraders (Ryckeboer et al. 2003; Hubbe et al. 2010; Wang et al. 2016; Lewin et al. 2016). In this study, 34.7 (compost55) and 15.8 % (compost76) of the assigned PLPs originated from *Actinobacteria.* At genus level, the assigned PLPs were affiliated to *Mycobacterium, Actinomadura, Thermomonospora, Streptomyces, Micromonospora, Pseudonocardia* and *Thermobifida* (Supplementary Table S14). Members of these genera have been reported as producers for lipases/esterasses (Wei et al. 1998; Alisch et al. 2004; Chahinian et al. 2005; Guo et al. 2010; Hu et al. 2010a; Brault et al. 2012; Mander et al. 2014; Sriyapai et al. 2015). Moreover, some of the corresponding families, such as *Micromonosporaceae, Streptomycetaceae* and *Thermomonosporaceae,* are commonly found in thermophilic composts (Schloss et al. 2003; Blaya et al. 2016; Lima-Junior et al. 2016).

*Proteobacteria* are also an abundant source for the assigned PLPs in compost55 (26.2 %) and compost76 (31.4 %) (Figure 2). Popovic et al., (2017) identified 80 LEs, of which, 65 % were proteobacterial origin by screening of 16 metagenomic DNA libraries prepared from seawater, soils, compost and wastewater. In our study, lipolytic genes exhibited high taxonomic diversity at genus level, they were distributed across 97 and 111 genera for compost55 and compot76, respectively (Supplementary Table S14).

The assigned PLPs affiliated to *Firmicutes* originated mainly from *Clostridiales* and *Bacillales* (Figure 2). By analyzing the microbial diversity and metabolic potential of compost metagenomes, members of *Clostridiales* and *Bacillales* were shown to play key roles in degradation of different organic compounds (Martins et al. 2013; Antunes et al. 2016). *Bacteroidetes* is the fourth most abundant phylum for assigned PLPs in compost55 (8.4 %) and compost76 (18.8 %) (Figure 2). At genus level, the assigned PLPs derived mainly from *Rhodothermus* in compost55, and *Sphingobacterium, Flavobacterium, Niastella* and *Flavihumibacter* in compost76 (Supplementary Table S14). Members of these genera are known as important fermenters during composting (Neher et al. 2013; Antunes et al. 2016; Lapébie et al. 2019).

Strikingly, the phylogenetic distribution of assigned PLPs in each sample, to some extent, corresponded well to the taxonomic composition revealed from the whole contigs (Figure 2) but with minor differences in the rank abundance order. The 16S rRNA amplicon (Supplementary Figure S3) and metagenomic datasets (Supplementary Figure S4) also showed a composition of dominant orders similar to that deduced from lipolytic genes/contigs (Figure 2). Wang et al. (2016) revealed that the phylogenetic distribution of CAZyme genes in the rice straw-adapted compost consortia was in accordance to its microbial composition. By mapping resistance gene dissemination between humans and their environment, Pehrsson et al. (2016) found that resistomes across habitats were generally structured by bacterial phylogeny along ecological gradients.

### Comparison between function-driven and sequence-based screening of LEs

Metagenomics allows tapping into the rich genetic resources of so far uncultured microorganisms (Simon and Daniel 2011) through function-driven or sequence-based approaches. The function-driven strategy targets a particular activity of metagenomic library-bearing hosts (Ngara and Zhang 2018). In this way, we identified 13 novel LEs (Supplementary Table S9, Supplementary Figure S6b), which confirmed functional screening as a valuable approach for discovering entirely novel classes of genes and enzymes, particularly when the function could not be predicted based on DNA sequence alone (Reyes-Duarte et al. 2012; Lam et al. 2015; Villamizar et al. 2017).

The sequence-based screening strategy is also frequently used due to the easy access to a wealth of metagenome sequence data and continuous advances in bioinformatics (Chan et al. 2010; Liu et al. 2015c; Maimanakos et al. 2016). Based on the ESTHER and Pfam database, our profile HMM-based search approach efficiently provided an overview of PLP distribution in the two compost metagenomes (Supplementary Figure S7). The hit rate for LEs was higher by sequence-based than by function-based screening, but the sequence-based derived hits need to be functionally verified. In addition, we noticed that only part of the functional screening-derived lipolytic genes were identified during sequence-based screening. By mapping the metagenomic short reads to the functional screening-derived lipolytic genes, 63 genes (out of 115 lipolytic genes in total) had a coverage of 100 % and 88 of ≥ 99 % (Supplementary Table S15). The BLAST-based comparison between lipolytic genes derived from function-driven and sequence-based approaches indicated that 31 genes from each approach exhibited 100 % sequence identity, and 64 over 99% identity (Supplementary Table S16).

In summary, function-driven and sequence-based strategies have advantages and disadvantages. The function-driven screenings are generally constrained by factors, such as labor-intensive operation, limitations of the employed host systems and low hit rate (Simon and Daniel 2011). However, function-based approaches are activity-directed, and sequence-and database-independent, thus, they bear the potential to discover entirely novel genes for proteins of interest (Rabausch et al. 2013; Lam et al. 2015). Sequencing-based screening, on the other hand, is effective in identifying sequences and potential genes encoding targeted biomolecules in metagenomes. Sequence-based screens largely rely on the used search algorithms, and quality and content of the reference databases to infer the functions of discovered candidate genes (Ngara and Zhang 2018). Thus, the best way to explore novel molecules is to combine the two approaches (Barriuso and Jesús Martínez 2015). Function-driven screens can be employed to complete and verify reference database entries on which sequence-based screening is dependent on. In addition, sequence-based approaches can serve as a pre-selection step for function-driven screens and analysis (Chan et al. 2010; Masuch et al. 2015; Pehrsson et al. 2016; Streit et al. 2018).. The known novel LEs identified by function-based approaches and the functional enzymes identified in this study were employed to expand the LE-specific profile HMM database and annotate the PLPs derived from sequence-based screening.

### Assigned PLPs are distributed by ecological factors

In this study, 175 metagenomes representing various ecology niches were selected for sequence-based searching of PLPs. In total, we have screened approx. 1.23 billion genes in 65 Gbp of assembled metagenomes and recovered approx. 0.22 million (absolute counts) PLP-encoding genes. The assigned PLPs (34 % of the total counts) were normalized to LPGM values for comparative analysis. In accordance with the function-based screening, samples subjected to certain enrichment processes, particularly lipid-related, tend to have a higher hit rate (Figure 3). For example, samples with high LPGM values were derived from a hydrocarbon resource environment and oil reservoir that are enriched with oil-degrading microbes (Liu et al. 2015a; Hu et al. 2016; Vigneron et al. 2017; Liu et al. 2018), and composts and wastewater bioreactors that are reservoir for microbes decomposing organic compounds (Dougherty et al. 2012; Silva et al. 2012; Antunes et al. 2016; Berini et al. 2017). Intriguingly, samples from human gut systems were also candidates for LEs (LPGM values > 7500). The human intestinal microorganisms play an import role in degrading diet components into metabolizable molecules (Wang et al. 2015). The function- and sequence-based study of human gut metagenomes have proved that the human gut microbiome is a rich source for various carbohydrate active enzymes (Li et al. 2009; Turnbaugh et al. 2009; Tasse et al. 2010; Moore et al. 2011).

**Figure 3.**
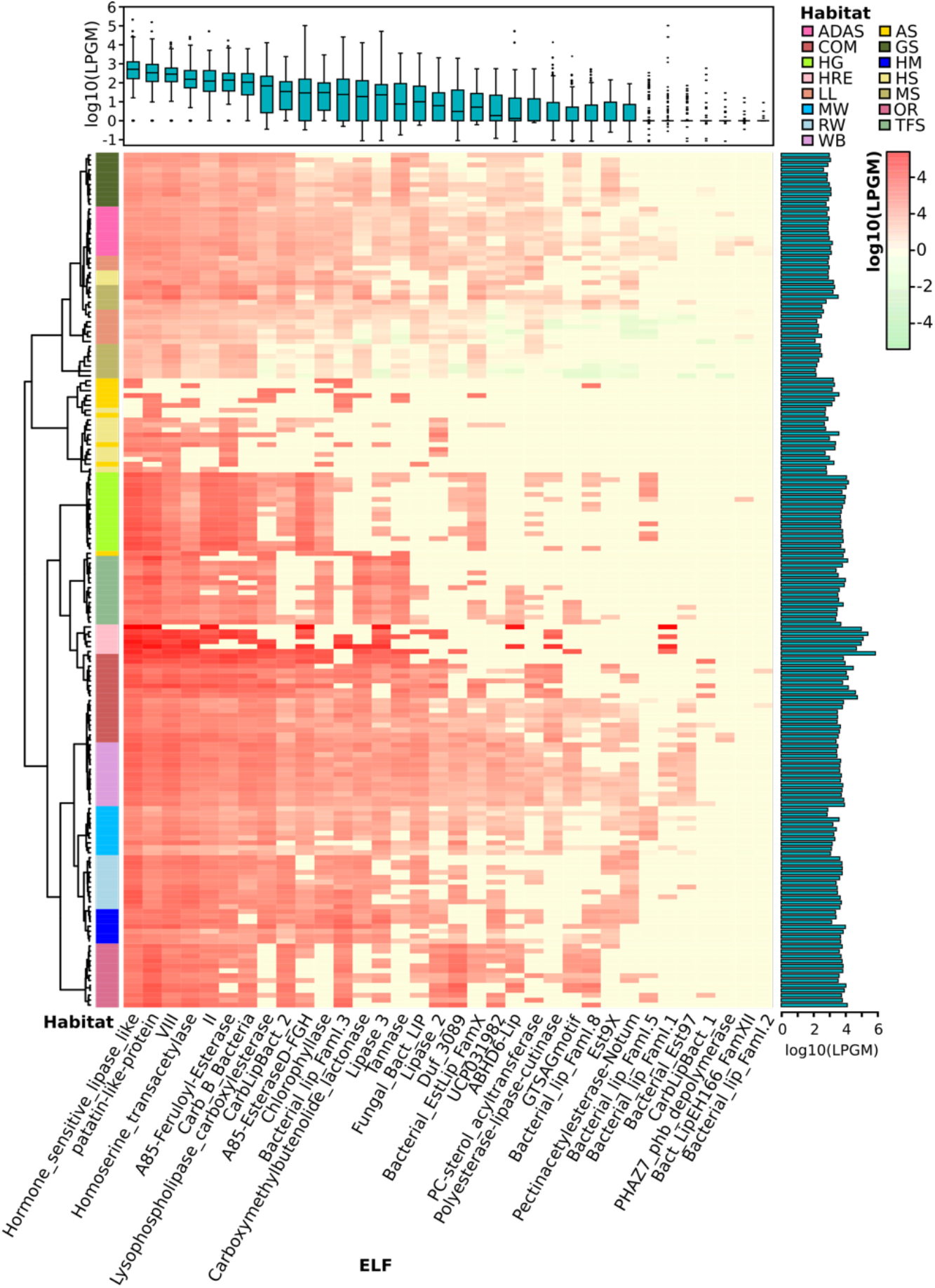
Lipolytic family profile of assigned PLPs across samples. Hierarchical clustering analysis of the lipolytic family profile in each sample was performed using the Ward.D clustering method and Bray-Curtis distance matrices. LPGM values were log_10_ transformed. The color intensity of the heat map (light green to red) indicates the change of LPGM values (low to high). The habitats are depicted by different colors. The lipolytic family profile in each sample was generally clustered by habitat (overall R value = 0.621, P<0.001, ANOSIM test). The boxplot (top) represents the distribution of the assigned PLPs in each ELF across samples. Mean values (n= 175 samples) are given. The bar plot (right) shows the total abundance of assigned PLPs by summing up the abundance in each family of each sample. Abbreviations of habitats: ADAS, anaerobic digestor active sludge; AS, agricultural soil; COM, compost; GS grassland soil; HG, human gut; HM, hypersaline mat; HRE, hydrocarbon resource environment; HS, hot spring; LL, landfill leachate; MS, marine sediment; MW, marine water; OR, oil reservoir; RW, river water; TFS, tropical forest soil; WB, wastewater bioreactor; ELF, ESTHER lipolytic family.

Overall, the assigned PLPs were classified into 34 lipolytic families (Fig 3). Members of the Hormone-sensitive_lipase_like and patatin-like-protein families were most abundant (average LPGM values across samples > 2000), followed by families of A85-EsteraseD-FGH, VIII and Bacterial_lip_FamI. 1 (average LPGM values > 700) (Fig 3). However, no family was shared by all samples. Nevertheless, members from families of Hormonesensitive-lipase-like, patatin-like-proteins, VIII, homoserine transacetylase, II and A85-Feruloyl-Esterase were detected in more than 90 % of samples (Figure 3). Enzymes belonging to families of PHAZ7_phb_depolymerase, Bact_LipEH166_FamXII and Bacterial_lip_FamI.2 were not or only rarely detected (< 6 % of all samples) and showed a low abundance (LPGM values < 1). The prevalence and abundance of a lipolytic family revealed by the sequence-based screening are dependent on the distribution of corresponding target genes in the microbial consortia (Wang et al. 2016). Taking members from the “abundant” family Hormone-sensitive_lipase_like as example, the corresponding genes are widely distributed in more than 1,200 species as recorded in the ESTHER database so far. This was, somehow, also reflected by the function-based screening, in which a large proportion of the identified LEs belonged to family Hormone-sensitive_lipase_like. In contrast, according to the ESTHER database, only 23, 8 and 6 species harboring LEs were affiliated to the “rare” families like PHAZ7_phb_depolymerase, PC-sterol_acyltransferase and Bact_LipEH166_FamXII, respectively (Supplementary Figure S12).

To investigate the distribution of assigned PLPs that cause the observed lipolytic family profiles across samples and habitats, a matrix with LPGM values representing the abundance of PLPs per lipolytic family identified in each metagenome was generated. The lipolytic family profiles clustered by habitats (Figure 3), which was confirmed by NMDS (stress level 0.2268; Supplementary Figure S9). ANOSIM (Clarke 1993) was used to pairwise compare the multivariate (group) differences of lipolytic family profiles between habitats. A R value-based matrix was generated among habitats (Supplementary Figure S9), a high R value (between 0 to 1) indicated a high group dissimilarity between two habitats. Generally, each habitat exhibited a distinctive pattern of lipolytic family profiles (overall R value = 0.6168; Supplementary Table S17). For example, PLPs detected in agricultural soils were only present in eight lipolytic families with low abundances. In contrast, PLPs in composts were detected in almost all lipolytic families, and with remarkably high abundance in families such as Hormone-sensitive_lipase_like, patatin-like-protein and VIII (Supplementary Figure S10). Notably, the lowest group dissimilarity was observed between the habitats compost and wastewater bioreactor (R=0.1941, P < 0.001, ANISOM; Supplementary Figure S9). The analysis of lipolytic profiles across habitats allows selecting suitable habitats for function-based screening, e.g. targeting LEs of a specific family or with some properties for desired applications. Metagenomes from composts are promising for recovering LEs in families LYsophospholipase_carboxylesterase (family VI), CarbLipBact_2 (family XIII-2/XVIII) and CarbLipBact_1 (family XIII-1) (Supplementary Figure S11).

### The phylogenetic distribution of assigned PLPs

More than 98 % of the assigned PLPs were encoded by bacterial community members. Although LEs are widely encoded in various microbial genomes (Hausmann and Jaeger 2010; Ramnath et al. 2016; Kovacic et al. 2019a), the assigned PLPs were mainly derived from the bacterial phyla *Proteobacteria* (66.5 %), *Bacteroidetes* (12.5 %), *Actinobacteria* (7.7 %), *Firmicutes* (6.7 %) (Figure 4). This is consistent with the taxonomic origin of reference LEs in ESTHER database (Supplementary Figure S12). Moreover, enzymes from members of *Proteobacteria* were dominant in almost all lipolytic families (Figure 4). At genus level, the phylogenetic origins of assigned PLPs were scattered across approx. 2,000 bacterial genera, with enriched abundance in the genera *Acinetobacter, Pseudomonas, Bacteroides, Bradyrhizobium* and *Mycobacterium* (average LPGM values across samples > 180). Many of the LEs from these genera were described as exoenzymes (Rudek and Haque 1976; Gilbert 1993;

**Figure 4.**
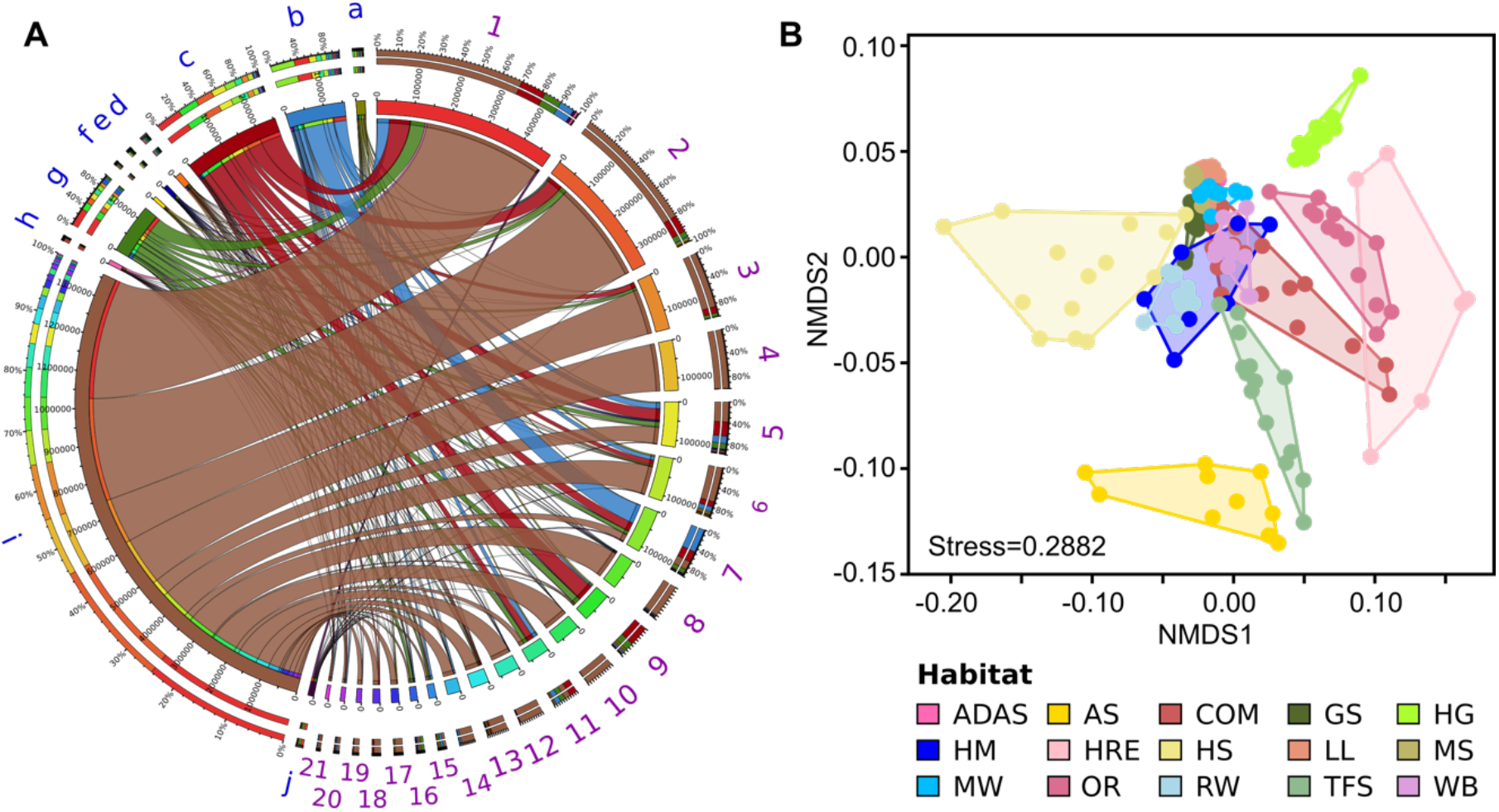
Phylogenetic distribution of assigned PLPs. **A**, phylogenetic distributions of assigned PLPs in abundant bacterial phyla possessing PLP-encoding genes across all the samples. The abundance inferred from LPGM values matrix of assigned PLPs per family identified in each bacterial phylum was generated by summing the corresponding LPGM values across all samples. The width of bars from each bacterial phylum and functional enzyme family indicates their relative abundances across all samples. a-j are bacterial phyla (in blue): a, *Acidobacteria;* b, *Actinobacteria;* c, *Bacteroidetes;* d, *Chloroflexi;* e, *Cyanobacteria;* f, *Deinococcus-Thermus;* g, *Firmicutes;* h, *Planctomycetes;* i, *Proteobacteria;* j, *Verrucomicrobia.* 1-21 are lipolytic families (in purple): 1, Hormone-sensitive_lipase_like; 2, patatin-like-protein; 3, A85-EsteraseD-FGH; 4, Bacterial_lip_FamI.1; 5, VIII; 6, Homoserine_transacetylase; 7, II; 8, Lipase_3; 9, A85-Feruloyl-Esterase; 10, ABHD6-Lip; 11, Carb_B_Bacteria; 12, Bacterial_lip_FamI.3; 13, Lysophospholipase_carboxylesterase; 14, Carboxymethylbutenolide_lactonase; 15, CarbLipBact_2; 16, Chlorophyllase; 17, Tannase; 18, Polyesterase-lipase-cutinase; 19, Duf_3089; 20, Fungal_Bact_LIP; 21, Lipase_2. Only phyla and lipolytic families with a relative abundance > 0.5 % are shown. **B**, Non-metric multidimensional scaling (NMDS) analysis of phylogenetic distribution of assigned PLPs across samples based on Bray-Curtis distances at bacterial genus level. Only genera with a mean LPGM values of ≥ 0.5 across all the samples were included. Abbreviations of habitats: ADAS, anaerobic digestor active sludge; AS, agricultural soil; COM, compost; GS grassland soil; HG, human gut; HM, hypersaline mat; HRE, hydrocarbon resource environment; HS, hot spring; LL, landfill leachate; MS, marine sediment; MW, marine water; OR, oil reservoir; RW, river water; TFS, tropical forest soil; WB, wastewater bioreactor; ELF, ESTHER lipolytic family.

Snellman and Colwell 2004; Guo et al. 2010; Liu et al. 2018). Notably, a similar taxonomic enrichment at genus level was also observed for the reference LEs in ESTHER database as 960 LEs were encoded by *Mycobacterium,* 410 by *Pseudomonas,* 260 by *Bacteroides,* 166 by *Acinetobacter,* and 164 by *Bradyrhizobium* species (Supplementary Table S18).

The taxonomic origin of assigned PLPs at genus level varied significantly across habitats (overall R value = 0.821, P <0.01), especially for the human gut system, oil reservoir and hydrocarbon resource environment (Supplementary Figure S14). The average R value was 0.98, 0.97 and 0.94, respectively (Supplementary Table S19). The lowest dissimilarity was observed between compost and wastewater bioreactor (R value = 0.2317, P <0.001, ANISOM).

### Habitats harboring prevalent and distinct microbial clusters are main drivers of PLP distribution

Bipartite association networks have been used to identify microbial taxa responsible for shifts in community structures (Hartmann et al. 2015; Dukunde et al. 2019). In this study, a bipartite association network was constructed to visualize the associations between bacterial members at genus level that harbor lipolytic genes and habitats or habitat combinations (Figure 5). 225 of the total 712 genera, were not significantly separated in abundance and frequency by habitat. These belonged mainly to *Proteobacteria* (82 genera), *Bacteroidetes* (43 genera), *Firmicutes* (33 genera), and *Actinobacteria* (25 genera) (Supplementary Table S20). These nonsignificant genera were conserved across different habitats, generally represented the “indigenous group” (Hartmann et al. 2015; Wemheuer et al. 2017), and formed the core microbiota harboring lipolytic genes. This core microbiota was also an indication of the prevalence of lipolytic genes across microbes and habitats (Bornscheuer 2002; Hasan et al. 2006; Barriuso and Jesús Martínez 2015; Berini et al. 2017). In contrast, the significant indicators, with respect to the “characteristic group” (Rime et al. 2016; Dukunde et al. 2019), highlighted the bacterial genera that were responsible for the change of assigned PLPs distribution across habitats (Figure 5). Particularly, the indicators associated with only one habitat defined the distinctiveness of microbiota in each habitat (Hartmann et al. 2015). In this study, the unique-associated indicators accounted for 76% of all significant indicators (Supplementary Table S20). This strongly resembled the ANISOM result, in which the high overall R value (0.8199) suggested a significant distinctiveness of the phylogenetic origins of assigned PLPs across habitats (Supplementary Table S19). With respect to each habitat, a high ratio of unique-associated indicators to the total significant genera in a habitat generally indicated a high R value (Pearson’s r correlation = 0.6672, P < 0.01, linear regression; Supplementary Figure S15). For example, out of the 75 indicators that were significantly associated to the habitat hydrocarbon resource environment, 65 were unique-associated indicators, with a mean R value of 0.93 (Supplementary Table 19). This is also the case for the habitats oil reservoir (60 out of 75; mean R value = 0.96) and human gut system (35 out of 41; mean R value = 0.97).

**Figure 5.**
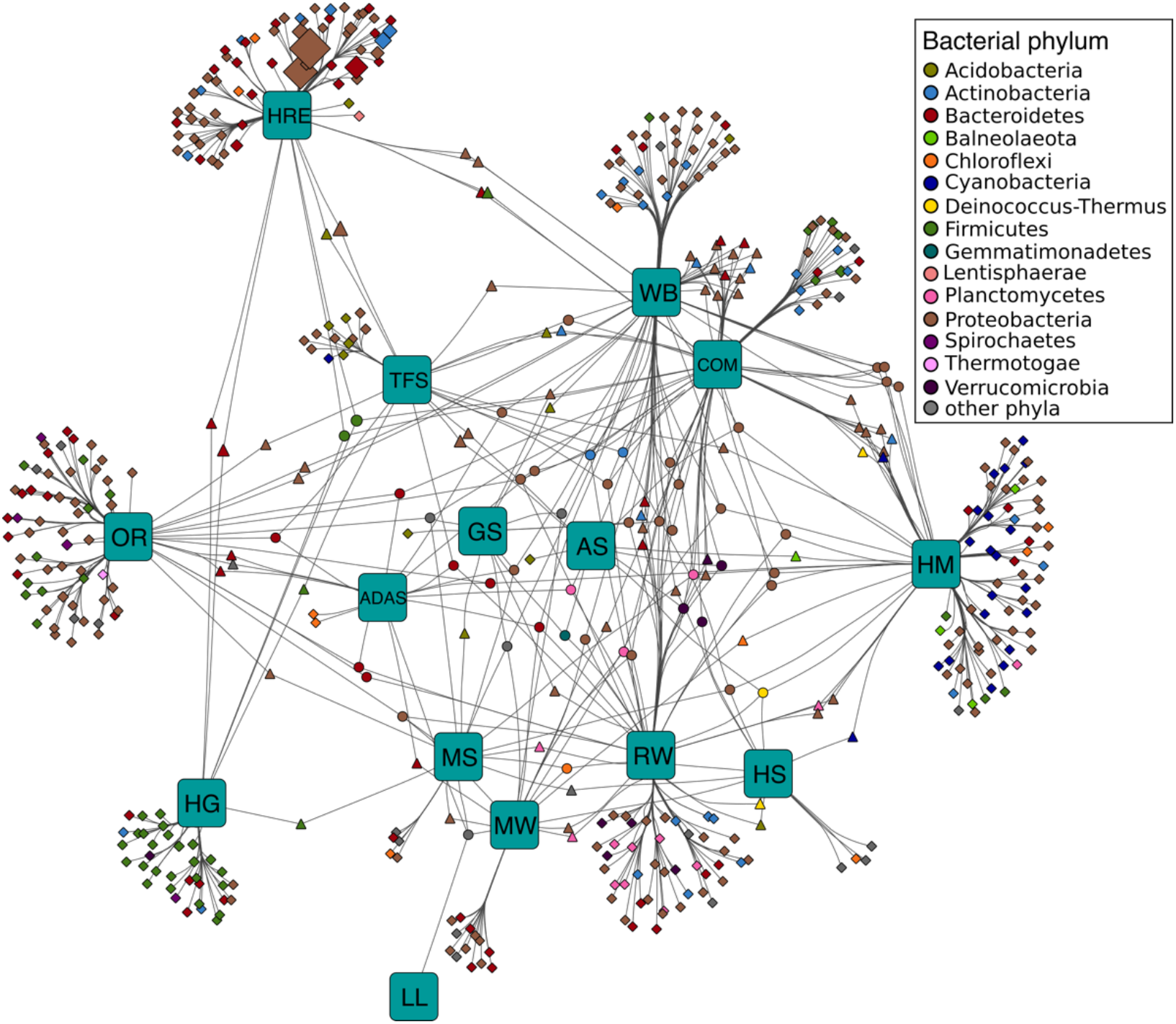
Association networks between bacterial origin of assigned PLPs at genus level and habitats. The abundance of PLPs in each genus per sample was presented by LPGM values, and only genera with mean LPGM values of ≥ 0.5 across all the samples were used. Source nodes (rounded squares) represent habitats, target node represent bacterial genera (circles, diamonds and triangles), and edges represent associations between habitats and bacterial genera. Target node size represent its mean abundance inferred from LPGM values across habitats. Target node is colored according to its phylogenetic origin at phylum level. The length of edges is weighted according to association strength. Unique clusters, which associate with only one habitat, consist of nodes shaped as diamond. Triangle and circle nodes represent genera with significant cross association between two and more habitats, respectively. Data only represents genera that showed significant positive association with habitats (P = 0.05). For ease of visualization, edges were bundled together, with a stress value of 3. Abbreviations of habitats: ADAS, anaerobic digestor active sludge; AS, agricultural soil; COM, compost; GS grassland soil; HG, human gut; HM, hypersaline mat; HRE, hydrocarbon resource environment; HS, hot spring; LL, landfill leachate; MS, marine sediment; MW, marine water; OR, oil reservoir; RW, river water; TFS, tropical forest soil; WB, wastewater bioreactor; ELF, ESTHER lipolytic family.

Only a small fraction of the indicators exhibited cross associations between two (14 % of the total indicators) or more (10 %) habitats. Nevertheless, the 29 cross-associated indicators between habitats compost and wastewater bioreactor explained the low dissimilarity of phylogenetic distributions of assigned PLPs between the two habitats (R=0.2317, P < 0.001, ANISOM).

Similar to the “indigenous group”, the “characteristic group” consisted mainly of genera affiliated to *Proteobacteria* (224 genera), *Bacteroidetes* (72), *Firmicutes* (49) and *Actinobacteria* (36). Among them, proteobacterial genera largely characterized the major habitats, such as tropical forest soil (83 %), wastewater bioreactor (67 %), hypersaline mat (52 %), hydrocarbon resource environment (51 %), oil reservoir (51 %), compost (50 %), marine water (46 %), river water (45 %), and grassland soil (42 %), whereas *Bacteroidetes* and *Firmicutes* characterized the human gut system (68 %) and the active sludge of an anaerobic digestor (53 %) (Supplementary Table S20). Noteworthy, the unique-associated indicators affiliated to *Cyanobacteria* were primarily enriched in the hypersaline mat (95 % indicators), which is also the case for *Planctomycetes* and *Verrucomicrobia* in river water (88 and 80 %, respectively). Pehrsson et al (2015) detected a link between microbial community structure and functional gene repertoire. This link could be extended to the distribution pattern of indicators in our study. For example, various studies have proved that the microbes in human gut systems were dominated by *Firmicutes* (Mahowald et al. 2009; Vital et al. 2014; Rinninella et al. 2019), which in turn leads to the *Firmicutes-*dominated indicators for lipolytic genes (Figure 5). Among all the habitats, only hypersaline mats were featured by the *Cyanobacteria-*dominated oxygenic layer for photosynthesis (Sørensen et al. 2005; Lindemann et al. 2013), which explained that almost all the *Cyanobacteria* indicators were associated with the hypersaline mat (Figure 5).

## Conclusions

In this study, two compost samples (compost55 and compost76) were used for metagenomic screening of potential lipolytic genes. Through the function-driven screening, 115 unique LEs were identified and assigned into 12 known lipolytic families. In addition, 7 LEs were not assigned to any known family, indicating new branches of lipolytic families. Our results show that functional screening is a promising approach to discover novel lipolytic genes, particularly for targeted genes, whose function is not predicted based on DNA sequence alone. For sequence-based screening, we have developed a search and annotation strategy specific for putative lipolytic genes in metagenomes (Supplementary Figure 1). Our profile HMM-based searching methods yielded higher sensitivity (recall) for LEs than the BLASTp-derived counterpart. The annotation method also remarkably increased the specificity and accuracy in distinguishing lipolytic from non-lipolytic proteins. With this sequence-based strategy, we identified the putative lipolytic genes within the two compost metagenomes. Analysis of the phylogenetic origin of these genes indicated a potential link between microbial taxa and their functional traits. By comparing the lipolytic hits identified by function-driven and sequence-based screening, we conclude that the best way for exploring and exploiting LEs is to combine both approaches.

In addition, assembled metagenomes from samples of various habitats were used for comparative analysis of the PLP distribution. We profiled the lipolytic family and phylogenetic origin of assigned PLPs for each sample. The two profiles were generally driven by ecological factors, i.e. the habitat. Moreover, the habitat also determined the conserved and distinctive microbial groups harboring the putative lipolytic genes.

PLPs were also mainly enriched in the bacterial phyla *Proteobacteria*, *Bacteroidetes*, *Actinobacteria*, *Firmicutes* (Supplementary Figure S16). The profile of the phylogenetic total PLPdistribution in each sample clustered also by habitats (Supplementary Figures S17, S18 and S19). The bipartite association network identified the conserved and distinctive microbial groups harboring PLP-encoding genes among the habitats (Supplementary Tables S21 and S22). Thus, our study provided a sequence-based strategy for effective identification and annotation of potential lipolytic genes in assembled metagenomes. More importantly, through this strategy, the overview of how the lipolytic genes distributed ecologically (in various habitats), functionally (in different lipolytic enzyme families) and phylogenetically (in diverse microbial groups) is an advantage for novel and/or industrially relevant LE identification.

## Supplementary Materials

The supplementary materials are also available online. The Supplementary Figures S1-S19 and the Supplementary Tables S1-S22 are presented in two files (Supplementary Figures S1-S19.pdf and Supplementary Tables S1-S22.xlsx, respectively)

## Acknowledgments

We thank Dr. Silja Brady and Mechthild Bömeke for providing technical assistance.

## Authors Approvals

All authors have seen and approved the manuscript. The manuscript has not been accepted or published elsewhere.

