## Supplementary Figures S1 to S19 for "Metagenomic screening for lipolytic genes reveals an ecology-clustered distribution pattern"

**Supplementary Figures S1 – S19 for the manuscript:**

### Metagenomic screening for lipolytic genes reveals an ecology-clustered distribution pattern

Mingji Lu[1], Dominik Schneider[1], Rolf Daniel[1*]

[1]Department of Genomic and Applied Microbiology, Institute of Microbiology and Genetics, Georg-August-University of Göttingen, Grisebachstraße 8, 37077, Göttingen, Germany

[*]Corresponding author: Rolf Daniel, Genomic and Applied Microbiology & Göttingen Genomics Laboratory, Institute of Microbiology and Genetics, University of Göttingen, Grisebachstr. 8, 37077 Göttingen, Germany. Phone: +49-551-3933827, Fax: +49-551-3912181,

#### Content

**Supplementary Figure S1.** Overall workflow for identification of lipolytic enzymes (LEs) through function-driven and sequence-based approaches in this study.

**Supplementary Figure S2.** Rarefaction curves of subsampled OTUs for 16S rRNA genes (DNA level) and transcripts (RNA level) in compost55 and compost76 at 97% similarity.

**Supplementary Figure S3.** Phylogenetic composition of bacterial communities in compost55 and compost76, revealed from 16S rRNA genes (DNA-level) and transcripts (RNA-level).

**Supplementary Figure S4.** Phylogenetic composition of bacterial communities in compost55 and compost76 as annotated by the MG-RAST platform.

**Supplementary Figure S5.** Functional distribution pattern in compost55 (blue filled circle) and compost76 (red filled circle) microbial consortia.

**Supplementary Figure S6a.** Multiple sequence alignments of partial amino acid sequences harboring homologous catalytic regions of homology. Lipolytic enzymes were from reported families.

**Supplementary Figure S6b.** Multiple sequence alignments of partial amino acid sequences harboring homologous catalytic regions of homology. Lipolytic enzymes were from putative novel families identified in this study.

**Supplementary Figure S7.** Phylogenetic distribution at phylum level of assigned PLPs in the most abundant lipolytic families.

**Supplementary Figure S8.** Protein Sequence similarity network for classification of assigned PLPs obtained by screening against from compost55 and compost76 assembled metagenomes.

**Supplementary Figure S9.** Functional lipolytic family profiles of assigned PLPs in different samples.

**Supplementary Figure S10.** Distribution of lipolytic families revealed from assigned PLPs of each habitat.

**Supplementary Figure S11.** Lipolytic families showing significant changes in abundance across different habitats.

**Supplementary Figure S12.** Phylogenetic origins of LEs in ESTHER database at phylum level in the most abundant lipolytic families.

**Supplementary Figure S13.** Taxonomic origins at genus level of the assigned PLPs across samples.

**Supplementary Figure S14.** Phylogenetic distribution of the assigned PLPs at phylum level in each habitat.

**Supplementary Figure S15.** Linear regression, the x was the ratio of unique indicators to the total significant indicators in a habitat, as demonstrated by the bipartite association network shown in Figure 5.

**Supplementary Figure S16.** Phylogenetic origin of the total PLPs (assigned and unassigned PLPs combined) at (A) domain and (B) phylum level.

**Supplementary Figure S17.** Heat map of the taxonomic origins at genus level of total PLPs across samples.

**Supplementary Figure S18.** Analysis of the phylogenetic profile at genus level of total PLPs across samples.

**Supplementary Figure S19.** Phylogenetic distribution of the total PLPs at phylum level of each habitat.

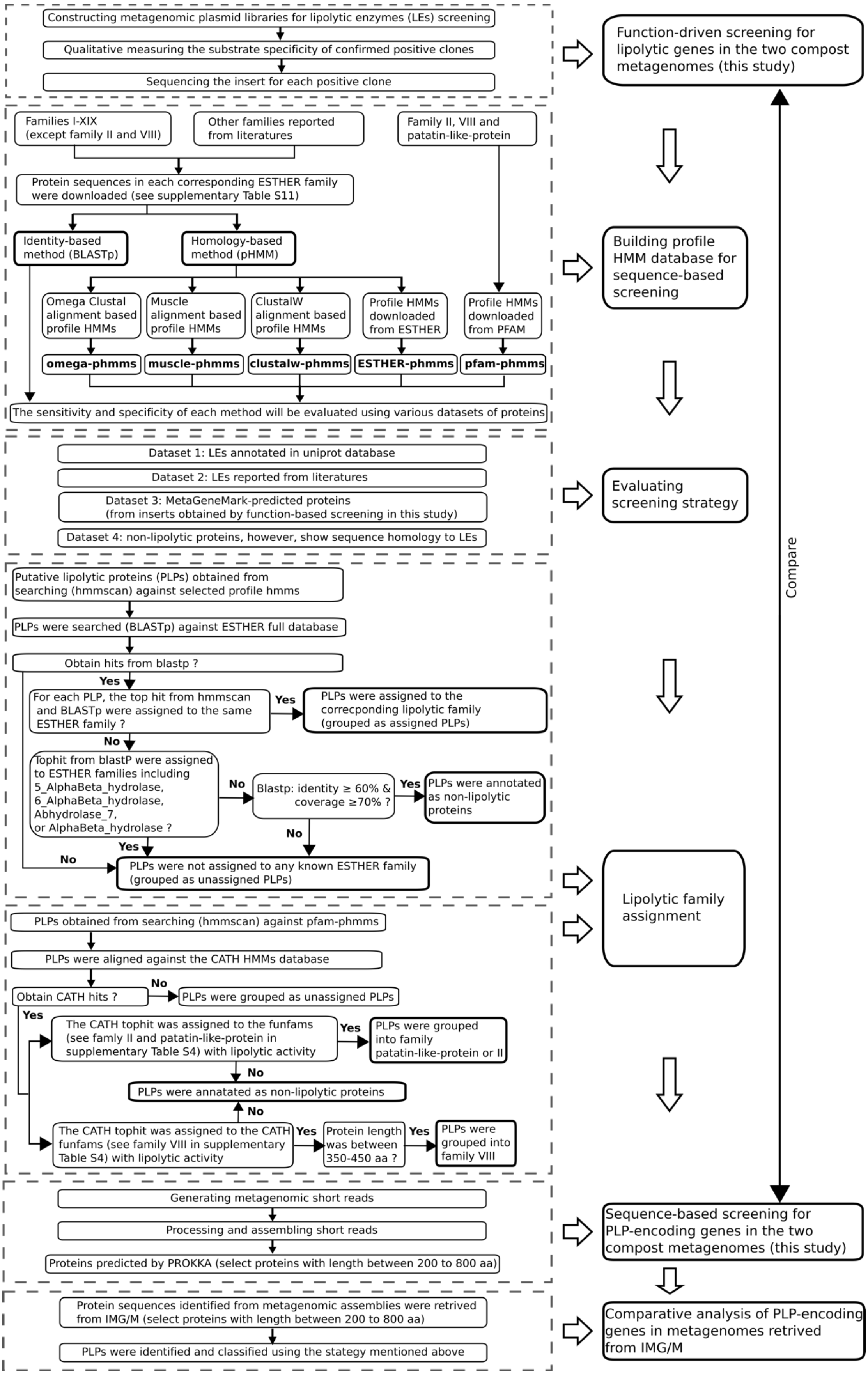

Constructing metagenomic plasmid libraries for lipolytic enzymes (LEs) screening
Qualitative measuring the substrate specificity of confirmed positive clones
Sequencing the insert for each positive clone
Function-driven screening for lipolytic genes in the two compost metagenomes (this study)
Families I-XIX (except family II and VIII)
Other families reported from literatures
Family II, VIII and patatin-like-protein
Protein sequences in each corresponding ESTHER family were downloaded (see supplementary Table S11)
Identity-based method (BLASTp)
Homology-based method (pHMM)
Omega Clustal alignment based profile HMMs
Muscle alignment based profile HMMs
ClustalW alignment based profile HMMs
Profile HMMs downloaded from ESTHER
Profile HMMs downloaded from PFAM
omega-phmms
muscle-phmms
clustalw-phmms
ESTHER-phmms
pfam-phmms
The sensitivity and specificity of each method will be evaluated using various datasets of proteins
Building profile HMM database for sequence-based screening
Dataset 1: LEs annotated in uniprot database
Dataset 2: LEs reported from literatures
Dataset 3: MetaGeneMark-predicted proteins (from inserts obtained by function-based screening in this study)
Dataset 4: non-lipolytic proteins, however, show sequence homology to LEs
Evaluating screening strategy
Compare
Putative lipolytic proteins (PLPs) obtained from searching (hmmscan) against selected profile hmms
PLPs were searched (BLASTp) against ESTHER full database
Obtain hits from blastp ?
Yes
For each PLP, the top hit from hmmscan and BLASTp were assigned to the same ESTHER family ?
Yes
PLPs were assigned to the correcponding lipolytic family (grouped as assigned PLPs)
No
Tophit from blastP were assigned to ESTHER families including 5_AlphaBeta_hydrolase, 6_AlphaBeta_hydrolase, Abhydrolase_7, or AlphaBeta_hydrolase ?
No
Blastp: identity ≥ 60% & coverage ≥70% ?
Yes
PLPs were annotated as non-lipolytic proteins
No
Yes
No
PLPs were not assigned to any known ESTHER family (grouped as unassigned PLPs)
Lipolytic family assignment
PLPs obtained from searching (hmmscan) against pfam-phmms
PLPs were aligned against the CATH HMMs database
Obtain CATH hits ?
No
PLPs were grouped as unassigned PLPs
Yes
The CATH tophit was assigned to the funfams (see famly II and patatin-like-protein in supplementary Table S4) with lipolytic activity
Yes
PLPs were grouped into family patatin-like-protein or II
No
PLPs were annatated as non-lipolytic proteins
No
The CATH tophit was assigned to the CATH funfams (see family VIII in supplementary Table S4) with lipolytic activity
Yes
Protein length was between 350-450 aa ?
Yes
PLPs were grouped into family VIII
Generating metagenomic short reads
Processing and assembling short reads
Proteins predicted by PROKKA (select proteins with length between 200 to 800 aa)
Sequence-based screening for PLP-encoding genes in the two compost metagenomes (this study)
Protein sequences identified from metagenomic assemblies were retrived from IMG/M (select proteins with length between 200 to 800 aa)
PLPs were identified and classified using the stategy mentioned above
Comparative analysis of PLP-encoding genes in metagenomes retrived from IMG/M

**Supplementary Figure S1**. Overall workflow for identification of lipolytic enzymes (LEs) through function-driven and sequence-based approaches in this study.

Firstly, LEs were identified by function-based screening of constructed metagenomic libraries. Positive clones were collected and inserts harboring lipolytic genes were sequenced. Lipolytic genes were subsequently revealed and classified. As for sequence-based screening, a search method based on the profile Hidden Markov Models (HMMs) was developed to identify and annotate the putative lipolytic proteins (PLPs) in assembled metagenomes. LEs can be generally divided into two major groups: α/β hydrolase or not α/β hydrolase. For LEs belong to the α/β hydrolase superfamily, four LE-specific profile HMM databases were retrieved (omega-phmms, muscle-phmms, clustalw-phmms and ESTHER-phmms). For LEs that are not α/β hydrolases, profile HMMs were retrieved from the pfam database (pfam-phmms). The prediction sensitivity and specificity of each profile HMM database were evaluated using four datasets, and the best one was selected for subsequent analysis. The lipolytic family assignment of PLPs obtained by screening against the selected profile HMM database (one of omega-phmms, muscle-phmms, clustalw-phmms and ESTHER-phmms) were generally conducted by combining the annotations from *hmmscan* against the profile HMM database and *blastp* against the full ESTHER database. For PLPs obtained by screening against pfam-phmms, the annotation was performed by the subsequent screening against the CATH HMMs database. Based on the strategies for sequence-based screening and lipolytic family assignment, PLPs in the two compost assembled metagenomes were identified and annotated. The results from function-driven and sequence-based screening were also compared. Finally, assembled metagenomes in various habitats were retrieved from the IMG/M database, and PLPs were identified for comparative analysis.

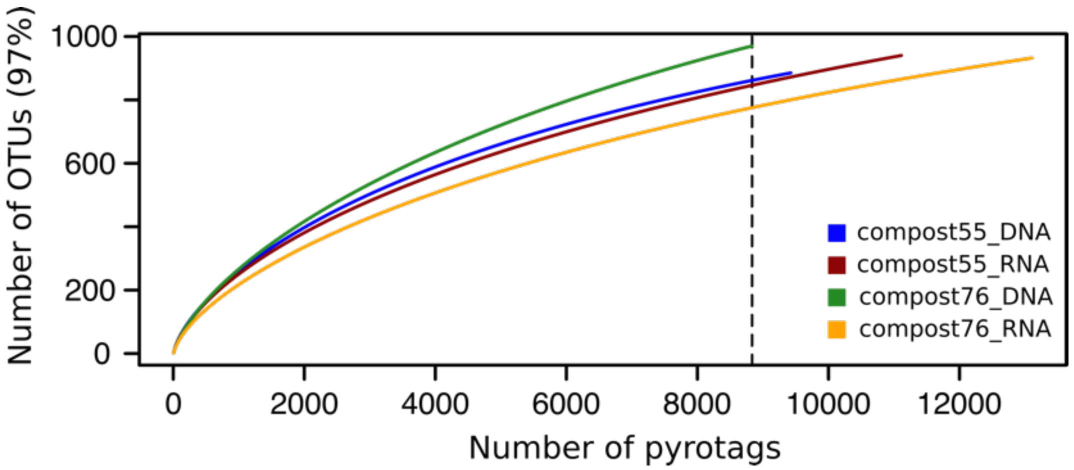


**Supplementary Figure S2.** Rarefaction curves of subsampled OTUs for 16S rRNA genes (DNA level) and transcripts (RNA level) in compost55 and compost76 at 97% similarity.

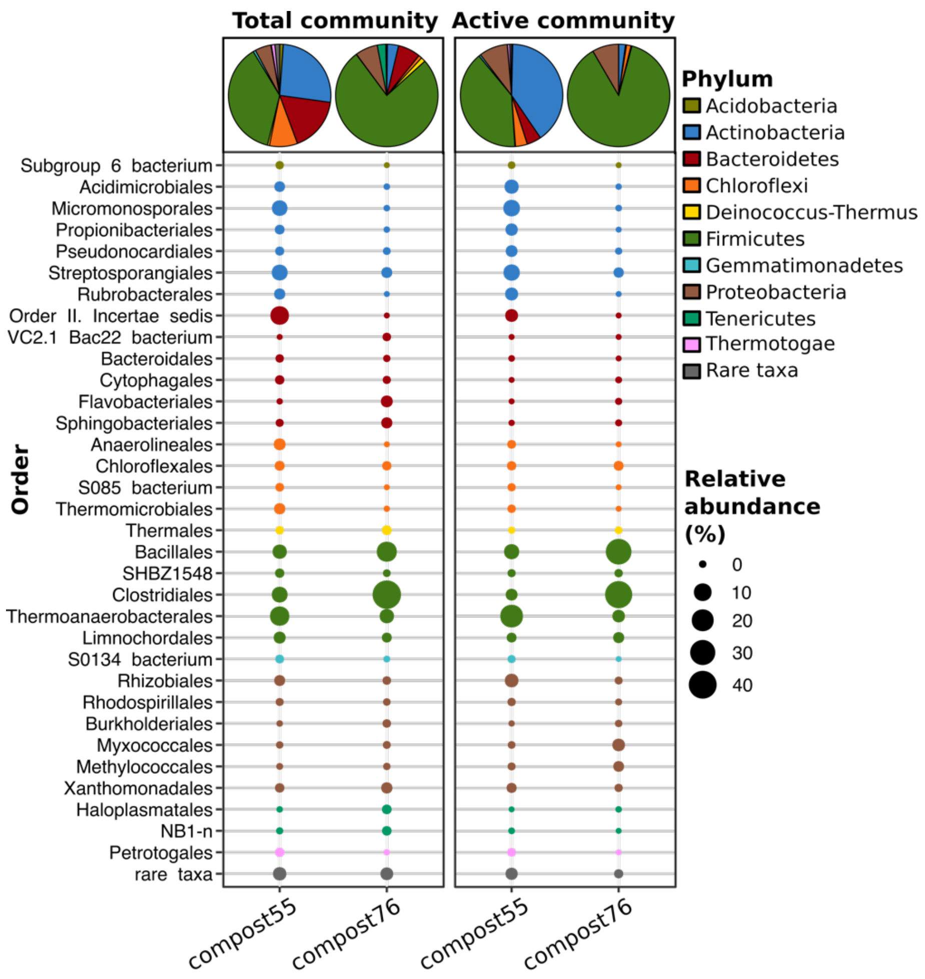


**Supplementary Figure S3.** Phylogenetic composition of bacterial communities in compost55 and compost76, revealed from 16S rRNA genes (DNA-level) and transcripts (RNA-level). Taxonomic specificity ranges from phylum level (pie chart) to order level (point chart) resolution when applicable. Taxonomic classification of DNA- or RNA-derived 16S rRNA gene sequences was performed according to SILVA SSU database 128 (Quast et al. 2013). Low relative abundant groups (<1% at phylum level or <0.5% at order level) were summarized as artificial group “rare taxa”.

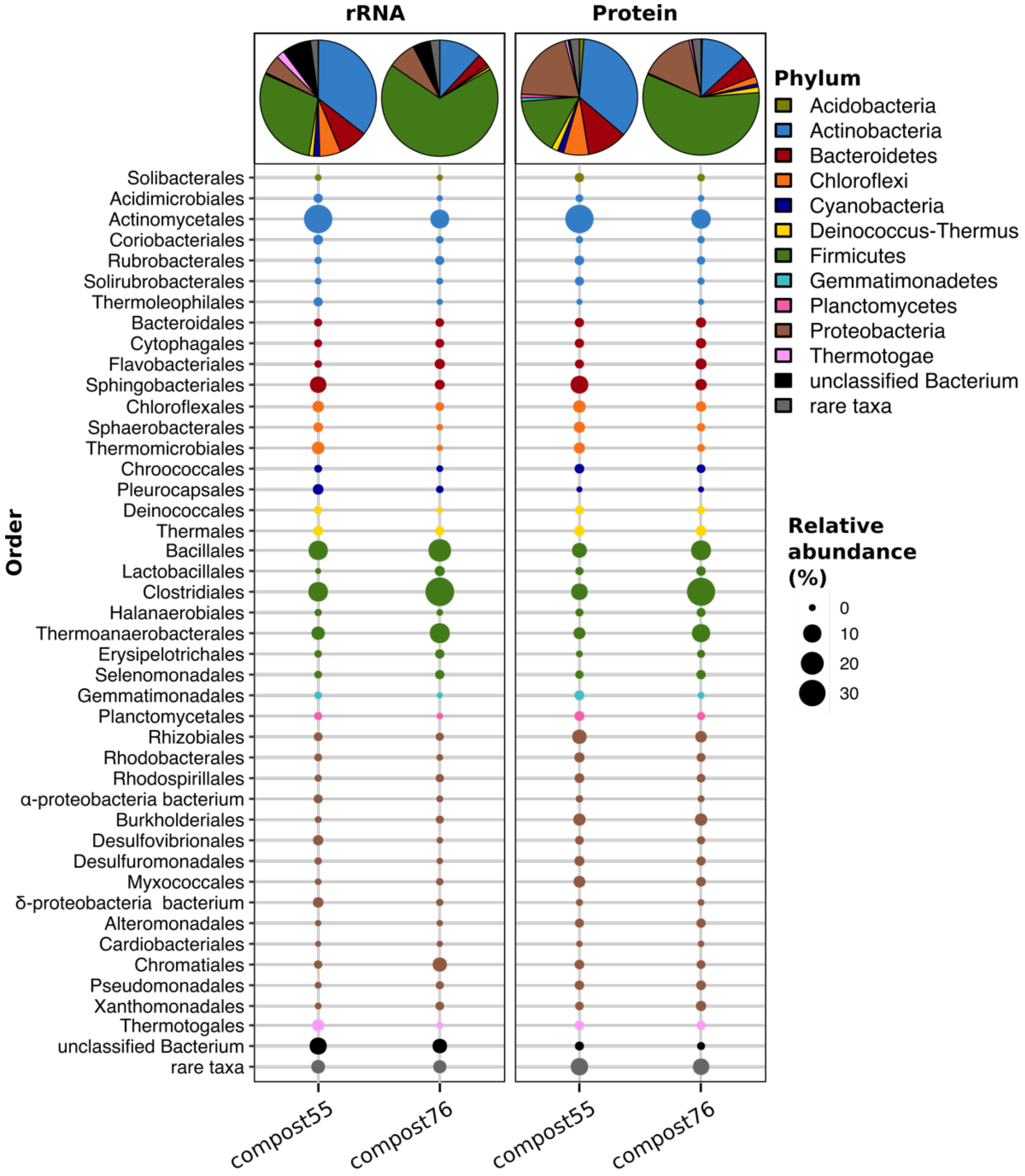


**Supplementary Figure S4.** Phylogenetic composition of bacterial communities in compost55 and compost76 as annotated by the MG-RAST platform (Keegan et al. 2016). Taxonomic specificity ranges from phylum level (pie chart) to order level (point chart) resolution when applicable. Microbial composition annotation was performed using MG-RAST best hit classification tool against the databases of M5RNA (Non-redundant multisource ribosomal RNA annotation) and M5NR (M5 non-redundant protein) available within MG-RAST with default settings. Low relative abundant groups (<1% at phylum level, <0.5% at order level) were summarized as artificial group "rare taxa".

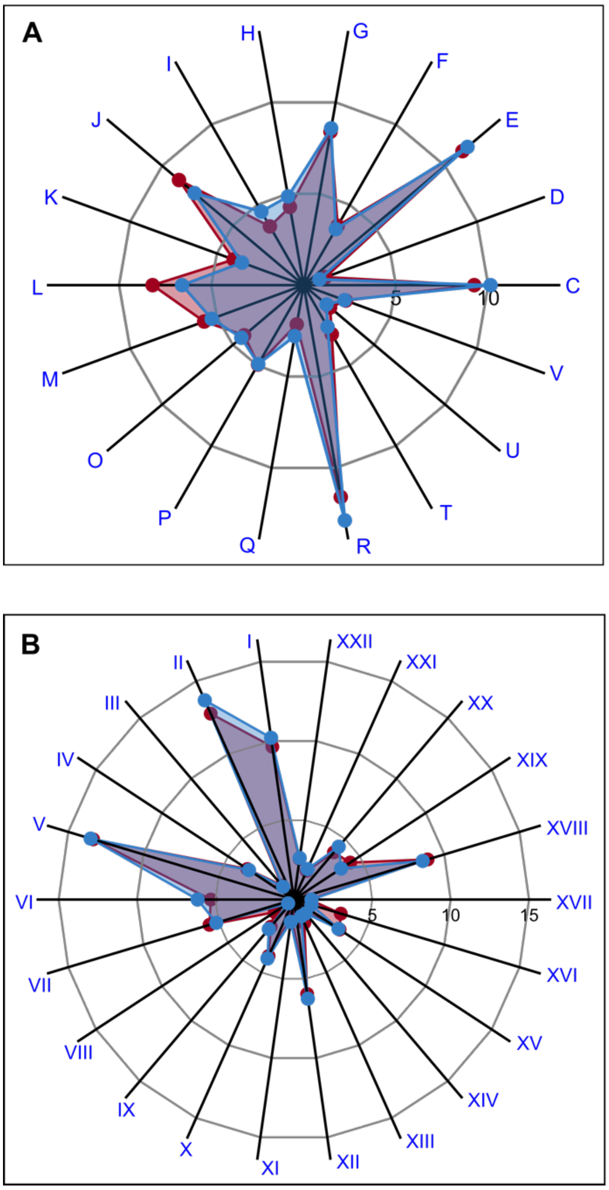


**COG catagory**

C-Energy production and conversion
D-Cell cycle control, cell division, chromosome partitioning
E-Amino acid transport and metabolism
F-Nucleotide transport and metabolism
G-Carbohydrate transport and metabolism
H-Coenzyme transport and metabolism
I-Lipid transport and metabolism
J-Translation, ribosomal structure and biogenesis
K-Transcription
L-Replication, recombination and repair
M-Cell wall/membrane/envelope biogenesis
O-Posttranslational modification, protein turnover, chaperones
P-Inorganic ion transport and metabolism
Q-Secondary metabolites biosynthesis, transport and catabolism
R-General function prediction only
T-Signal transduction mechanisms
U-Intracellular trafficking, secretion, and vesicular transport
V-Defense mechanisms

**Subsystems catagory**

I-Amino Acids and Derivatives
II-Carbohydrates
III-Cell Division and Cell Cycle
IV-Cell Wall and Capsule
V-Clustering-based subsystems
VI-Cofactors, Vitamins, Prosthetic Groups, Pigments
VII-DNA Metabolism
VIII-Dormancy and Sporulation
IX-Fatty Acids, Lipids, and Isoprenoids
X-Membrane Transport
XI-Metabolism of Aromatic Compounds
XII-Miscellaneous
XIII-Motility and Chemotaxis
XIV-Nitrogen Metabolism
XV-Nucleosides and Nucleotides
XVI-Phages, Prophages, Transposable elements, Plasmids
XVII-Phosphorus Metabolism
XVIII-Protein Metabolism
XIX-RNA Metabolism
XX-Respiration
XXI-Stress Response
XXII-Virulence, Disease and Defense

**Supplementary Figure S5**. Functional distribution pattern in compost55 (**blue** filled circle) and compost76 (**red** filled circle) microbial consortia. MG-RAST annotation against COG (**A**) and (**B**) subsystems database are shown. Only functional categories with relative abundance of more than 1 % were shown. The functional annotation of metagenomic reads was performed by MG-RAST pipeline (Keegan et al. 2016).

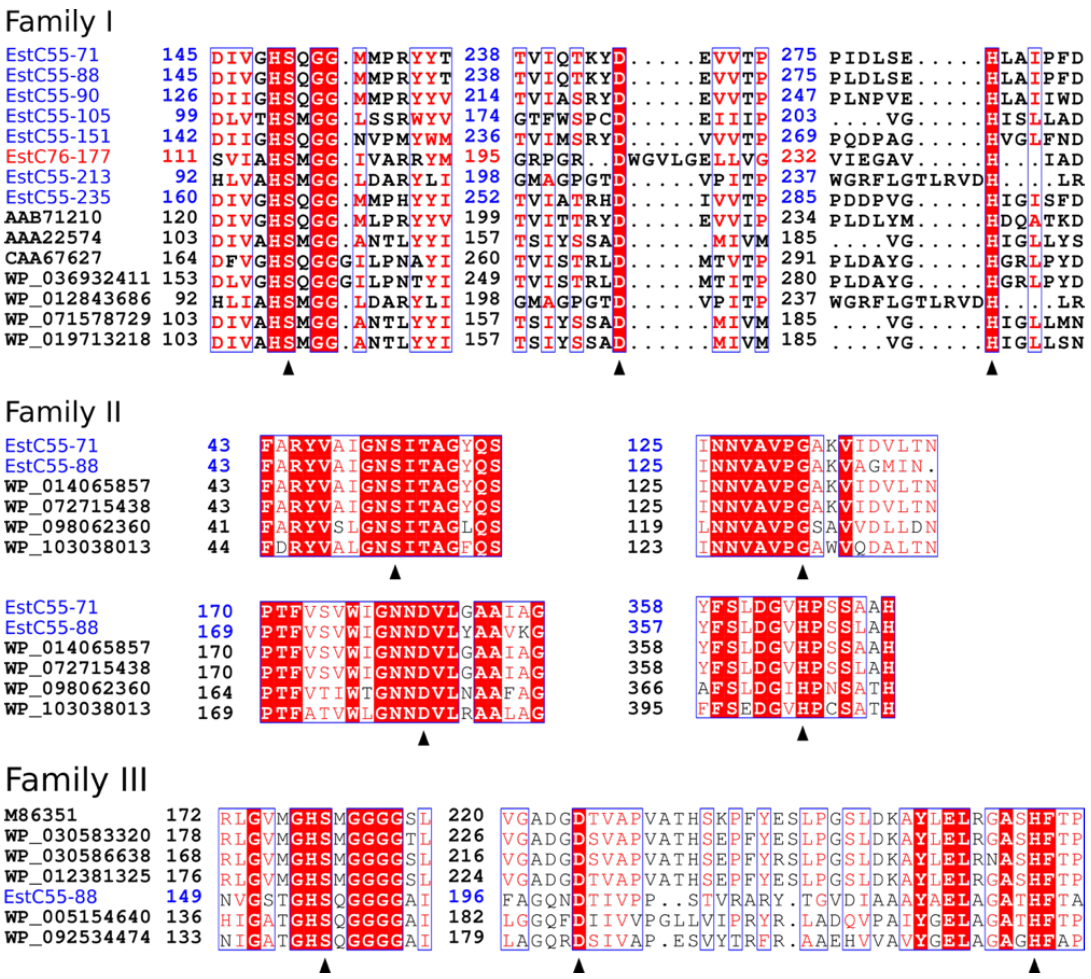
Family I
EstC55-71 145 DIVGHSQGG.MMPRYYT 238 TVIQTKYD.....EVVTP 275 PIDLSE.....HLAIPFD
EstC55-88 145 DIVGHSQGG.MMPRYYT 238 TVIQTKYD.....EVVTP 275 PLDLSE.....HLAIPFD
EstC55-90 126 DIIGHSQGG.MMPRYYV 214 TVIASRYD.....EVVTP 247 PLNPVE.....HLAIIWD
EstC55-105 99 DLVTHSMGG.LSSRWYV 174 GTFWSPCD.....EIIIP 203 ....VG.....HISLLAD
EstC55-151 142 DIIGHSQGG.NVPMYWM 236 TVIMSRYD.....VVVTP 269 PQDPAG.....HVGLFND
EstC76-177 111 SVIAHSMGG.IVARRYM 195 GRPGR..DWGVLGELLVG 232 VIEGAV.....H...IAD
EstC55-213 92 HLVAHSMGG.LDARYLI 198 GMAGPGTD.....VPITP 237 WGRFLGTLRVDH....LR
EstC55-235 160 DIVGHSQGG.MMPHYYI 252 TVIATRHD.....IVVTP 285 PDDPVG.....HIGISFD
AAB71210 120 DIVGHSQGG.MLPRYYV 199 TVITTRYD.....EVVTP 234 PLDLYM.....HDQATKD
AAA22574 103 DIVAHSMGG.ANTLYYI 157 TSIYSSAD.....MIVM 185 ....VG.....HIGLLYS
CAA67627 164 DFVGHSQGGGILPNAYI 260 TVISTRLD.....MTVTP 291 PLDAYG.....HGRLPYD
WP_036932411 153 DLVGHSQGGGILPNTYI 249 TVISTRLD.....MTITP 280 PLDAYG.....HGRLPYD
WP_012843686 92 HLIAHSMGG.LDARYLI 198 GMAGPGTD.....VPITP 237 WGRFLGTLRVDH....LR
WP_071578729 103 DIVAHSMGG.ANTLYYI 157 TSIYSSAD.....MIVM 185 ....VG.....HIGLLMN
WP_019713218 103 DIVAHSMGG.ANTLYYI 157 TSIYSSAD.....MIVM 185 ....VG.....HIGLLSN
Family II
EstC55-71 43 FARYVAIGNSITAGYQS 125 INNVAVPGAKVIDVLTN
EstC55-88 43 FARYVAIGNSITAGYQS 125 INNVAVPGAKVAGMIN.
WP_014065857 43 FARYVAIGNSITAGYQS 125 INNVAVPGAKVIDVLTN
WP_072715438 43 FARYVAIGNSITAGYQS 125 INNVAVPGAKVIDVLTN
WP_098062360 41 FARYVSLGNSITAGLQS 119 LNNVAVPGSAVVDLLDN
WP_103038013 44 FDRYVALGNSITAGFQS 123 INNVAVPGAWVQDALTN
EstC55-71 170 PTFVSVWIGNNDVLGAAIAG 358 YFSLDGVHPSSAAH
EstC55-88 169 PTFVSVWIGNNDVLYAAVKG 357 YFSLDGVHPSSLAH
WP_014065857 170 PTFVSVWIGNNDVLGAAIAG 358 YFSLDGVHPSSAAH
WP_072715438 170 PTFVSVWIGNNDVLGAAIAG 358 YFSLDGVHPSSLAH
WP_098062360 164 PTFVTIWTGNNDVLNAAFAG 366 AFSLDGIHPNSATH
WP_103038013 169 PTFATVWLGNNDVLRAALAG 395 FFSEDGVHPCSATH
Family III
M86351 172 RLGVMGHSMGGGGSL 220 VGADGDTVAPVATHSKPFYESLPGSLDKAYLELRGASHFTP
WP_030583320 178 RLGVMGHSMGGGGTL 226 VGADGDSVAPVATHSEPFYESLPGSLDKAYLELRGASHFTP
WP_030586638 168 RLGVMGHSMGGGGSL 216 VGADGDSVAPVATHSEPFYRSLPGSLDKAYLELRNASHFTP
WP_012381325 176 RLGVMGHSMGGGGSL 224 VGADGDTVAPVATHSEPFYESLPGSLDKAYLELRGASHFTP
EstC55-88 149 NVGSTGHSQGGGGAI 196 FAGQNDTIVPP..STVRARY.TGVDIAAAYAELAGATHFTA
WP_005154640 136 HIGATGHSQGGGGAI 182 LGGQFDIIVVPGLLVIPRYR.LADQVPAIYGELAGATHFTP
WP_092534474 133 NIGATGHSQGGGGAI 179 LAGQRDSIVAP.ESVYTRFR.AAEHVVAVYGELAGAGHFAP

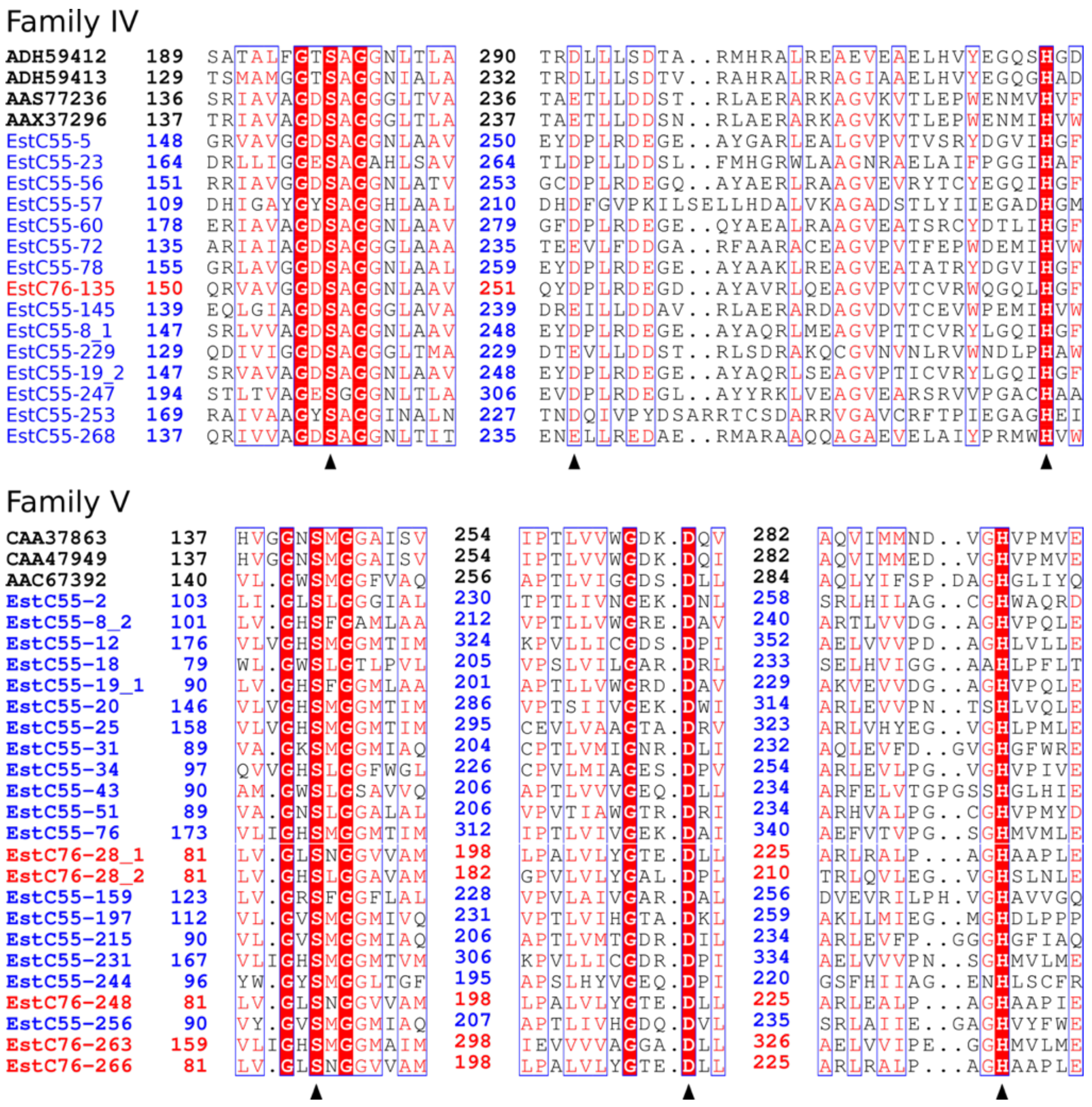
Family IV
ADH59412 189 SATALFGTSAGGNLTLA 290 TRDLLLSDTA..RMHRALREAEVEAELHVYEGQSHGD
ADH59413 129 TSMAMGGTSAGGNIALA 232 TRDLLLSDTV..RAHRALRRAGIAAELHVYEGQGHAD
AAS77236 136 SRIAVAGDSAGGGLTVA 236 TAETLLDDST..RLAERARKAGVKVTLEPWENMVHVF
AAX37296 137 TRIAVAGDSAGGGLTLA 237 TAETLLDDSN..RLAERARKAGVKVTLEPWENMIHVW
EstC55-5 148 GRVAVGGDSAGGNLAAV 250 EYDPLRDEGE..AYGARLEALGVPVTVSRYDGVIHGF
EstC55-23 164 DRLLIGGESAGAHLSAV 264 TLDPLLDDSL..FMHGRWLAAGNRAELAIFPGGIHAF
EstC55-56 151 RRIAVGGDSAGGNLATV 253 GCDPLRDEGQ..AYAERLRAAGVEVRYTCYEGQIHGF
EstC55-57 109 DHIGAYGYSAGGHLAAL 210 DHDFGVPKILSELLHDALVKAGADSTLYIIEGADHGM
EstC55-60 178 ERIAVAGDSAGGNLAAV 279 GFDPLRDEGE..QYAEALRAAGVEATSRCYDTLIHGF
EstC55-72 135 ARIAIAGDSAGGGLAAA 235 TEEVLFDDGA..RFAARACEAGVPVTFEPWDEMIHVW
EstC55-78 155 GRLAVGGDSAGGNLAAL 259 EYDPLRDEGE..AYAAKLREAGVEATATRYDGVIHGF
EstC76-135 150 QRVAVGGDSAGGNLAAV 251 QYDPLRDEGD..AYAVRLQEAGVPVTCVRWQGQLHGF
EstC55-145 139 EQLGIAGDSAGGGLAVA 239 DREILLDDAV..RLAERARDAGVDVTCEVWPEMIHVW
EstC55-8_1 147 SRLVVAGDSAGGNLAAV 248 EYDPLRDEGE..AYAQRLMEAGVPTTCVRYLGQIHGF
EstC55-229 129 QDIVIGGDSAGGGLTMA 229 DTEVLLDDST..RLSDRAKQCGVNVNLRVWNDLPHAW
EstC55-19_2 147 SRVAVAGDSAGGNLAAV 248 EYDPLRDEGE..AYAQRLSEAGVPTICVRYLGQIHGF
EstC55-247 194 STLTVAGESGGGNLTLA 306 EVDPLRDEGL..AYYRKLVEAGVEARSRVVPGACHAA
EstC55-253 169 RAIVAAGYSAGGINALN 227 TNDQIVPYDSARRTCSDARRVGAVCRFTPIEGAGHEI
EstC55-268 137 QRIVVAGDSAGGNLTIT 235 ENELLREDAE..RMARAAQQAGAEVELAIYPRMWHVW
Family V
CAA37863 137 HVGGNSMGGAISV 254 IPTLVVWGDK.DQV 282 AQVIMMND..VGHVPMVE
CAA47949 137 HVGGNSMGGAISV 254 IPTLVVWGDK.DQI 282 AQVIMMED..VGHVPMVE
AAC67392 140 VL.GWSMGGFVAQ 256 APTLVIGGDS.DLL 284 AQLYIFSP.DAGHGLIYQ
EstC55-2 103 LI.GLSLGGGIAL 230 TPTLIVNGEK.DNL 258 SRLHILAG..CGHWAQRD
EstC55-8_2 101 LV.GHSFGAMLAA 212 VPTLLVWGRE.DAV 240 ARTLVVDG..AGHVPQLE
EstC55-12 176 VLVGHSMGGMTIM 324 KPVLLICGDS.DPI 352 AELVVVPD..AGHLVLLE
EstC55-18 79 WL.GWSLGTLPVL 205 VPSLVILGAR.DRL 233 SELHVIGG..AAHLPFLT
EstC55-19_1 90 LV.GHSFGGMLAA 201 APTLLVWGRD.DAV 229 AKVEVVDG..AGHVPQLE
EstC55-20 146 VLVGHSMGGMTIM 286 VPTSIIVGEK.DWI 314 ARLEVVPN..TSHLVQLE
EstC55-25 158 VLVGHSMGGMTIM 295 CEVLVAAGTA.DRV 323 ARLVHYEG..VGHLPMLE
EstC55-31 89 VA.GKSMGGMIAQ 204 CPTLVMIGNR.DLI 232 AQLEVFD..GVGHGFWRE
EstC55-34 97 QVVGHSLGGFWGL 226 CPVLMIAGES.DPV 254 ARLEVLPG..VGHVPIVE
EstC55-43 90 AM.GWSLGSAVVQ 206 APTLVVVGEQ.DLL 234 ARFELVTGPGSSHGLHIE
EstC55-51 89 VA.GNSLGGALAL 206 VPVTIAWGTR.DRI 234 ARHVALPG..CGHVPMYD
EstC55-76 173 VLIGHSMGGMTIM 312 IPTLVIVGEK.DAI 340 AEFVTVPG..SGHMVMLE
EstC76-28_1 81 LV.GLSNGGVVAM 198 LPALVLYGTE.DLL 225 ARLRALP...AGHAAPLE
EstC76-28_2 81 LV.GHSLGGAVAM 182 GPVLVLYGAL.DPL 210 TRLQVLEG..VGHSLNLE
EstC55-159 123 LV.GRSFGGFLAL 228 VPVLAIVGAR.DAL 256 DVEVRILPH.VGHAVVGQ
EstC55-197 112 VL.GVSMGGMIVQ 231 VPTLVIHGTA.DKL 259 AKLLMIEG..MGHDLPPP
EstC55-215 90 VL.GVSMGGMIAQ 206 APTLVMTGDR.DIL 234 ARLEVFP..GGGHGFIAQ
EstC55-231 167 VLIGHSMGGMTVM 306 KPVLLICGDR.DPI 334 AELVVVPN..SGHMVLME
EstC55-244 96 YW.GYSMGGLTGF 195 APSLHYVGEQ.DPI 220 GSFHIIAG..ENHLSCFR
EstC76-248 81 LV.GLSNGGVVAM 198 LPALVLYGTE.DLL 225 ARLEALP...AGHAAPIE
EstC55-256 90 VY.GVSMGGMIAQ 207 APTLIVHGDQ.DVL 235 SRLAIIE..GAGHVYFWE
EstC76-263 159 VLIGHSMGGMAIM 298 IEVVVVAGGA.DLL 326 AELVVIPE..GGHMVLME
EstC76-266 81 LV.GLSNGGVVAM 198 LPALVLYGTE.DLL 225 ARLRALP...AGHAAPLE

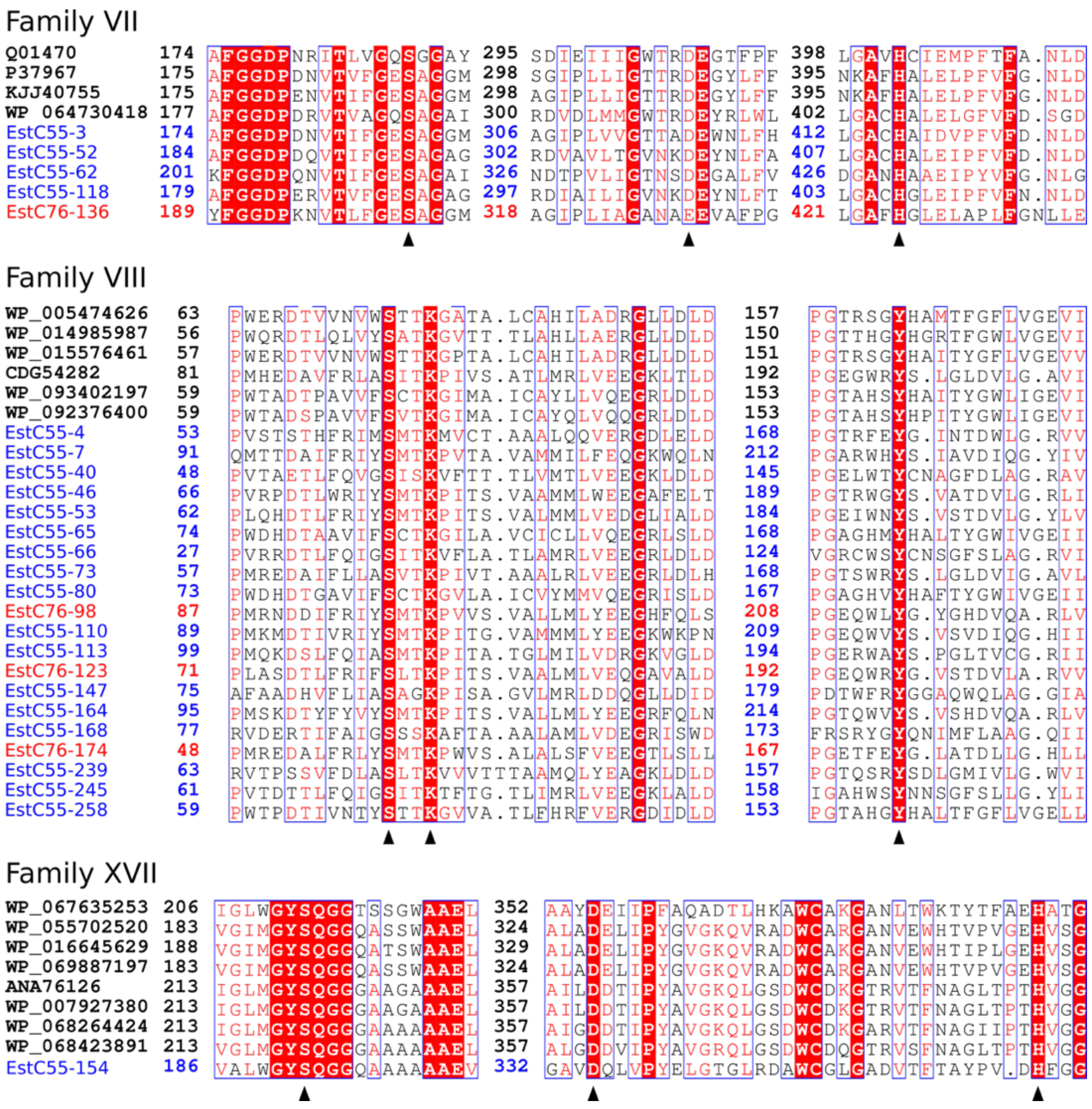
Family VII
Q01470 174 AFGGDPNRITLVGQSGGAY 295 SDIEIIIGWTRDEGTFPF 398 LGAVHCIEMPFTFA.NLD
P37967 175 AFGGDPDNVTVFGESAGGM 298 SGIPLLIGTTRDEGYLFF 395 NKAFHALELPFVFG.NLD
KJJ40755 175 AFGGDPENVTIFGESAGGM 298 AGIPLLIGTTRDEGYLFF 395 NKAFHALELPFVFG.NLD
WP 064730418 177 AFGGDPDRVTVAGQSAGAI 300 RDVDLMMGWTRDEYRLWL 402 LGACHALELGFVFD.SGD
EstC55-3 174 AFGGDPDNVTIFGESAGGM 306 AGIPLVVGTTADEWNLFH 412 LGACHAIDVPFVFD.NLD
EstC55-52 184 AFGGDPDQVTIFGESAGAG 302 RDVAVLTGVNKDEYNLFA 407 LGACHALEIPFVFD.NLD
EstC55-62 201 KFGGDPQNVTIFGESAGAI 326 NDTPVLIGTNSDEGALFV 426 DGANHAAEIPYVFG.NLG
EstC55-118 179 AFGGDPERVTVFGESAGAG 297 RDIAILIGVNKDEYNLFT 403 LGACHGLEIPFVFN.NLD
EstC76-136 189 YFGGDPKNVTLFGESAGGM 318 AGIPLIAGANAEEVAFPG 421 LGAFHGLELAPLFGNLLE
Family VIII
WP_005474626 63 PWERDTVVNVWSTTKGATA.LCAHILADRGLLDLD 157 PGTRSGYHAMTFGFLVGEVI
WP_014985987 56 PWQRDTLQLVYSATKGVTT.TLAHLLAERGLLDLD 150 PGTTHGYHGRTFGWLVGEVI
WP_015576461 57 PWERDTVVNVWSTTKGPTA.LCAHILADRGLLDLD 151 PGTRSGYHAITYGFLVGEVV
CDG54282 81 PMHEDAVFRLASITKPIVS.ATLMRLVEEGKLTLD 192 PGEGWRYS.LGLDVLG.AVI
WP_093402197 59 PWTADTPAVVFSCTKGIMA.ICAYLLVQEGRLDLD 153 PGTAHSYHAITYGWLIGEVI
WP_092376400 59 PWTADSPAVVFSVTKGIMA.ICAYQLVQQGRLDLD 153 PGTAHSYHPITYGWLIGEVI
EstC55-4 53 PVSTSTHFRIMSMTKMVCT.AAALQQVERGDLELD 168 PGTRFEYG.INTDWLG.RVV
EstC55-7 91 QMTTDAIFRIYSMTKPVTA.VAMMILFEQGKWQLN 212 PGARWHYS.IAVDIQG.YIV
EstC55-40 48 PVTAETLFQVGSISKVFTT.TLVMTLVEEGKLDLD 145 PGELWTYCNAGFDLAG.RAV
EstC55-46 66 PVRPDTLWRIYSMTKPITS.VAAMMLWEEGAFELT 189 PGTRWGYS.VATDVLG.RLI
EstC55-53 62 PLQHDTLFRIYSMTKPITS.VALMMLVEDGLIALD 184 PGEIWNYS.VSTDVLG.YLV
EstC55-65 74 PWDHDTAAVIFSCTKGILA.VCICLLVQEGRLSLD 168 PGAGHMYHALTYGWIVGEII
EstC55-66 27 PVRRDTLFQIGSITKVFLA.TLAMRLVEEGRLDLD 124 VGRCWSYCNSGFSLAG.RVI
EstC55-73 57 PMREDAIFLLASVTKPIVT.AAALRLVEEGRLDLH 168 PGTSWRYS.LGLDVIG.AVL
EstC55-80 73 PWDHDTGAVIFSCTKGVLA.ICVYMMVQEGRISLD 167 PGAGHVYHAFTYGWIVGEII
EstC76-98 87 PMRNDDIFRIYSMTKPVVS.VALLMLYEEGHFQLS 208 PGEQWLYG.YGHDVQA.RLV
EstC55-110 89 PMKMDTIVRIYSMTKPITG.VAMMMLYEEGKWKPN 209 PGEQWVYS.VSVDIQG.HII
EstC55-113 99 PMQKDSLFQIASMTKPITA.TGLMILVDRGKVGLD 194 PGERWAYS.PGLTVCG.RII
EstC76-123 71 PLASDTLFRIFSLTKPITS.VAALMLVEQGAVALD 192 PGEQWRYG.VSTDVLA.RVV
EstC55-147 75 AFAADHVFLIASAGKPISA.GVLMRLDDQGLLDID 179 PDTWFRYGGAQWQLAG.GIA
EstC55-164 95 PMSKDTYFYVYSMTKPITS.VALLMLYEEGRFQLN 214 PGTQWVYS.VSHDVQA.RLV
EstC55-168 77 RVDERTIFAIGSSSKAFTA.AALAMLVDEGRISWD 173 FRSRYGYQNIMFLAAG.QII
EstC76-174 48 PMREDALFRLYSMTKPWVS.ALALSFVEEGTLSLL 167 PGETFEYG.LATDLLG.HLL
EstC55-239 63 RVTPSSVFDLASLTKVVVTTTAAMQLYEAGKLDLD 157 PGTQSRYSDLGMIVLG.WVI
EstC55-245 61 PVTDTTLFQIGSITKTFTG.TLIMRLVEEGKLALD 158 IGAHWSYNNSGFSLLG.YLI
EstC55-258 59 PWTPDTIVNTYSTTKGVVA.TLFHRFVERGDIDLD 153 PGTAHGYHALTFGFLVGELL
Family XVII
WP_067635253 206 IGLWGYSQGGTSSGWAAEL 352 AAYDEIIPFAQADTLHKAWCAKGANLTWKTYTFAEHATG
WP_055702520 183 VGIMGYSQGGQASSWAAEL 324 ALADELIPYGVGKQVRADWCARGANVEWHTVPVGEHVSG
WP_016645629 188 VGIMGYSQGGQATSWAAEL 329 ALADELIPYGVGKQVRADWCAKGANVEWHTIPLGEHVSG
WP_069887197 183 VGIMGYSQGGQASSWAAEL 324 ALADELIPYGVGKQVRADWCARGANVEWHTVPVGEHVSG
ANA76126 213 IGLMGYSQGGGAAGAAAEL 357 AILDDTIPYAVGKQLGSDWCDKGTRVTFNAGLTPTHVGG
WP_007927380 213 IGLMGYSQGGGAAGAAAEL 357 AILDDTIPYAVGKQLGSDWCDKGTRVTFNAGLTPTHVGG
WP_068264424 213 IGLMGYSQGGGAAAAAAEL 357 AIGDDTIPYAVGKQLGSDWCDKGARVTFNAGIIPTHVGG
WP_068423891 213 VGLMGYSQGGGAAAAAAEL 357 ALGDDVIPYAVGRQLGSDWCDQGTRVSFNAGLTPTHVGG
EstC55-154 186 VALWGYSQGGQAAAAAAEV 332 GAVDQLVPYELGTGLRDAWCGLGADVTFTAYPV.DHFGG

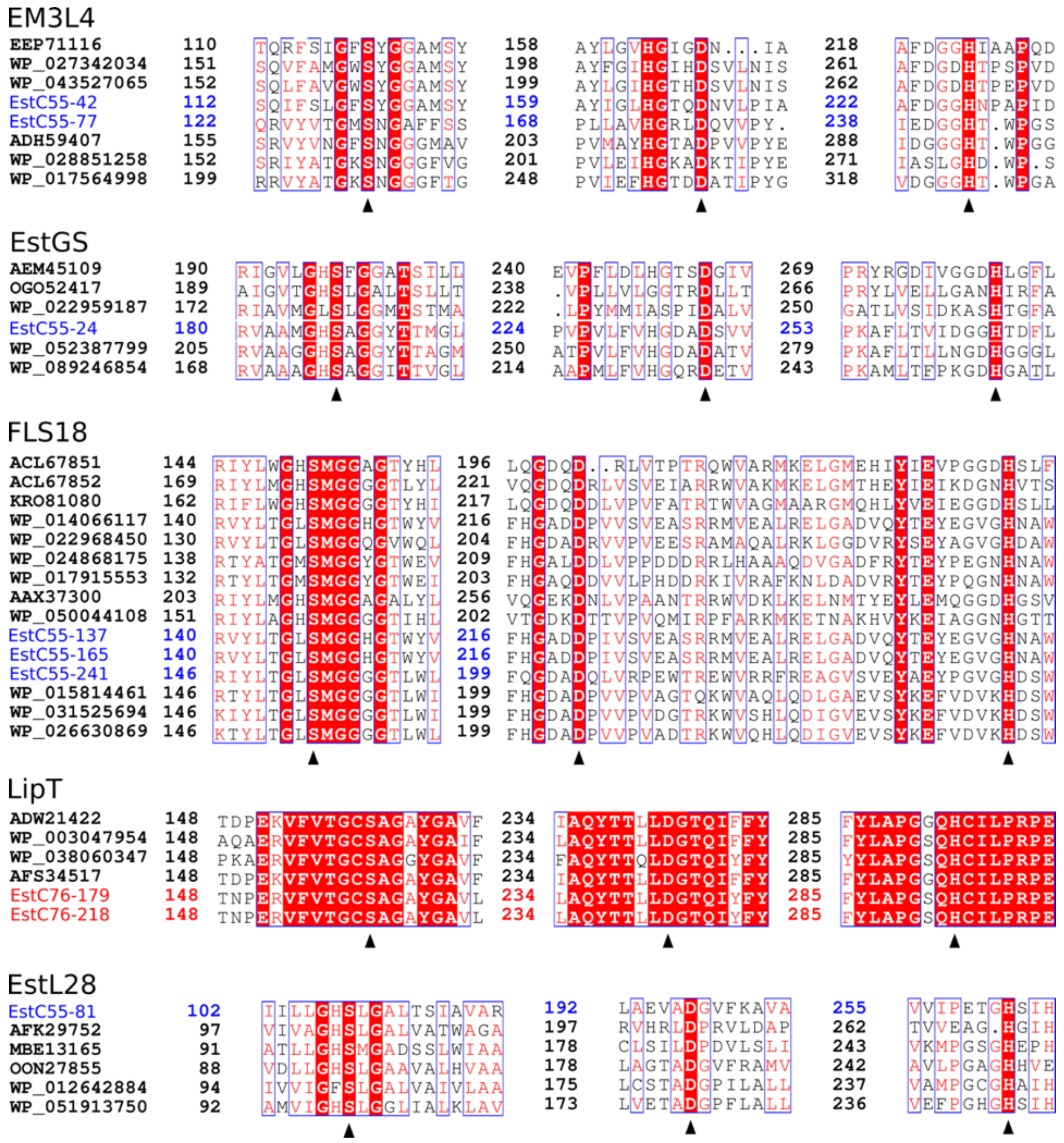
EM3L4
EstGS
FLS18
LipT
EstL28

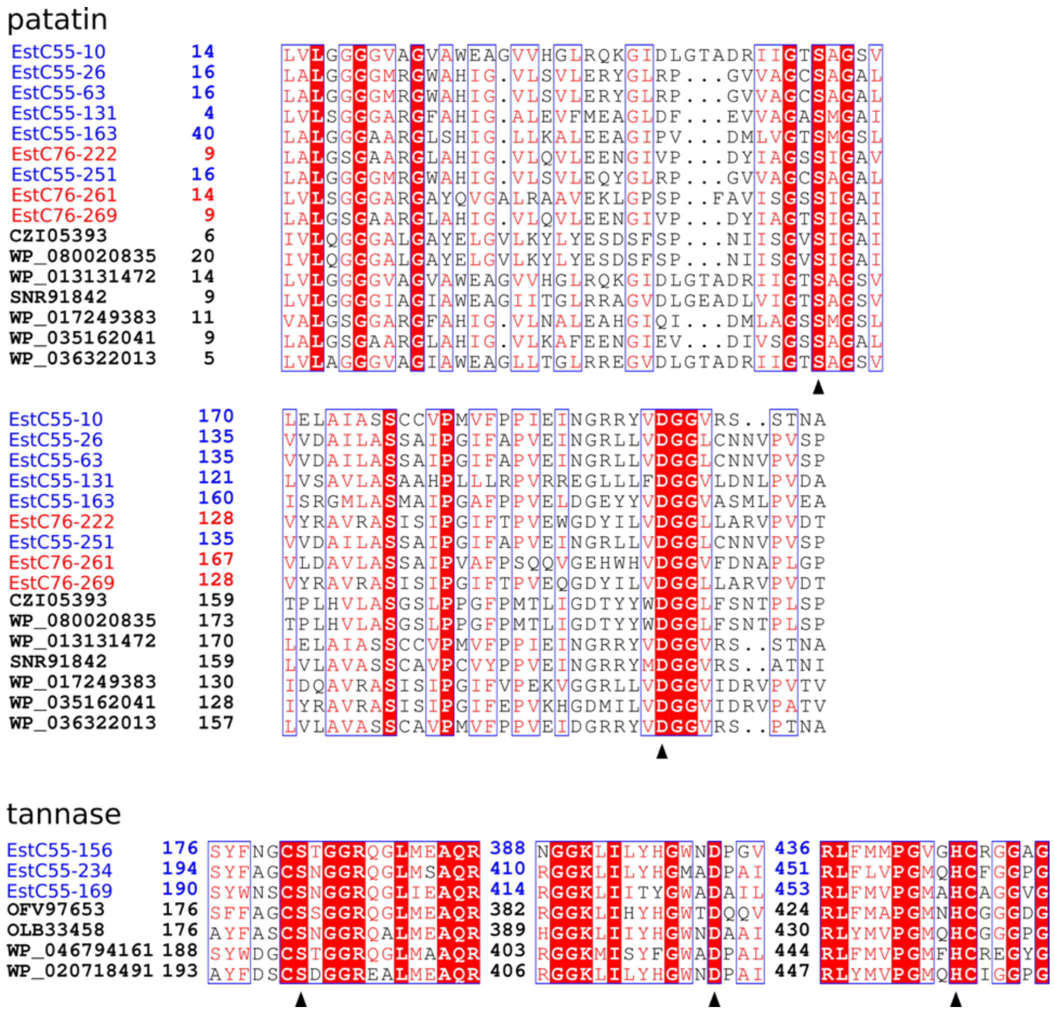


**Supplementary Figure S6a.** Multiple sequence alignments of partial amino acid sequences harboring homologous catalytic regions. Lipolytic enzymes were from reported families. Residues, which are partially consistent, are in frames. Identical residues are shaded in red. Triangles underneath residues indicate the catalytic triad. The functionally identified LEs were assigned to known lipolytic families. **Family I**: EstC55-71, EstC55-88, EstC55-90, EstC55-105, EstC55-151, EstC55-213 and EstC55-235, functionally derived LEs from sample compost55 (this study); EstC76-177, functionally derived LE from sample compost76 (this study); AAB71210, lipase LipA from *Streptomyces cinnamoneus*; AAA22574, lipase from *Bacillus subtilis*; CAA67627, triacylglycerol lipase from *Cutibacterium acnes*; WP_036932411, triacylglycerol lipase from *Cutibacterium avidum*; WP_012843686, alpha/beta fold hydrolase from *Rhodothermus marinus*; WP_071578729, triacylglycerol lipase from *Bacillus* sp. FMQ74; WP_019713218, triacylglycerol lipase from *Bacillus subtilis*. **Family II**: EstC55-111 and EstC55-150, functionally derived LEs from sample compost55 (this study); WP_014065857, SGNH/GDSL hydrolase family protein from *Rhodothermus marinus*; WP_072715438, hypothetical protein from *Rhodothermus profundi*; WP_098062360, hypothetical protein from *Longimonas halophila*; WP_103038013, SGNH/GDSL hydrolase family protein from *Salinivenus iranica*. **Family III**: EstC55-95, functionally derived LE from sample compost55 (this study); M86351, triacylglycerol acylhydrolase from *Streptomyces* sp.; WP_030583320, lipase from *Streptomyces globisporus*; WP_030586638, lipase from *Streptomyces anulatus*; WP_012381325, alpha/beta hydrolase from *Streptomyces*; WP_005154640, lipase from *Amycolatopsis azurea*; WP_092534474, acetylxylan esterase from *Yuhushiella deserti*.

**Family IV**: EstC55-5, EstC55-23, EstC55-56, EstC55-57, EstC55-60, EstC55-72, EstC55-78, EstC55-145, EstC55-8_1, EstC55-229, EstC55-19_2, EstC55-247, EstC55-253 and EstC55-268, functionally derived LEs from sample compost55 (this study); EstC76-135, F functionally derived LE from sample compost76 (this study); ADH59412, esterase from uncultured bacterium; ADH59413, esterase from uncultured bacterium; AAS77236, lipase/esterase from uncultured bacterium; AAX37296, lipase/esterase from uncultured bacterium. **Family V**: EstC55-2, EstC55-8_2, EstC55-12, EstC55-18, EstC55-19_1, EstC55-20, EstC55-25, EstC55-31, EstC55-34, EstC55-43, EstC55-51, EstC55-76, EstC55-159, EstC55-197, EstC55-215, EstC55-231, EstC55-244 and EstC55-256, functionally derived LEs from sample compost55 (this study); EstC76-28_1, EstC76-28_2, EstC76-248, EstC76-263 and EstC76-266, functionally derived LEs from sample compost76 (this study); CAA37863, triacylglycerol lipase from *Moraxella* sp.; CAA47949, triacylglycerol lipase from *Psychrobacter immobilis*; AAC67392, lipolytic enzyme from *Sulfolobus acidocaldarius*. **Family VII**: EstC55-3, EstC55-52, EstC55-62 and EstC55-118, functionally derived LEs from sample compost55 (this study); EstC76-136, functionally derived LE from sample compost76 (this study); Q01470, serine esterase from *Pseudarthrobacter oxydans*; P37967, para-nitrobenzyl esterase from *Bacillus subtilis*; KJJ40755, para-nitrobenzyl esterase from *Bacillus subtilis*; WP_064730418, carboxylesterase from *Streptomyces parvulus*. **Family VIII**: EstC55-4, EstC55-7, EstC55-40, EstC55-46, EstC55-53, EstC55-65, EstC55-66, EstC55-73, EstC55-80, EstC55-110, EstC55-113, EstC55-147, EstC55-164, EstC55-168, EstC55-239, EstC55-245 and EstC55-258, functionally derived LEs from sample compost55 (this study); EstC76-98, EstC76-123 and EstC76-174, functionally derived LEs from sample compost76 (this study); WP_005474626, esterase from *Streptomyces bottropensis*; WP_014985987, esterase from *Nocardia brasiliensis*; WP_015576461, esterase from *Streptomyces*; CDG54282, esterase EstB from *Halomonas* sp. A3H3; WP_093402197, carboxylesterase from *Verrucosispora sediminis*; WP_092376400, carboxylesterase from *Xiangella phaseoli*. **Family XVII**: EstC55-154, functionally derived LE from sample compost55 (this study); WP_067635253, triacylglycerol lipase from *Actinomadura latina*; WP_055702520, lipase from *Streptomyces silaceus*; WP_016645629, inactive lipase from *Streptomyces aurantiacus*; WP_069887197, lipase from *Streptomyces luteocolor*; ANA76126, secretory lipase LipJ2 from *Janibacter* sp. R02; WP_007927380, secretory lipase from *Janibacter hoylei*; WP_068264424, lipase from *Janibacter limosus*; WP_068423891, lipase from *Janibacter terrae*. **EM3L4**: EstC55-42 and EstC55-77, functionally derived LEs from sample compost55 (this study); EEP71116, ferruloyl esterase fee1B from *Micromonospora* sp. ATCC 39149; WP_027342034, cellulose-binding protein from *Hamadaea tsunoensis*; WP_043527065, cellulose-binding protein from *Actinoplanes utahensis*; ADH59407, esterase/lipase from uncultured bacterium; WP_028851258, hypothetical protein from *Thermocrispum municipal*; WP_017564998, hypothetical protein from *Nocardiopsis synnemataformans*. **EstGS**: EstC55-24, functionally derived LE from sample compost55 (this study); AEM45109, hypothetical protein from uncultured organism; OGO52417, hypothetical protein from *Chloroflexi* bacterium; WP_022959187, alpha/beta hydrolase from *Spongiibacter tropicus*; WP_052387799, alpha/beta hydrolase from *Dactylosporangium aurantiacum*; WP_089246854, chlorophyllase from *Asanoa hainanensis*. **FLS18**: EstC55-137, EstC55-165 and EstC55-241, functionally derived LEs from sample compost55 (this study); ACL67851, esterase/lipase from uncultured bacterium FLS18; ACL67852, esterase/lipase from uncultured bacterium FLS18; KRO81080, hypothetical protein from OM182 bacterium; WP_014066117, phospholipase from *Rhodothermus marinus*; WP_022968450, phospholipase from *Arenimonas oryziterrae*; WP_024868175, phospholipase from *Pseudoxanthomonas suwonensis*; WP_017915553, phospholipase from *Xanthomonas* sp. SHU 308; AAX37300, lipase/esterase from uncultured bacterium; WP_050044108, alpha/beta hydrolase from *Verrucomicrobia* bacterium SCGC AAA168-F10; WP_015814461, phospholipase/carboxylesterase from *Dyadobacter fermentans*; WP_031525694, phospholipase/carboxylesterase from *Dyadobacter crusticola*; WP_026630869, phospholipase/carboxylesterase from *Dyadobacter alkalitolerans*. **LipT**: EstC76-179 and EstC76-218, functionally derived LEs from sample compost76 (this study); ADW21422, putative esterase from *Thermus scotoductus* SA-01; WP_003047954, esterase from *Thermus aquaticus*; WP_038060347, esterase from *Thermus filiformis*; AFS34517, LipT from uncultured bacterium. EstL28: EstC55-81, functionally derived esterase from sample compost55 (this study); AFK29752, esterase from uncultured bacterium; MBE13165, hypothetical protein from *Chloroflexi* bacterium; OON27855, hypothetical protein from *Micromonospora* sp. Rc5; WP_012642884, alpha/beta hydrolase from *Thermomicrobium roseum*; WP_051913750, alpha/beta hydrolase from *Thermorudis peleae*; **Patatin-like-protein**: EstC55-10, EstC55-26, EstC55-63, EstC55-131, EstC55-163 and EstC55-251, functionally derived LEs from sample compost55 (this study); EstC76-222, EstC76-261 and EstC76-269 functionally derived LEs from sample compost76 (this study); CZI05393, patatin from *Legionella pneumophila*; WP_080020835, patatin-like phospholipase family protein from *Legionella pneumophila*; WP_013131472, patatin from *Thermobispora bispora*; SNR91842, NTE family protein from *Streptosporangium subroseum*; WP_017249383, esterase from *Brevibacillus brevis*; WP_035162041, esterase from *Caloranaerobacter azorensis*; WP_036322013, patatin-like phospholipase family protein from *Microbispora* sp.. **Tannase**: EstC55-156, EstC55-234 and EstC55-269, functionally derived LEs from sample compost55 (this study); OFV97653, hypothetical protein from *Acidobacteria* bacterium; OLB33458, feruloyl esterase from *Acidobacteria bacterium*; WP_046794161, tannase/feruloyl esterase family alpha/beta hydrolase from *Rhizobium* sp.; WP_020718491, tannase/feruloyl esterase family alpha/beta hydrolase from *Acidobacteriaceae* bacterium.

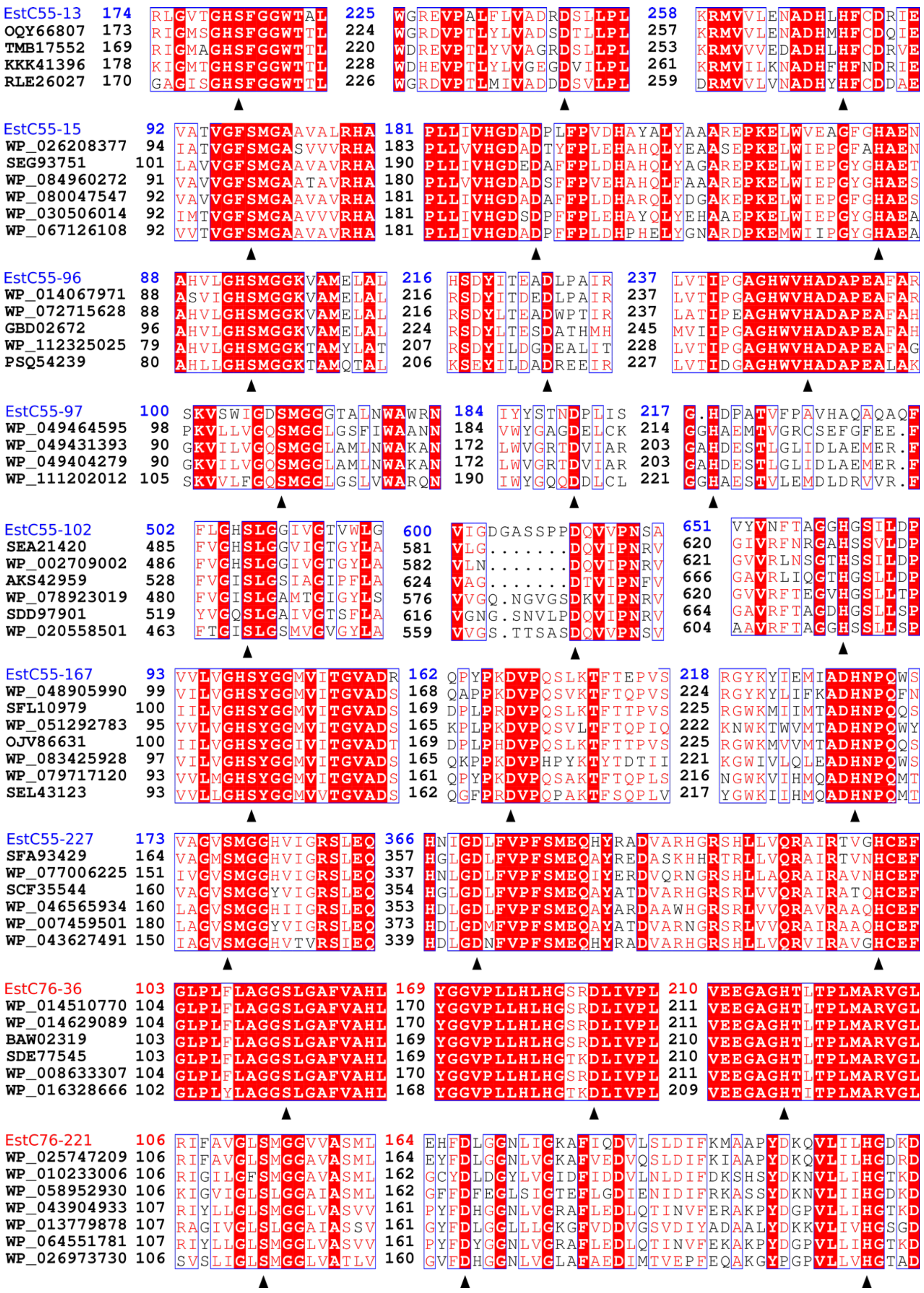


**Supplementary Figure S6b.** Multiple sequence alignments of partial amino acid sequences harboring homologous catalytic regions. Lipolytic enzymes were from putative novel families identified in this study. Residues, which are partially consistent, are in frames. Identical residues are shaded in red. Triangles underneath residues indicate the catalytic triad. **Putative new family 1**: EstC55-13, functionally derived LE from sample compost55 (this study); OQY66807, hypothetical protein from *Polyangiaceae* bacterium UTPRO1; TMB17552, hypothetical protein from *Deltaproteobacteria*

bacterium; KKK41396, Alpha/beta hydrolase family protein from *Lokiarchaeum* sp. GC14_75; RLE26027, hypothetical protein from *Actinobacteria bacterium*. **Putative new family 2**: EstC55-15, functionally derived LE from sample compost55 (this study); WP_026208377, hydrolase from *Catelliglobosispora koreensis*; SEG93751, Alpha/beta hydrolase family protein from *Nonomuraea solani*; WP_084960272,alpha/beta hydrolase from *Thermoactinospora rubra*; WP_080047547, alpha/beta hydrolase from *Nonomuraea* sp. ATCC 55076; WP_030506014, alpha/beta hydrolase from *Microbispora rosea*; WP_067126108, hydrolase from *Microtetraspora malaysiensis*. **Putative new family 3**: EstC55-96, functionally derived LE from sample compost55 (this study); WP_014067971, alpha/beta fold hydrolase from *Rhodothermus marinus*; WP_072715628, alpha/beta fold hydrolase from *Rhodothermus profundi*; GBD02672, Esterase YbfF from *bacterium* HR18; WP_112325025, alpha/beta fold hydrolase from *Rhodothermaceae* bacterium; PSQ54239, alpha/beta hydrolase from *Bacteroidetes* bacterium QH_10_64_37. **Putative new family 4**: EstC55-97, functionally derived LE from sample compost55 (this study); WP_049464595, alpha/beta hydrolase from *Stenotrophomonas maltophilia*; WP_049431393, alpha/beta hydrolase from *Stenotrophomonas maltophilia*; WP_049404279, alpha/beta hydrolase from *Stenotrophomonas maltophilia*; WP_111202012.1, hypothetical protein from *Stenotrophomonas maltophilia*. **Putative new family 5**: EstC55-102, functionally derived LE from sample compost55 (this study); SEA21420, alpha/beta hydrolase family protein from *Thiothrix caldifontis*; WP_002709002, lipase from *Thiothrix nivea*; AKS42959, Extracellular lipase, Pla-1/cef family from *Wenzhouxiangella marina*; WP_078923019, lipase from *Thiothrix eikelboomii*; SDD97901, Alpha/beta hydrolase family protein from *Aquimonas voraii*; WP_020558501, hypothetical protein from *Thiothrix flexilis*. **Putative new family 6**: EstC55-167, functionally derived LE from sample compost55 (this study); WP_048905990, alpha/beta hydrolase from *Pedobacter* sp. V48; SFL10979, alpha/beta hydrolase family protein from *Porphyromonadaceae* bacterium KH3CP3RA; WP_051292783, alpha/beta hydrolase from *Olivibacter sitiensis*; OJV86631, alpha/beta hydrolase from *Bacteroidia* bacterium 44-10; WP_083425928, alpha/beta hydrolase from *Zhouia amylolytica*; WP_079717120, alpha/beta hydrolase from *Parapedobacter luteus*; SEL43123, Pimeloyl-ACP methyl ester carboxylesterase from *Parapedobacter koreensis*. **Putative new family 7**: EstC55-227, functionally derived LE from sample compost55 (this study); SFA93429, hypothetical protein SAMN05216266_102299 from *Amycolatopsis marina*; WP_077006225, hypothetical protein from *Saccharothrix* sp. ALI-22-I; SCF35544, Alpha/beta hydrolase family from *Micromonospora saelicesensis*; WP_046565934, alpha/beta hydrolase from *Micromonospora* sp. HK10; WP_007459501, alpha/beta hydrolase from *Micromonospora lupini*; WP_043627491, alpha/beta hydrolase from *Nonomuraea candida*. **Putative new family 8**: EstC76-36, functionally derived LE from sample compost76 (this study); WP_014510770, alpha/beta hydrolase from *Thermus thermophilus*; WP_014629089, phospholipase from *Thermus thermophilus*; BAW02319, esterase from *Thermus thermophilus*; SDE77545, hypothetical protein SAMN04488243_1105 from *Thermus arciformis*; WP_008633307, phospholipase from *Thermus parvatiensis*; WP_016328666, alpha/beta hydrolase from *Thermus oshimai*. **Putative new family 9**: EstC76-221, functionally derived LE from sample compost76 (this study); WP_025747209, alpha/beta hydrolase from *Caldicoprobacter*; WP_010233006, alpha/beta hydrolase from *Clostridium arbusti*; WP_058952930 alpha/beta fold hydrolase from *Clostridium tyrobutyricum*; WP_043904933 alpha/beta fold hydrolase from *Parageobacillus genomosp.*; WP_013779878 alpha/beta hydrolase from *Mahella australiensis*; WP_064551781 alpha/beta fold hydrolase from *Parageobacillus thermoglucosidasius*; WP_026973730 alpha/beta fold hydrolase from *Alicyclobacillus contaminans*.

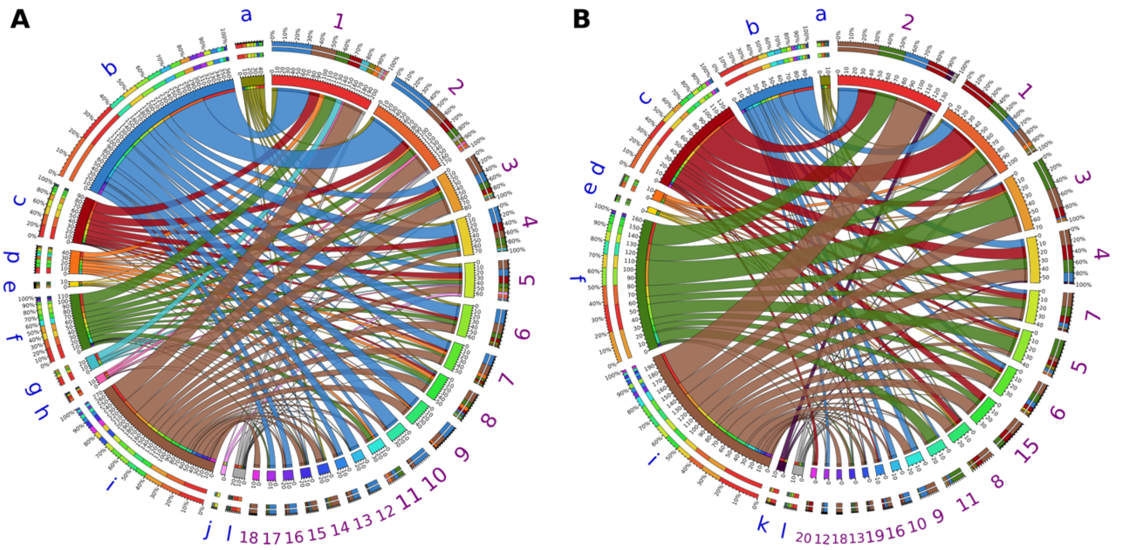


**Supplementary Figure S7.** Phylogenetic distribution at phylum level of assigned PLPs in the most abundant lipolytic families. Assigned PLPs were identified by screening the assembled metagenome of (**A**) compost55 and (**B**) compost76. Taxonomic information of PLP-encoding genes was annotated by KAIJU (Menzel et al. 2016). The data was visualized via Circos software. The width of bars from each phylum (blue) and ESTHER family (purple) indicates their relative abundance. Bacterial phyla: a, *Acidobacteria*; b, *Actinobacteria*; c, *Bacteroidetes*; d, *Chloroflexi* e, *Deinococcus-Thermus*; f, *Firmicutes*; g, *Gemmatimonadetes*; h, *Planctomycetes*; i, *Proteobacteria*; j, *Thermotogae*; k, *Verrucomicrobia*; l, unclassified Bacteria. Lipolytic families in ESTHER databases: 1, VIII ; 2, Hormone-sensitive_lipase_like; 3, patatin-like-protein; 4, II; 5, A85-Feruloyl-Esterase; 6, Carb_B_Bacteria; 7, Homoserine_transacetylase; 8, Lysophospholipase_carboxylesterase; 9, Carboxymethyl-butenolide_lactonase; 10, Polyesterase-lipase-cutinase; 11, CarbLipBact_2; 12, Lipase_2; 13, Chlorophyllase; 14, Tannase; 15, *A85*-EsteraseD-FGH; 16, Fungal_Bact_LIP; 17, Est9X; 18, Bacterial_lip_FamI.3; 19, PC-sterol_acyltransferase; 20, Lipase_3. Only phyla and protein families with a relative abundance >1% are shown.

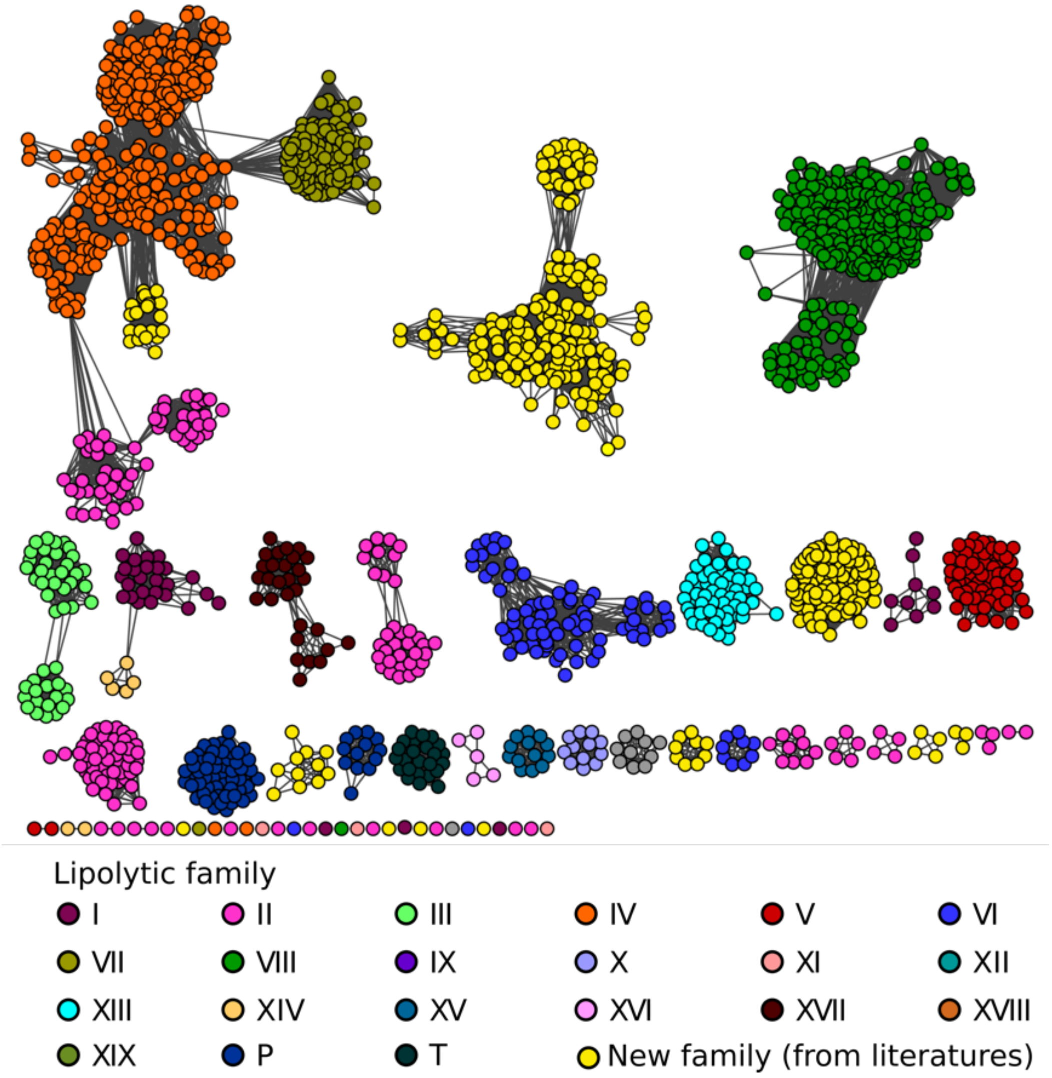


**Supplementary Figure S8**. Protein Sequence similarity network for classification of assigned PLPs obtained by screening against from compost55 and compost 76 assembled metagenomes. Assigned PLPs were pooled and clustered at 100 % identity using CD-HIT (Huang et al. 2010). Then, the resulting sequences were submitted to the EFI-EST (Gerlt et al. 2015) to generate the network. Each node represents an assigned PLP and is colored according to its lipolytic family. Each edge in the network represents a BLAST connection with an E-value cutoff of $\leq 1e^{-15}$. At this cut-off, sequences have a median percent identity and alignment length of 35% and 291 amino acids, respectively. Lengths of edges are not meaningful except that sequences in tightly clustered groups are relatively more similar to each other than sequences with few connections. Nodes were arranged using the *yFiles* organic layout provided with Cytoscape version 3.4.0 (Shannon et al. 2003).

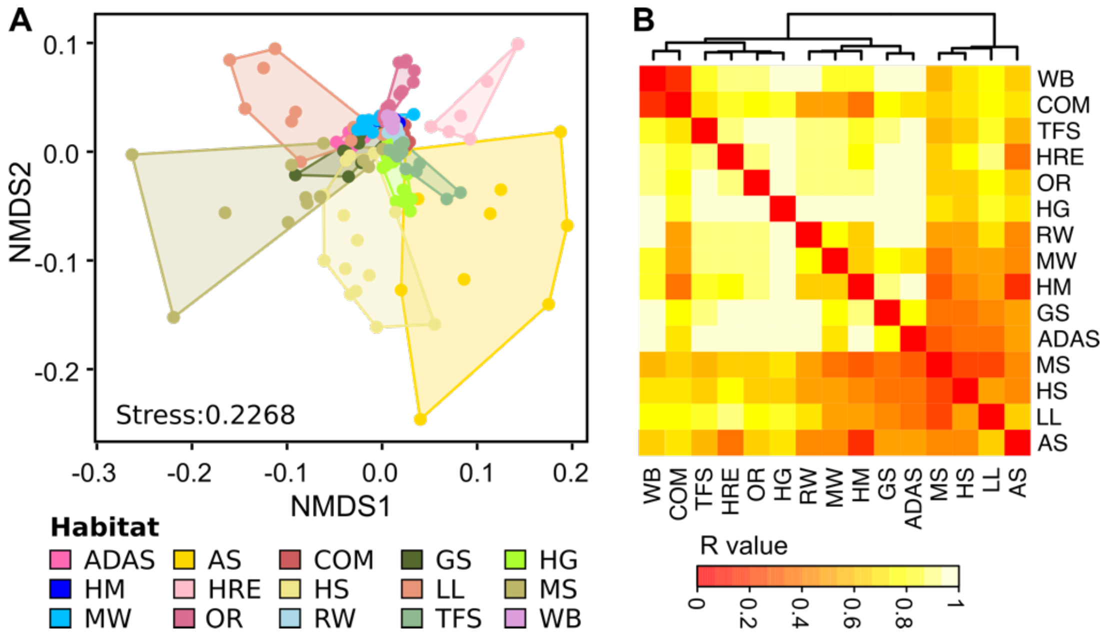


**Supplementary Figure S9.** Functional lipolytic family profiles of assigned PLPs in different samples. A, Non-metric multidimensional scaling (NMDS) analysis of the lipolytic family profiles across samples was performed. Samples were colored by its habitat source. B, ANOSIM test the group dissimilarity of lipolytic family profiles between habitats (9999 permutations, p <0.001). The resulting R values are shown by the heatmap, and the color intensity (red to light yellow) indicates the change of R values (0 to 1). Hierarchical clustering analysis of R values was performed to generate the cluster dendrogram using the Ward.D clustering method based on Bray-Curtis distance matrices. For all the analysis, LPGM values were $\log_{10}$ transformed. Abbreviations of habitats: ADAS, anaerobic digestor active sludge; AS, agricultural soil; COM, compost; GS grassland soil; HG, human gut; HM, hypersaline mat; HRE, hydrocarbon resource environment; HS, hot spring; LL, landfill leachate; MS, marine sediment; MW, marine water; OR, oil reservoir; RW, river water; TFS, tropical forest soil; WB, wastewater bioreactor.

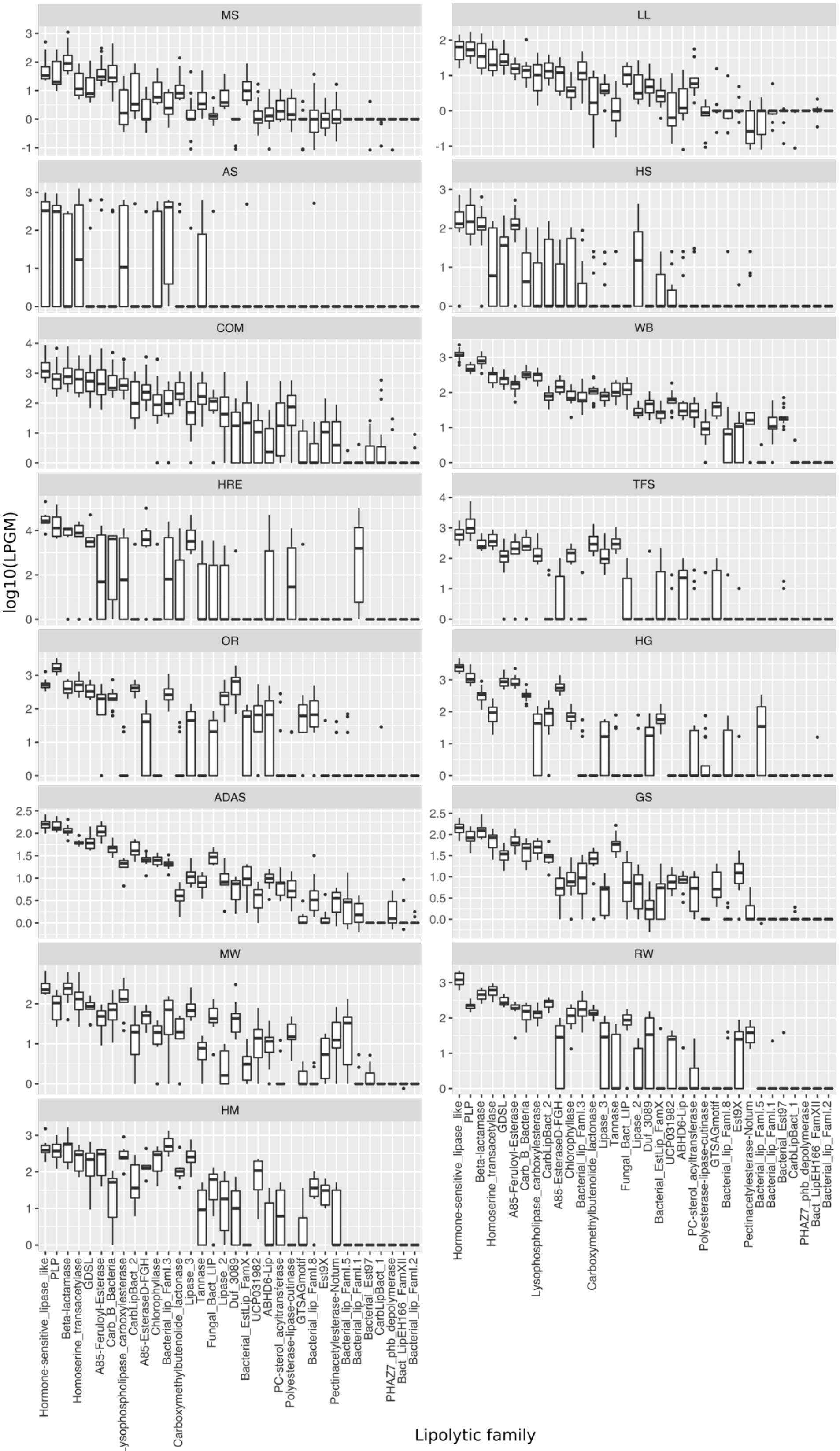


**Supplementary Figure S10.** Distribution of lipolytic families revealed from assigned PLPs of each habitat. The abundance was inferred from $log_{10}$ scaled LPGM values.

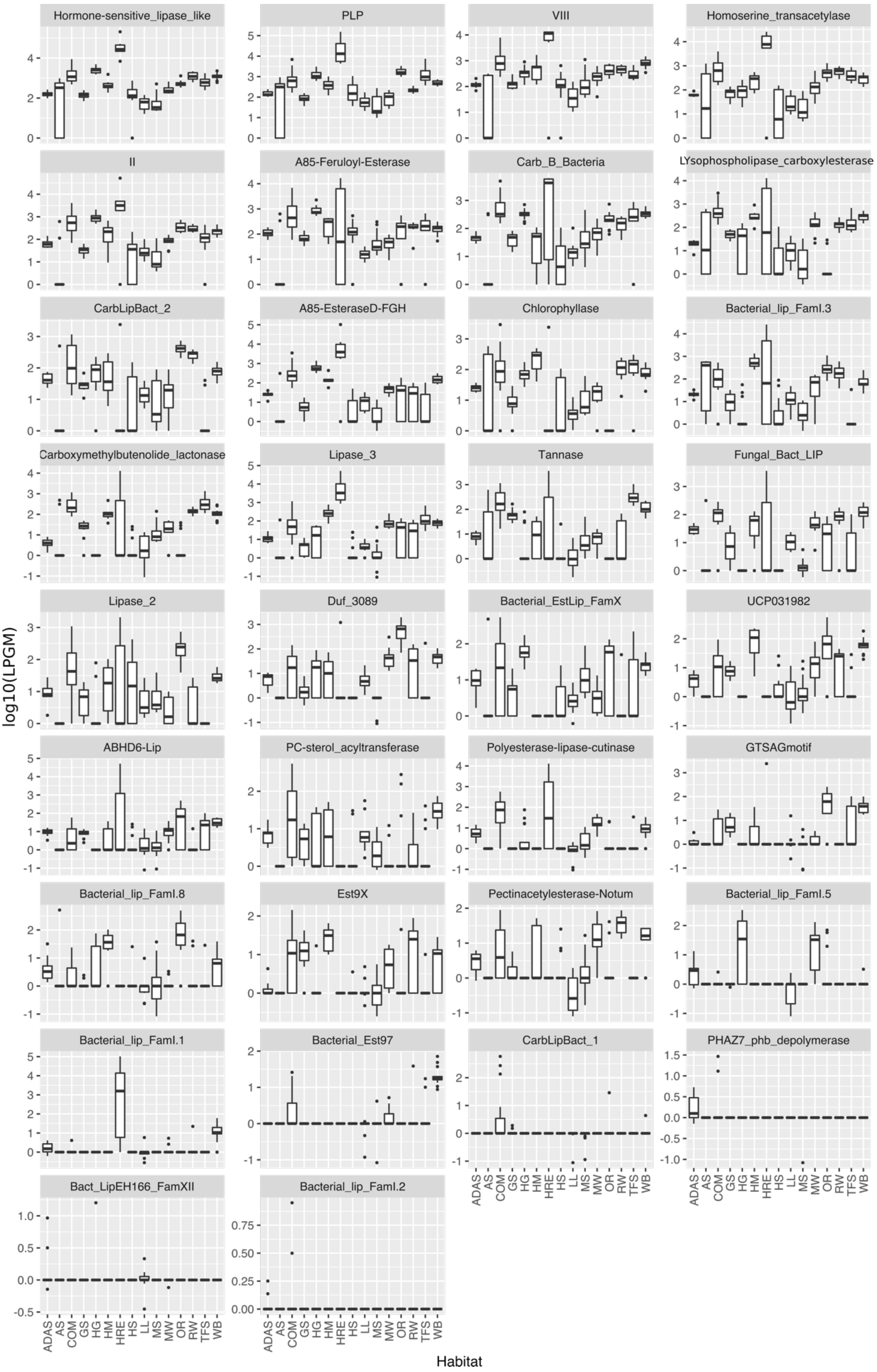


**Supplementary Figure S11.** Lipolytic families showing significant changes in abundance across different habitats. The abundance was inferred from $\log_{10}$ scaled LPGM values.

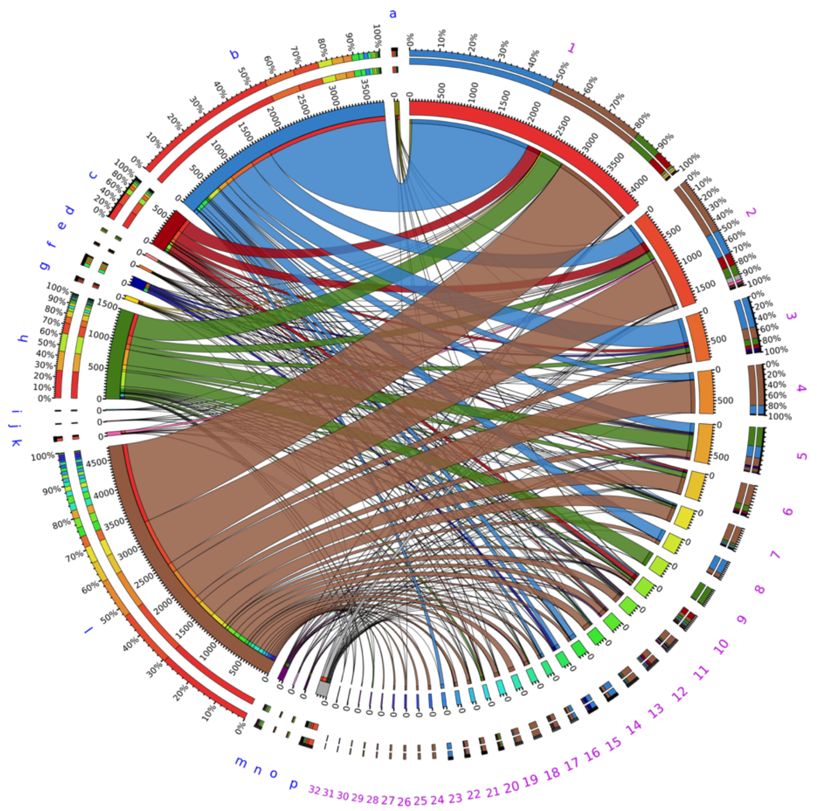


**Supplementary Figure S12.** Phylogenetic origins of LEs in ESTHER database at phylum level in the most abundant lipolytic families. Taxonomic information of lipolytic genes was retrieved from ESTHER database (Lenfant et al. 2013). Visualization was performed via Circos software. The width of bars from each phylum (blue) and lipolytic family (purple) indicates their relative abundance. Bacterial phyla: a, *Acidobacteria*; b, *Actinobacteria*; c, *Bacteroidetes*; d, Chlorobi; e, *Chloroflexi*; f, Cyanobacteria; g, *Deinococcus-Thermus*; h, *Firmicutes*; i, *Gemmatimonadetes*; j, *Lentisphaerae*; k, *Planctomycetes*; l, *Proteobacteria*; m, *Spirochaetes*; n, *Thermotogae*; o, *Verrucomicrobia*; p, unclassified Bacteria. Lipolytic family based on ESTHER database (Arpingy classification): 1, Carb_B_Bacteria (VII) ; 2, Hormone-sensitive_lipase_like (IV); 3, Chlorophyllase (EstGS); 4, Carboxymethylbutenolide_lactonase (V.2); 5, CarbLipBact_2 (XIII-2); 6, A85-EsteraseD-FGH (lp_3505/FLS12/EstA); 7, Homoserine_transacetylase (Est22); 8, Fungal-Bact_LIP (X-2/XVII/XIX); 9, CarbLipBact_1 (XIII-1/XVIII); 10, A85-Feruloyl-Esterase (Rlip1/EstSt7); 11, LYsophospholipase_carboxylesterase (VI); 12, Duf_3089 (XV); 13, UCP031982 (V.3); 14, Est9X (Est9X); 15, Lipase_2 (I.4/I.7); 16, Polyesterase-lipase-cutinase (III); 17, Pectinacetylesterase-Notum (LipT); 18, Bacterial_lip_FamI.8 (I.8); 19, GTSAGmotif (IV); 20, Bacterial_EstLip_FamX (X); 21, Lipase_3 (XI); 22, ABHD6-Lip (V.1); 23, Bacterial_Est97 (XVI); 24, Bacterial_lip_FamI.1 (I.1); 25, PHAZ7_phb_depolymerase (IX); 26, Tannase (Tannase); 27, Bacterial_lip_FamI.3 (I.3); 28, Bacterial_lip_FamI.2 (I.2); 29, Bacterial_lip_FamI.5 (I.5); 30, Bacterial_lip_FamI.6 (I.6); 31, PC-sterol_acyltransferase (XIV); 32, Bact_LipEH166_FamXII (XII). Only phyla and ESTHER families with a relative abundance >1% are shown..

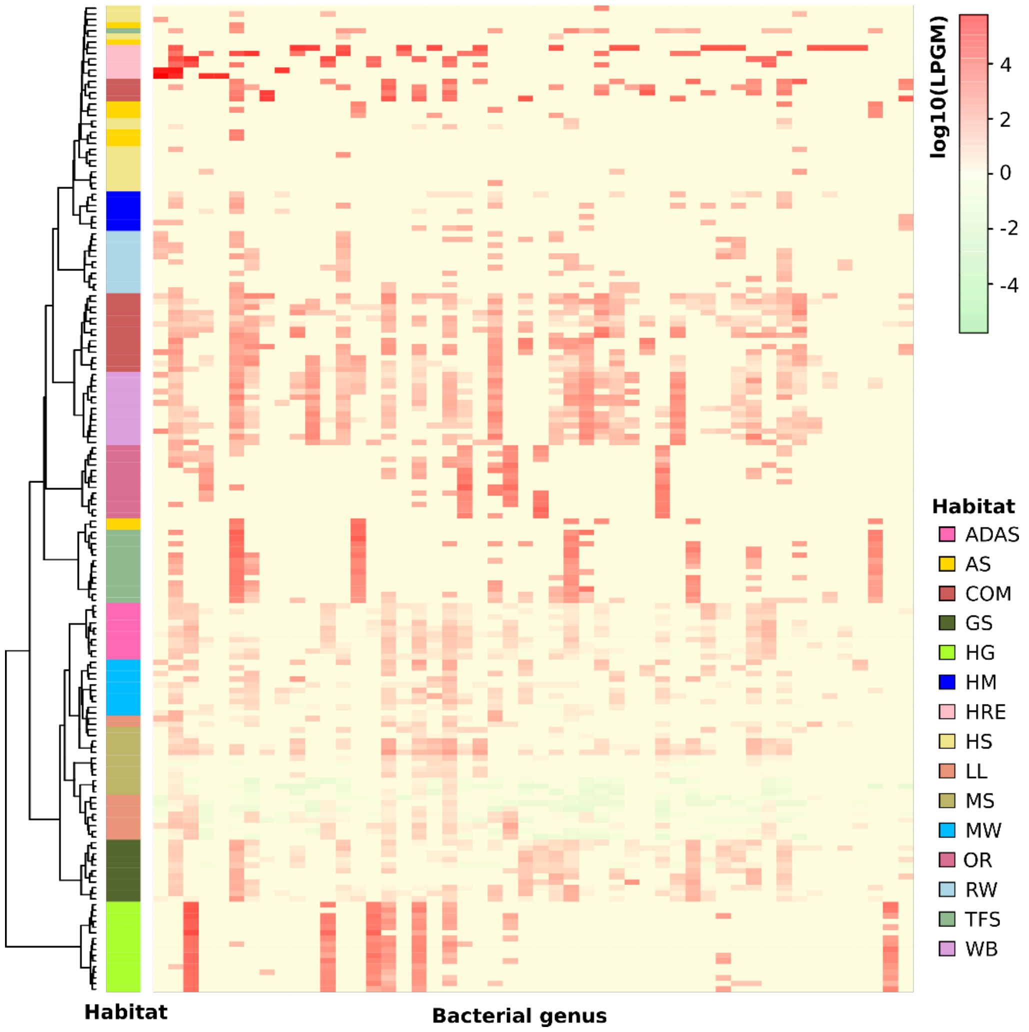


**Supplementary Figure S13.** Taxonomic origins at genus level of the assigned PLPs across samples. The abundance of assigned PLPs per each genus in each sample was inferred from LPGM values. Only genera with a mean LPGM value of ≥0.5 across all the samples were used for analysis, only the top 50 genera areshown here (ranked by the mean LPGM values across samples). Hierarchical clustering analysis of the phylogenetic distribution profile in each sample was performed using the Ward.D clustering method based on Bray-Curtis distance matrices. The color intensity of the heat map (light green to red) indicates the change of abundance (low to high). The habitats were presented by different colors. The phylogenetic distribution of assigned PLPs in each sample was generally clustered by habitats (overall R value = 0.8199, P <0.001, ANOSIM test). Abbreviations: ADAS, anaerobic digestor active sludge; AS, agriculture soil; COM, compost; GS grassland soil; HG, human gut; HM, hypersaline mat; HRE, hydrocarbon resource environments; HS, hot spring; LL, landfill leachate; MS, marine sediment; MW, marine water; OR, oil reservoir; RW, river water; TFS, tropical forest soil; WB, wastewater bioreactor.

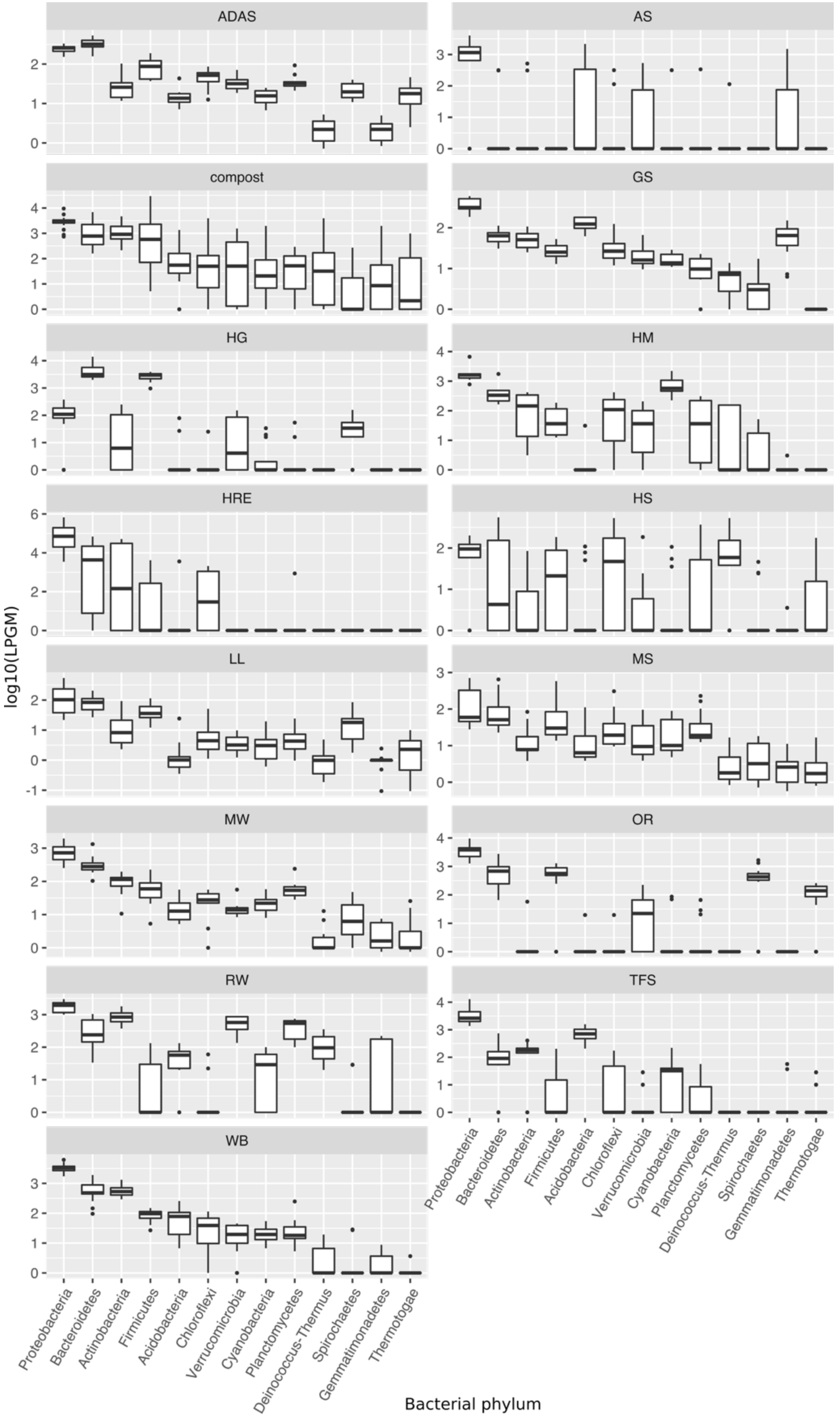


**Supplementary Figure S14.** Phylogenetic distribution of the assigned PLPs at phylum level in each habitat. The abundance was inferred from $\log_{10}$ scaled LPGM values.

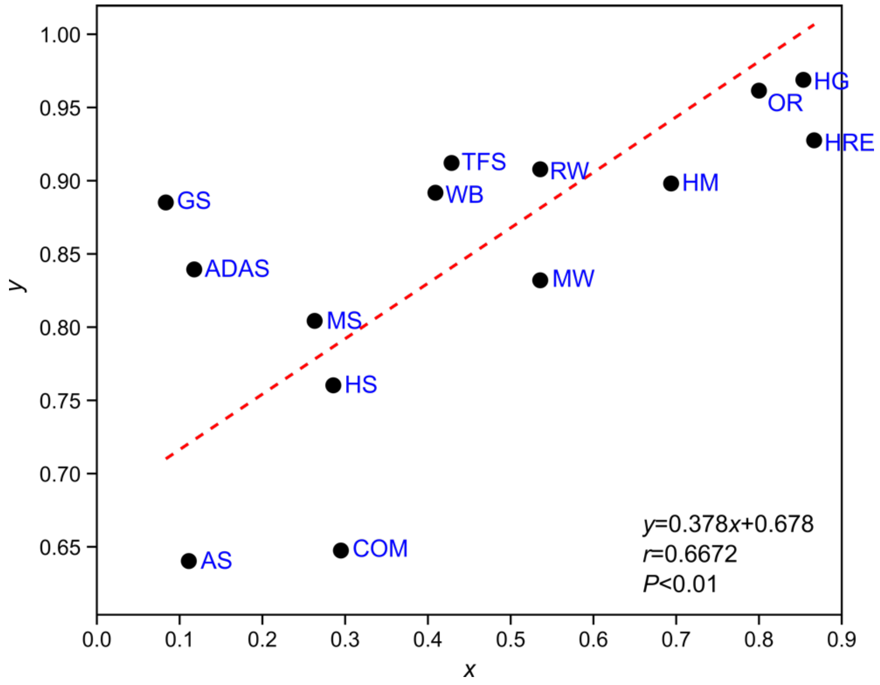


**Supplementary Figure S15.** Linear regression. The x was the ratio of unique indicators to the total significant indicators in a habitat, as demonstrated by the bipartite association network shown in Figure 5. The corresponding y was the mean dissimilarity of the taxonomic profile of assigned PLPs across habitats, in terms of averaged R values generated by the ANOSIM test (P <0.001; see Supplementary Table S18). The Reduced Major Axis (RMA) algorithm was used for regression. The permutation test on correlation uses 9,999 replicates. Pearson's r correlation r=0.6672, p<0.01.

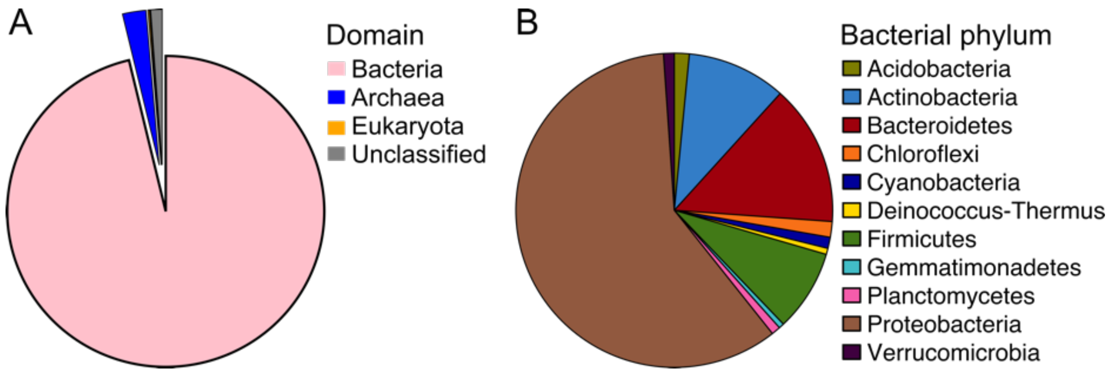


**Supplementary Figure S16.** Phylogenetic origin of the total PLPs (assigned and unassigned PLPs combined) at (A) domain and (B) phylum level. The abundance of PLPs in each domain or phylum was calculated by summing the LPGMs values across samples.

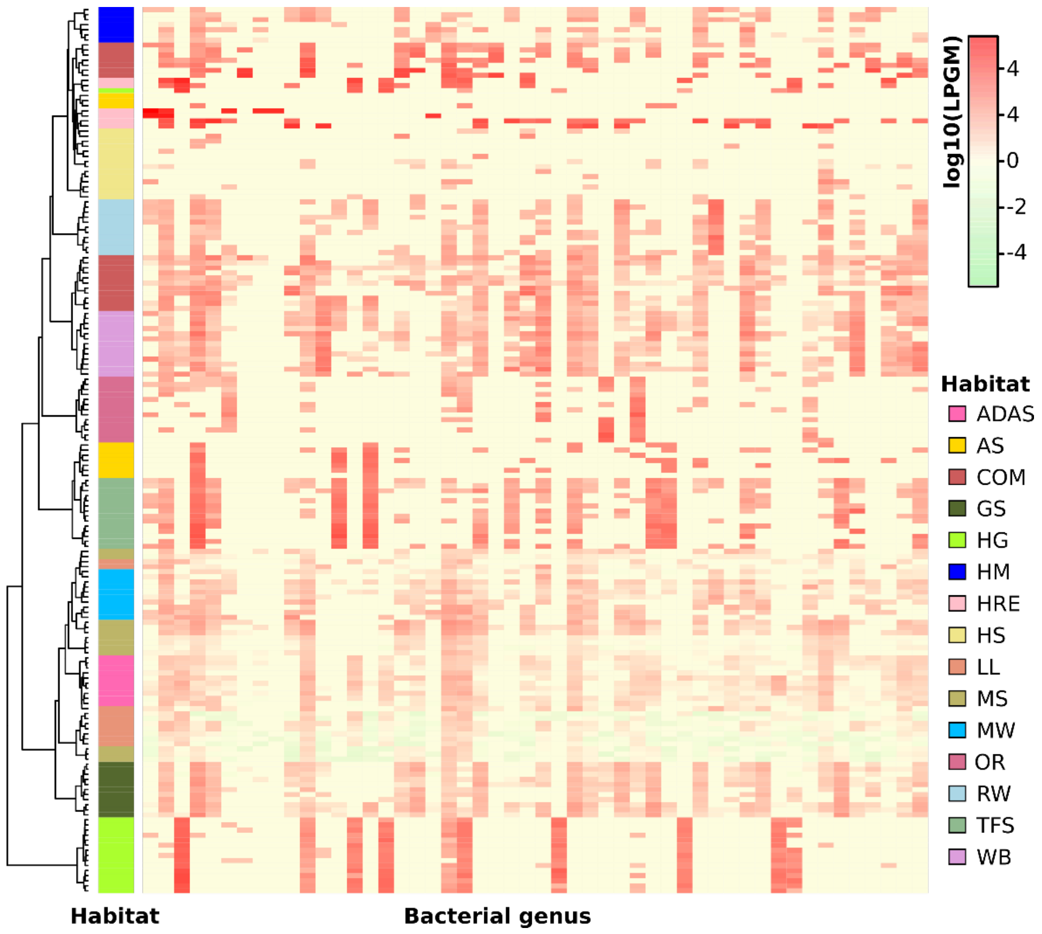


**Supplementary Figure S17.** Heat map of the taxonomic origins at genus level of total PLPs across samples. The abundance of total PLPs per genus in each sample was inferred from LPGM values. Only genera with a mean LPGM value of ≥0.5 across all samples were used for analysis and only the top 50 genera are shown here (ranked by the mean LPGM values across samples). The clustering analysis was performed using the Ward.D clustering method based on Bray-Curtis distance matrices. The color intensity of the heat map (light green to red) indicates the change of abundance (low to high). The habitats are presented by different colors. The phylogenetic distribution of assigned PLPs in each sample clustered generally by habitats (overall R value = 0.821, P <0.001, ANOSIM test). Abbreviations: ADAS, anaerobic digestor active sludge; AS, agriculture soil; COM, compost; GS grassland soil; HG, human gut; HM, hypersaline mat; HRE, hydrocarbon resource environments; HS, hot spring; LL, landfill leachate; MS, marine sediment; MW, marine water; OR, oil reservoir; RW, river water; TFS, tropical forest soil; WB, wastewater bioreactor.

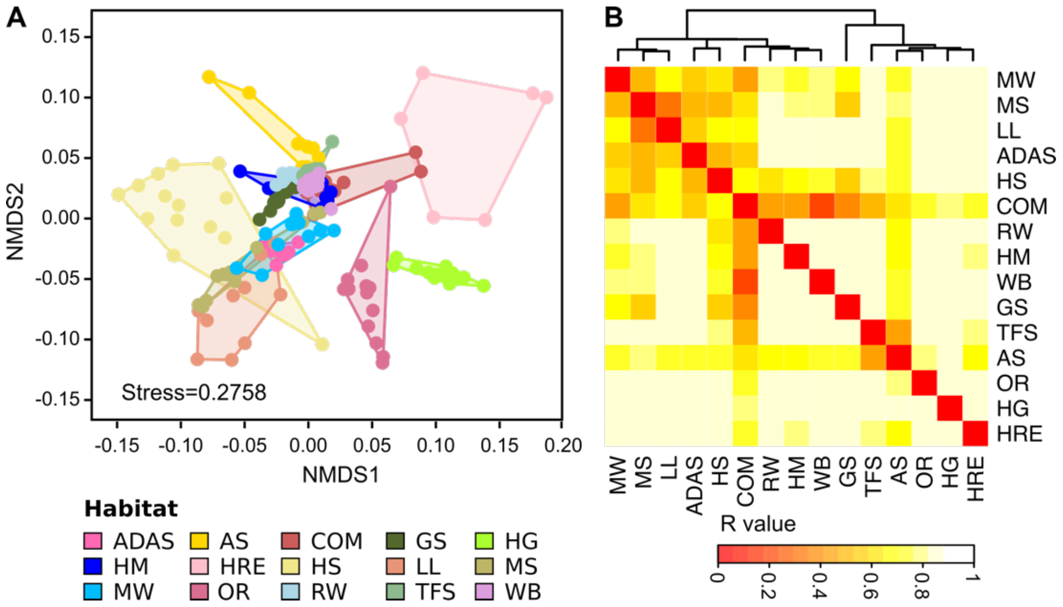


**Supplementary Figure S18.** Analysis of the phylogenetic profile at genus level of total PLPs across samples. A, Non-metric multidimensional scaling (NMDS) analysis of the phylogenetic profile of total PLPs across samples was performed. Samples were colored by its habitat origin. The abundance of PLPs in each genus per sample is presented by the LPGM values. Only genera with mean LPGM values of ≥0.5 across all the samples were used for analysis. B, ANOSIM test the group dissimilarity of these phylogenetic profiles between habitats (9999 putations, P <0.001). The resulting R values are shown by the heatmap, and the color intensity (red to light yellow) indicates the change of R values (0 to 1). Hierarchical clustering analysis of R values was performed to generate the cluster dendrogram using the Ward.D clustering method based on Bray-Curtis distance matrices

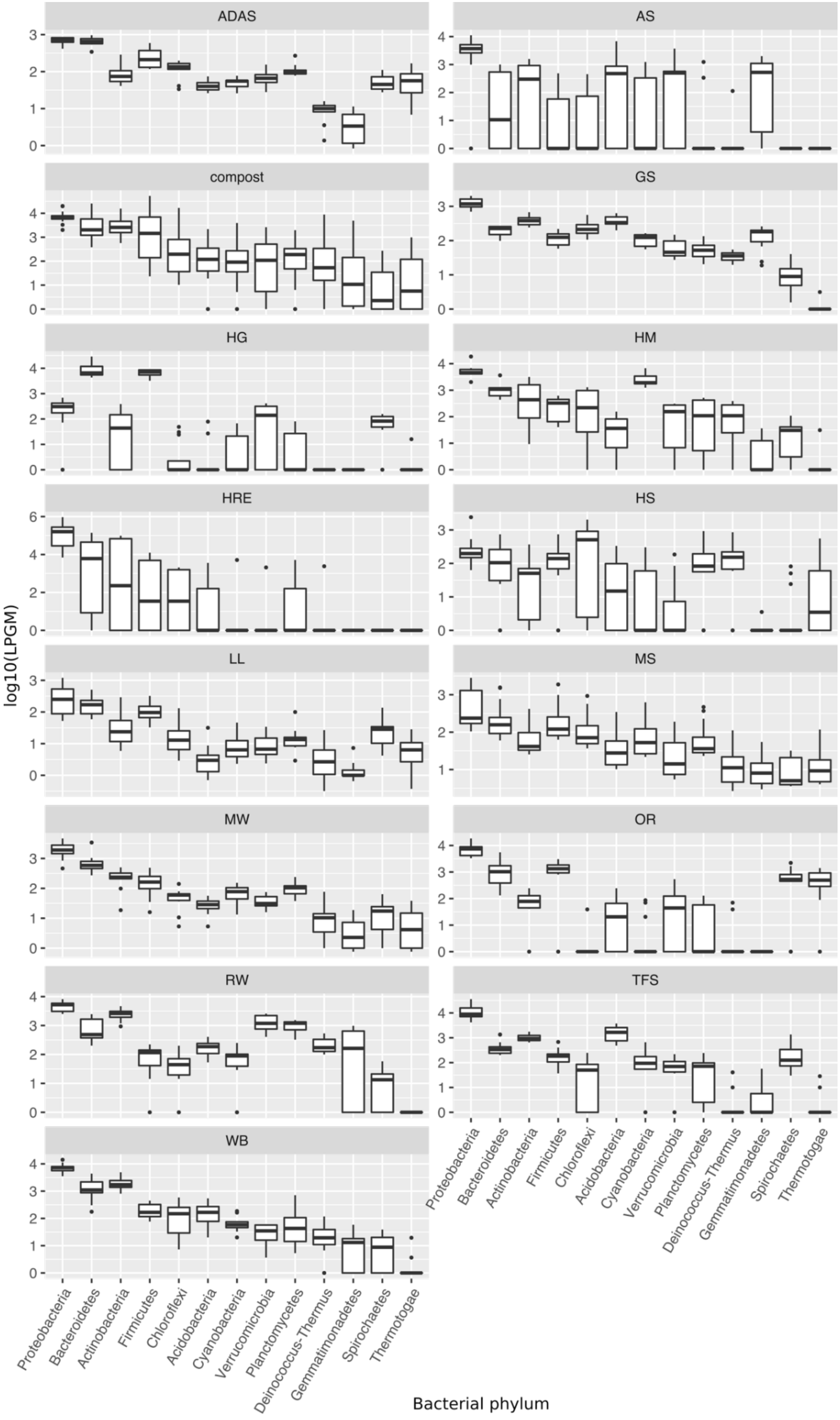


**Supplementary Figure S19.** Phylogenetic distribution of the total PLPs at phylum level of each habitat. The abundance was inferred from $log_{10}$ scaled LPGM values.
